# A dual hit of α-synuclein internalization and immune challenge leads to the formation and maintenance of Lewy body-like inclusions in human dopaminergic neurons

**DOI:** 10.1101/2023.05.29.542776

**Authors:** Armin Bayati, Riham Ayoubi, Adriana Aguila, Cornelia E. Zorca, Chanshuai Han, Emily Banks, Emmanuelle Nguyen-Renou, Wen Luo, Irina Shlaifer, Esther Del Cid-Pellitero, Moein Yaqubi, Edward A. Fon, Jo Anne Stratton, Thomas M. Durcan, Patrick C. Nahirney, Peter S. McPherson

**Author notes:** Corresponding authors: Armin Bayati, Dr. Peter S. McPherson, Department of Neurology and Neurosurgery, Montreal Neurological Institute McGill University, 3801 University Street Montreal, QC H3A 2B4 Canada. Authors contributed equally: Second co-authors (Riham Ayoubi, Adriana Aguila).

## Abstract

Lewy bodies (LBs), rich in α-synuclein, are a hallmark of Parkinson’s disease (PD). Understanding their biogenesis is likely to provide insight into the pathophysiology of PD, yet a cellular model for LB formation remains elusive. The realization that the immune challenge is a trigger for neurodegenerative diseases has been a breakthrough in the understanding of PD. Here, iPSC-derived human dopaminergic (DA) neurons from multiple healthy donors were found to form LB-like inclusions following treatment with α- synuclein preformed fibrils, but only when coupled to an immune challenge (interferon-gamma or interleukin-1 beta) or when co-cultured with activated microglia. Human cortical neurons derived from the same iPSC lines did not form LB-like inclusions. Exposure to interferon-gamma impairs autophagy in a lysosomal-specific manner *in vitro,* similar to the disruption of proteostasis pathways that contribute to PD. We find that lysosomal membrane proteins LAMP1 and LAMP2 and transcription factors regulating lysosomal biogenesis and function are downregulated in DA but not cortical neurons. Finally, due to the excellent sample preservation afforded by cells compared to post-mortem PD brain tissue, we conclude that the LB-like inclusions in DA neurons are membrane-bound, suggesting they are not limited to the cytoplasmic compartment.

**In Brief:** 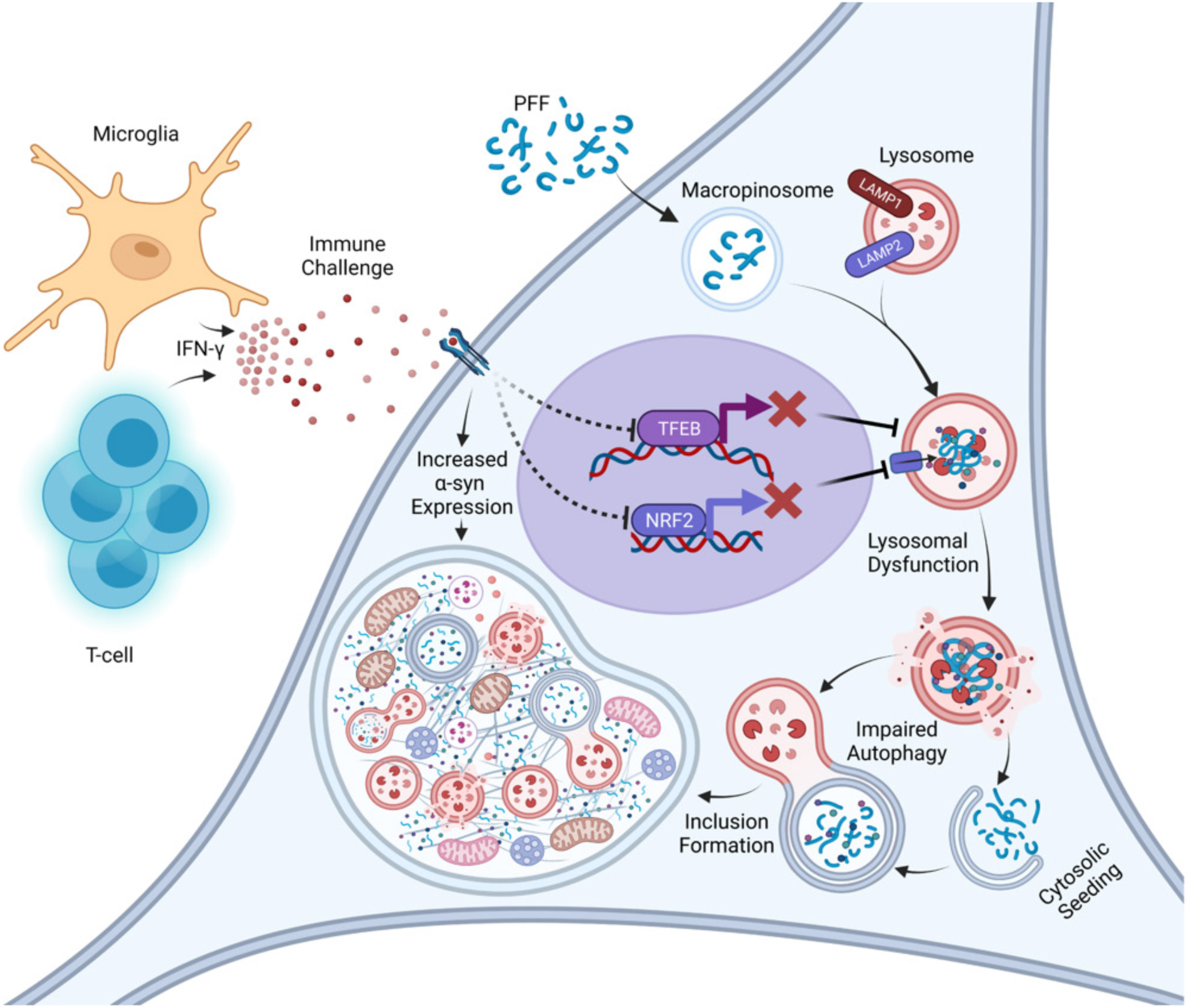

Bayati et al. identify that iPSC-derived dopaminergic neurons undergoing a dual hit treatment of exogenous α-synuclein fibrils and proinflammatory cytokines form Lewy body-like inclusions. The dual hit treatment also led to the downregulation of lysosomal proteins. Characterization of inclusions revealed that inclusions were membrane-bound and LC3B-positive, suggesting they are dysfunctional autophagosomes.

**Highlights:**

- α-synuclein preformed fibril administration coupled with Interferon-gamma exposure leads dopaminergic neurons to form Lewy body-like inclusions
- Inclusions are filamentous, membranous, and filled with aberrant organelles
- Impaired autophagic flux and downregulation of TFEB, NRF2, LAMP1, and LAMP2 correlated with inclusion formation
- Activation of NRF2 through the treatment of neurons with the antioxidant perillaldehyde, prevents inclusion formation

## INTRODUCTION

Lewy bodies (LBs) are proteinaceous inclusions that are a prevalent pathological marker in Parkinson’s disease (PD) ^1,2^. Their presence in patients can only be confirmed using post-mortem tissue. LBs are present in most patients diagnosed with idiopathic/sporadic PD ^3^, which comprises almost 80-90% of cases ^4^. LBs also appear in many of the familial forms of PD. Coupled with the appearance of LBs, dysfunctional neurons also exhibit granular and filamentous deposits, referred to as Lewy neurites (LNs) ^5^. Despite their incidence in PD and other synucleinopathies such as Lewy body dementia (LBD) ^6,7^, and their histological discovery in the early 20^th^ century ^8^, insights into the formation, structure, and function of LBs remain limited. This is primarily due to reliance on post-mortem tissue, in which LB morphology and content may be affected by alterations in tissue pH, oxygen levels, and autolytic activity during the post-mortem interval ^9^. Using superior approaches to collecting and processing post-mortem tissue, Shahmoradian et al. ^10^ demonstrated that LBs are not exclusively filamentous aggregates but are composed of a collection of membrane-bound compartments and organelle fragments. Their data, corroborating other previous studies ^11-13^, has spurred renewed interest in the structural study of LBs, the role that dysfunctional organelles play in the formation of LBs, and the underlying causes leading to the formation of membranous, proteinaceous, and organelle-filled inclusions.

While LBs may be a symptom of cellular dysfunction, they might also play a role in PD pathophysiology, neurotoxicity, or neuroprotection^6^. LBs have been hypothesized to be an aggresomal response to the accumulation of aggregated proteins ^14^, which is perhaps a symptom of dysfunction in the degradative/autophagic machinery of the cell, a pattern also observed in lysosomal storage diseases ^15^. PD bears many similarities with lysosomal storage diseases ^16^, and PD has been linked to impaired autophagy ^17^. Moreover, mutations in mitochondrial proteins PINK1 and parkin result in dysfunctional mitophagy ^18^. It is thus plausible that deficits in autophagy found in PD are responsible for LB formation ^19^.

Consistent with protein degradation defects in LB formation, lysosomal proteins including GBA and TMEM175, are implicated in PD pathophysiology ^18,20^. LAMP1 and LAMP2 are major lysosomal proteins that maintain lysosome integrity, pH, catabolism, and both function in autophagy ^21,22^. Interestingly, there are reduced levels of both proteins in PD ^19^. Such perturbations to the protein degradation machinery are not only common in PD, but a combination of such disturbances to lysosomal activity may play a crucial role in PD pathophysiology.

There is growing interest in the role of microglia and the immune system in neurodegenerative diseases. A seminal paper by Matheoud et al. ^23^ reveals that PINK1^-/-^ mice only exhibit PD-like symptoms when introduced to infections, emphasizing the importance of an immune-related trigger in the manifestation of PD motor symptoms. Further, the authors found cytotoxic T-cells, trained on recognizing mitochondrial antigen-presenting cells, targeted PINK1-knockout dopaminergic neurons in culture. The role of an immune challenge in PD has been highlighted by other studies, focusing on LRRK2 and cytokines ^24,25^. In a mouse model of amyotrophic lateral sclerosis, the same line housed at two different facilities had different phenotypic outcomes, with more severe motor symptoms in animals at a facility where they had measurably more exposure to bacterial pathogens ^26^. Clearly, the immune system and cytokines play a role in neurodegenerative diseases ^27-29^.

Given the link between the immune system and neurodegenerative disease, we hypothesized that a dual hit of a PD-insult followed by an immune challenge leads to the formation of LBs. We exposed iPSC-derived dopaminergic (DA) neurons to α-synuclein (α-syn) preformed fibrils (PFFs), which are rapidly internalized and transported to lysosomes ^30^. After a delay, we presented the neurons with an acute treatment of IFN-gamma (IFN-γ), a compound shown to be released by microglia and immune cells in response to infections ^31-34^. As Matheoud *et al.* ^23^ demonstrated the targeting of PINK1- knockout DA neurons by T-cells, who are major producers of IFN-γ ^32,35-37^, we decided that IFN-γ would be a good candidate to represent the involvement of the immune system. Exposure to PFF and IFN-γ results in the formation and long-term maintenance of PFF-positive inclusions absent in neurons treated with either substance alone. Like LBs, the LB-like inclusions formed in DA neurons are organellar, membranous and filamentous as determined by light and electron microscopy. LB-like inclusions also formed when interleukin-1 beta (IL-1β), a more microglia-specific proinflammatory cytokine, was used instead of IFN-γ. Remarkably, the resulting LB-like inclusions were membrane-bound, suggesting that they do not form in the cytoplasm but within an organelle lumen, likely in the endo/lysosomal system, or that they originate in the cytoplasm but are eventually engulfed by membranous organelles such as autophagosomes. These inclusions do not form in other cell types, including cortical neurons. Analysis of protein expression in response to the dual hit treatment reveals DA neuron-specific downregulation of transcription factor EB (TFEB) and nuclear factor erythroid 2-related factor 2 (NRF2), transcription factors involved in lysosomal biogenesis and function, along with the downregulation of lysosomal membrane proteins LAMP1 and LAMP2. These data provide a unique cellular model for LB formation.

## RESULTS

### DA neurons form LB-like inclusions following a dual hit of PFF internalization and immune challenge

α-syn PFFs, a potentially toxic form of the protein, are internalized into a variety of cell types by macropinocytosis and are rapidly transported to lysosomes and multivesicular bodies (MVBs) where they remain for several days ^30^. Given the growing link between immune challenges and neurodegenerative diseases, we sought to examine if an immune challenge could alter the cellular fate of internalized PFFs. IFN-γ, a cytokine released by immune cells and microglia in response to infections ^31^, affects autophagy and inhibits lysosomal activity by downregulating lysosomal membrane proteins LAMP1 and LAMP2^38^. Our treatment regime, described in Fig. 1A and in S1A and S1B, involves the addition of sonicated PFFs (Fig. S1C) and/or IFN-γ to DA neurons derived from human iPSCs (Fig. S1D), which express IFN-γ receptor 1 and 2 (IFNGR1 and IFNGR2; Fig. S2A). Remarkably, we observe the formation of PFF-positive inclusions using both light and electron microscopy analysis but only when the PFF treatment is combined with 0.2 mg/mL of IFN-γ (Fig. 1B, 1C, and S2B). PFFs pre-labeled with nanogold are seen in dark lysosomes and autolysosomes inside the inclusions, which also contain mitochondria and filamentous materials, all characteristic of LBs ^10,39^. These LB-like inclusionary bodies are ∼5-10 µm, located in the perinuclear region, and in most cases, there is only one of these bodies per neuron. Approximately 15-20 % of neurons form LB-like inclusions. Some neurons also form inclusions within neurites (Fig. 1D), similar to Lewy neurites in PD. Formation of LB-like inclusions was achieved in DA neurons generated from three different iPSC cell lines (Fig. S2C and S2D). Fig. S3 shows a collection of additional inclusions found in DA neurons from each iPSC cell line.

**Figure 1.**
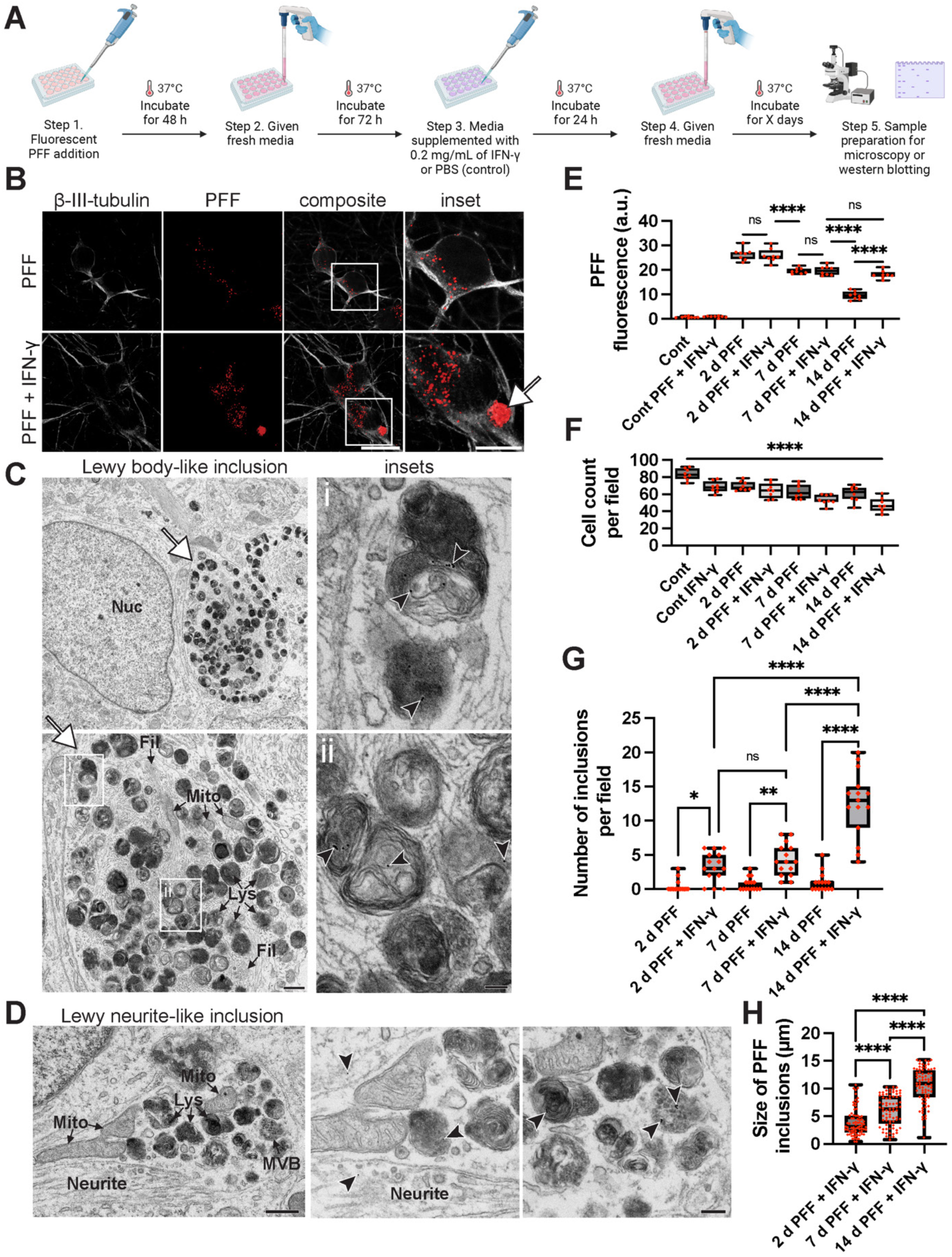
IFN-γ treated DA neurons form PFF-positive inclusions. **(A)** Graphical representation indicates the steps involved in the dual hit regime. iPSC-derived DA neurons underwent a 14 d dual hit treatment, in which fluorescent PFFs were administered for 48 h, incubated in fresh media for 72 h, followed by the administration of IFN-γ (or PBS for control) for 24 h. Neurons were then incubated for 8 d, in fresh media. **(B)** Samples were fixed, stained with β-III-tubulin antibody, and prepared for confocal microscopy. Confocal images indicate the formation of PFF-positive inclusions in PFF + IFN-γ-treated neurons. These inclusions are not present in neurons only treated with PFF at 14 d. Arrowhead points to the PFF-positive inclusion. Scale bar = 20 µm and 10 µm for inset. (**C**) Following the same protocol as in **A**, but using nanogold-labeled PFF, neurons underwent the dual hit treatment regime, were fixed, and processed for EM. Lys (lysosome), Mito (mitochondria), Nuc (nucleus), Fil (filaments). Arrowhead points to the inclusion spotted in the neuron. Arrows point to nanogold-labeled PFF. Scale bar = 500 nm and 100 nm for insets. (**D**) Neurons undergoing the same treatment described in **A** showed inclusions within neurites. MVB (multivesicular bodies). Scale bar = 500 nm and 200 nm for insets. (**E**) Quantification of PFF fluorescence over 14 d in neurons both treated and not treated with IFN-γ, n = 8 for each condition, and quantification was done on data collected from three independent samples. One-way ANOVA and *post-hoc* Tukey’s test were conducted to ascertain the significance between means. (**F**) Cell count was quantified over 14 days, n = 8 for each condition, and quantification was done on three independent samples. The control (Cont) and IFN-only conditions were fixed and counted simultaneously to the 14 d samples; this allows our control conditions to serve as a better control for our main experimental conditions: 14 d PFF and 14 d PFF + IFN. Conditions were analyzed using one-way ANOVA. (**G**) The number of inclusions per 387.5 x 387.5 µm images was counted. PFF puncta of 2 µm and larger were counted as inclusions, n = 15 for each condition, from three independent experiments. One-way ANOVA and *post hoc* Tukey’s test were conducted for statistical analysis. (**H**) The size/diameter of inclusions was calculated, n = 75 for each condition, and data was collected from three independent experiments, with ANOVA and *post hoc* Tukey’s test for statistical analysis. **** denotes p < 0.0001, *** denotes p < 0.001, and ** denotes p < 0.01. ns = not significant.

In the treatment regime, we added PFF for 48 h, and then waited 3 days to add IFN-γ. This maximized cell survival compared to when we added PFF and IFN-γ simultaneously. PFF fluorescence and cell counts were quantified over 14 d (Fig. 1E and F). Neurons treated with PFFs alone exhibited significantly less PFF fluorescence than those treated with PFFs and IFN-γ, suggesting disruption of lysosomal degradation caused by IFN-γ treatment. A general decrease in cell count was observed with the treatment of PFF and IFN-γ. There was also a significant increase in the number and size of inclusions with increased incubation time (Fig. 1G and H).

Lastly, to ascertain whether the formation of LBs using the dual hit treatment regime was IFN-γ specific, we conducted the same experiments but replaced IFN-γ with 50ng/mL of IL-1β. We found that DA neurons treated with PFF and then IL-1β also formed PFF-positive inclusions (Fig. S4A). Using EM, we found that these inclusions are filled with aberrant lysosomes, dysfunctional mitochondria, and are also membrane bound as seen with the IFN-γ treated samples (Fig. S4B).

### Evaluation of the formation of LB-like inclusions

Following LB-like inclusions in cultured neurons allows for observation of the sequence of events leading to their formation and maturation. The inclusions formed in DA neurons (Fig. 1C, Fig. S2C and D, and Fig. S3) are very similar to those observed by Shahmoradian *et al.* ^10^ in PD brains in that they both contain a medley of organelles, membranous fragments, filaments, lysosomes, autolysosomes, and mitochondria. To explore how they develop, a timeline of inclusion formation was established over 14 d (Fig. 2A and B). At 2 d (Fig. S1B), an accumulation of nanogold PFFs in lysosomes occurs in both PFF-only and the PFF + IFN-γ conditions. In the 6 d samples, where neurons were recently treated with IFN-γ (or vehicle in the PFF-only samples), we see dramatic signs of cellular stress including swollen mitochondria and endoplasmic reticulum, along with dark, aberrant lysosomes. In the 7 d samples, the formation of miniature LB-like inclusions containing aberrant lysosomal structures can be observed. In the 10 d samples, a larger LB-like inclusion has formed, containing not only aberrant lysosomal structures but also mitochondrial fragments and a plethora of filamentous structures.

**Figure 2.**
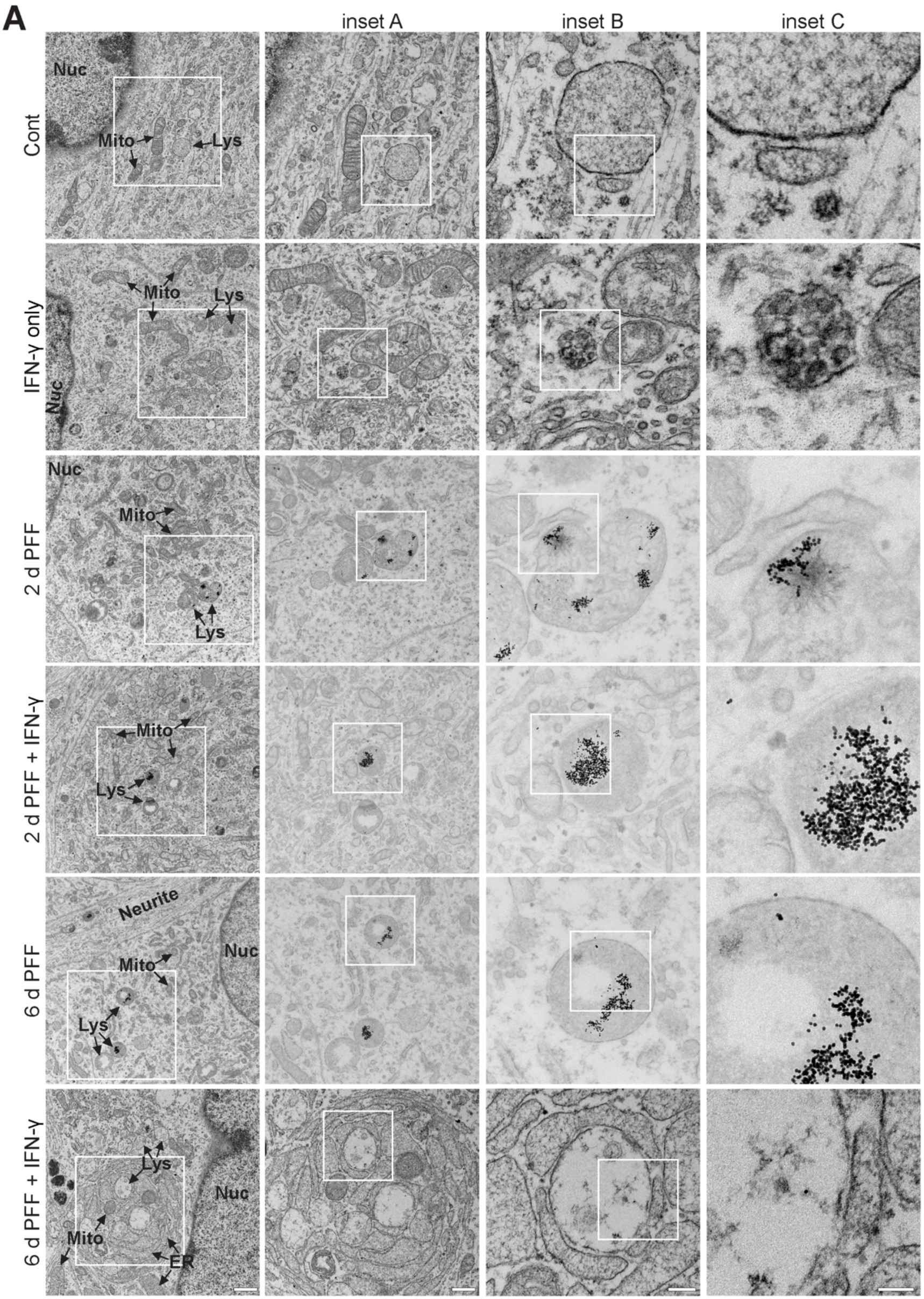
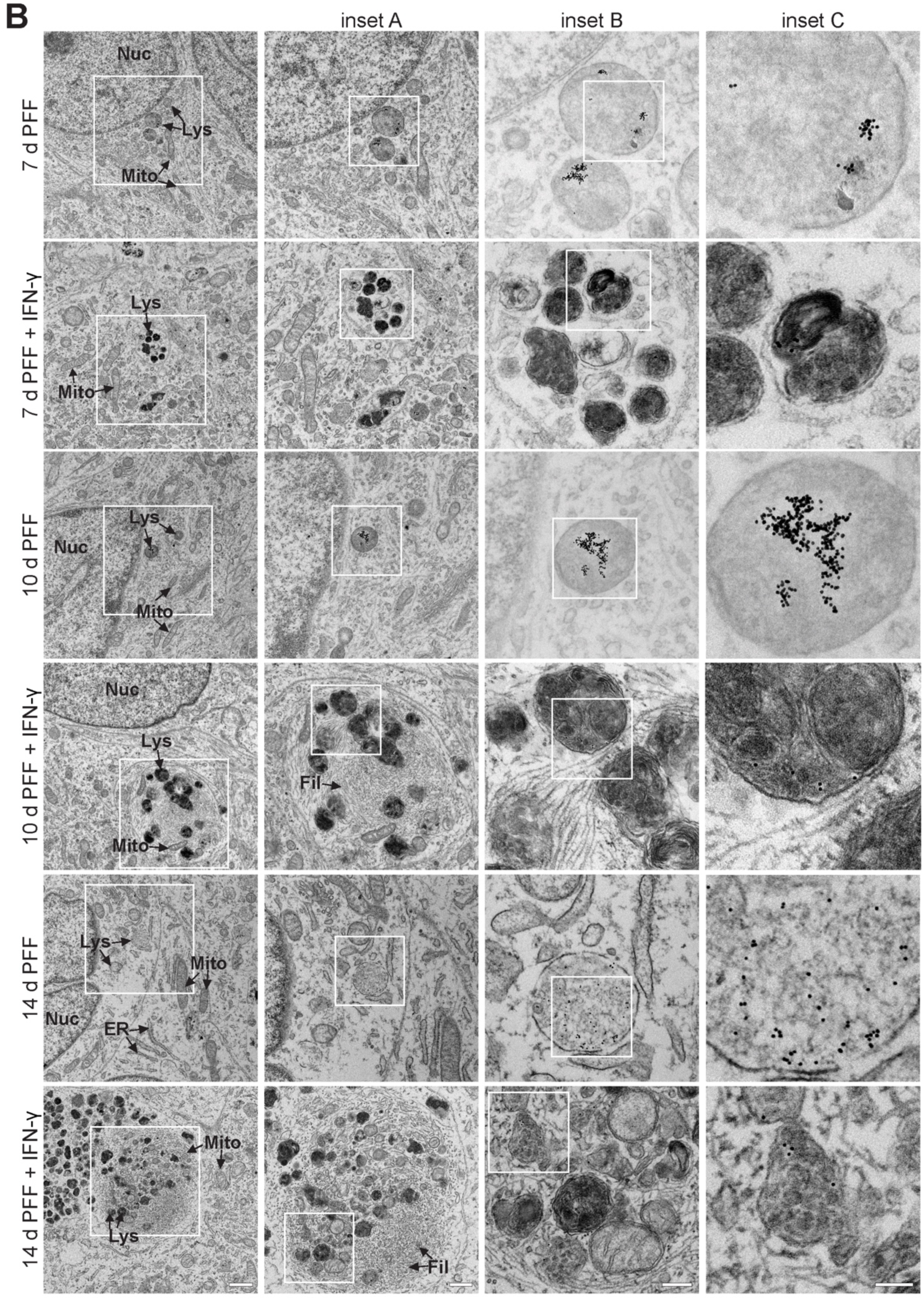
LB-like inclusions form over 14 d. (**A**) DA neurons’ ultrastructure and organelle morphology were analyzed over 14 d, as neurons were put through the dual hit treatment. In the control (Cont) sample, lysosomes are unremarkable and appropriately sized (∼1 µm in diameter), and mitochondria cristae are well defined. In the IFN-only condition, mitochondria are more prominent and swollen, and the cristae are not as perfectly defined as in Cont. Neurons in the 2 d PFF condition exhibit lysosomes that have accumulated nanogold-labeled PFFs and lysosomes that have increased in size compared to the control. In the 2 d PFF + IFN-γ sample, mitochondria have irregular cristae, and an accumulation of nanogold-labeled PFF is seen. 6 d PFF neurons show accumulation of nanogold-PFF in their lumen, with enlarged lysosomes compared to control. 6 d PFF + IFN-γ samples show signs of drastic levels of ER stress, swollen mitochondria, and lysosomes with nanogold-PFFs. (**B**) 7 d PFF samples show lower PFF accumulation levels in lysosomes than 2 d PFF neurons. 7 d PFF and IFN-γ treated neurons exhibit miniature LB-like inclusions with aberrant lysosomal structures within them. 10 d PFF samples show very similar morphology to 7 d PFF neurons. 10 d PFF + IFN-γ-treated neurons exhibit larger LB-like structures (compared to the 7 d PFF + IFN-γ samples) with aberrant lytic structures and abnormal mitochondria with irregular cristae. Accumulation of a dense web of cytoskeletal filaments is evident around some of the dense lysosomes. 14 d PFF neurons look very similar to the control, except that they show nanogold-PFF accumulation. Finally, 14 d PFF and IFN-γ samples show large inclusions containing a collection of filaments, granules, lysosomes, MVBs, and mitochondria. Lys (lysosome, autolysosomes), Mito (mitochondria), ER (endoplasmic reticulum), Nuc (nucleus), and Fil (filaments). Scale bar = 1 µm, 500 nm for inset A, 200 nm for inset B, and 100 nm for inset C.

The most noticeable difference between the PFF-treated neurons and those treated with PFF + IFN-γ is seen at 14 d. The PFF + IFN-γ samples exhibit large inclusions that contain dark lysosomes, MVBs, filaments, and mitochondria. Additional examples can be found in Fig. S2C and D along with Fig. S3A-G. Almost all inclusions formed by the 14 d PFF + IFN-γ treated DA neurons are membrane-bound. We followed the LB-like inclusions for up to 30 days, revealing highly compact, complex, filamentous/organelle-filled, membrane-enclosed inclusions (Fig. 3A-C). Additional examples of inclusions formed in 30 d DA neurons can be found in Fig. S5A-D.

**Figure 3.**
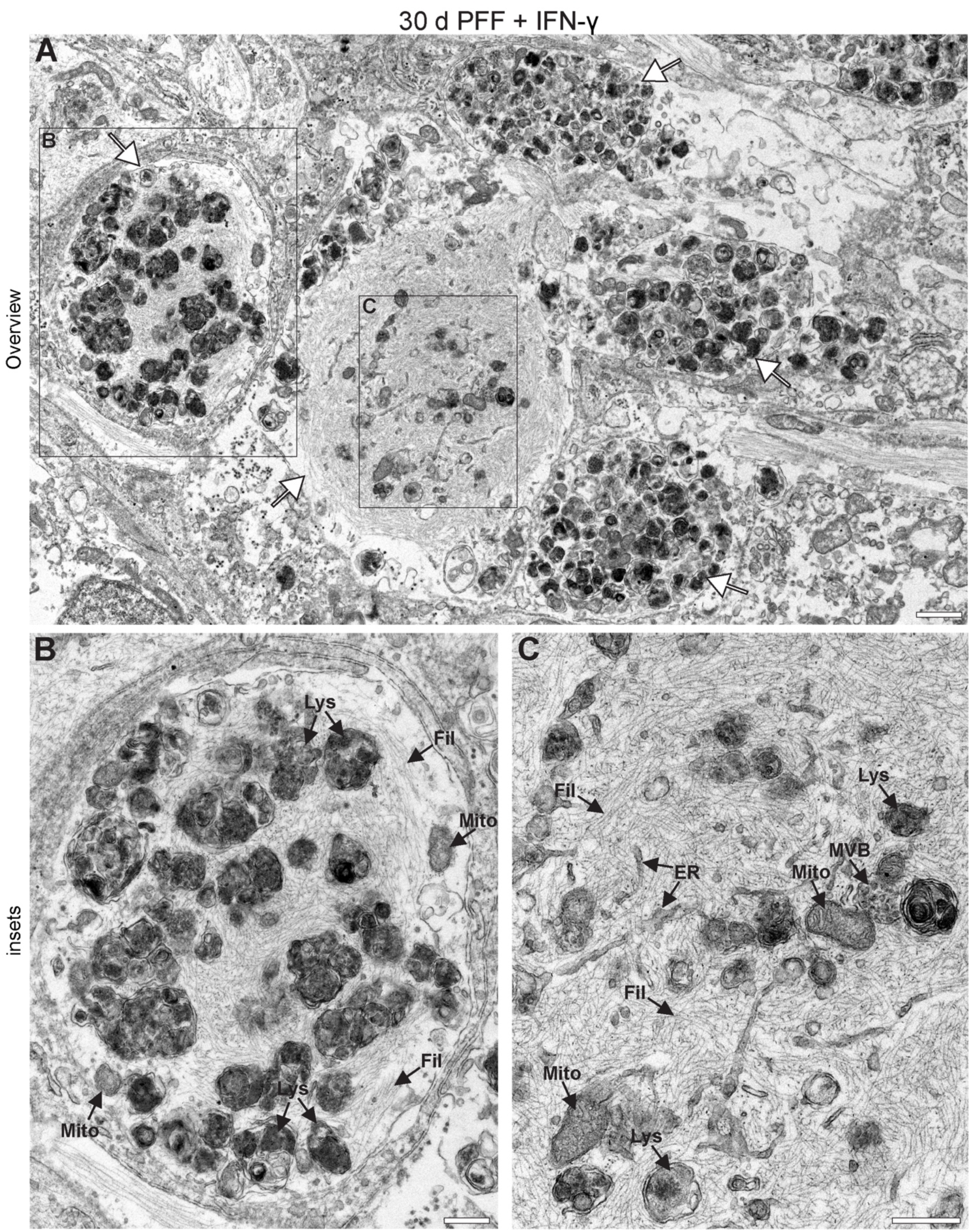
Highly dense, mature inclusions form in DA neurons with increased incubation time. (**A**) DA neurons underwent the 14 d dual hit treatment regime and were then incubated for an additional 16 d, for a total of 30 d in culture since the beginning of the experiment. Since the end of the 14 d treatment, neurons were given fresh media every 3 d and incubated at 37°C. A large overview shows the presence of five inclusions (arrows) in close proximity. The inclusions appear to be at various stages of development within the neurites, some containing an extensive collection of cytoskeletal filaments (Fil). Magnified views of the enclosed areas are seen in **B** and **C**. Mito (Mitochondria), Lys (lysosome/autolysosome), MVB (multivesicular body). Scale bar = 1 µm and 0.5 µm for insets.

### IFN-γ-treated samples exhibit lysosomal leaking of PFFs

The mechanism by which internalized oligomeric α-syn, which is in the lumen of the endosomal system, leads to seeding and misfolding of endogenous α-syn found in the cytoplasm has been a topic of study for many years ^40^. More recently, the presence and characterization of lysosomal membrane permeabilization have provided clues as to how this process might occur ^41,42^. Here, we directly observe lysosomal rupture and leakage in the PFF + IFN-γ treated samples at early time points (Fig. S6A), where ruptured lysosomes have nanogold PFFs within their lumen and immediately outside their lumen. Coinciding with leakage of PFF into the cytosol, IFN-γ-treated samples show many large autolysosome/autophagosome structures containing organelle fragments (Fig. S6B), indicating impairment in autophagy ^43^. In the 14 d PFF + IFN-γ samples, where neurons have formed inclusions, leakage of PFFs into the cytosol, is a common occurrence (Fig. S6C). Multiple examples of PFFs leaking outside of lytic compartments can be seen in the collection of LB-like inclusions shown in Fig. S3A-G.

### Biochemical extraction of inclusions

We extracted the inclusions formed in DA neurons by subcellular fractionation. Their large size allowed for pelleting with low-speed centrifugation after cellular homogenization. DA neurons undergoing the dual hit treatment regime exhibited an accumulation of PFF in the pelleted inclusions compared to the total homogenate (Fig. S7A), whereas those not receiving IFN-γ treatment had a far less prominent accumulation of PFF. The same trend was present when examining the accumulation of phospho-α-syn. Compared to DA neurons, U2OS, an osteosarcoma cell line, had no noticeable accumulation of PFF or phospho-α-syn in the pellet fractions in either PFF-only or PFF + IFN-γ treatment (Fig. S7B). Additionally, we noted that the α-syn signal in DA neurons consists of a double band, a likely indication of α-syn phosphorylation in DA neurons that does not occur in U2OS. This is further confirmed by our phospho-α-syn blots. Inclusions formed in DA neurons were concentrated and prepared for EM. Detergent-free cell lysis preserved the structure of the inclusions and the resulting samples showed morphologically similar structures to those observed within DA neurons (Fig. S7C-E). The isolated inclusions were filled with lytic compartments (lysosomes and autolysosomes), MVBs, and most importantly, even in their isolated state, still maintained a relatively intact surrounding membrane, further suggesting that these inclusions are membrane-enclosed structures.

### Activated microglia cause inclusion formation in neighboring DA neurons

Proinflammatory cytokine-producing immune cells infiltrate the central nervous system in multiple sclerosis and other neurodegenerative diseases ^44-48^. To assess whether communication between microglia, which produce both IL-1β and IFN-γ ^31,32,49-51^, and neurons leads to formation of LB-like inclusions in DA neurons, we co-cultured a human microglia-like cell line (HMC3), treated with LPS (10 µg/mL) to induce microglial activation, with fully differentiated DA neurons (Fig. 4A). Neurons co-cultured with activated microglia-like cells formed PFF-positive inclusions. Large-field images of co-cultured microglia and neuronal samples reveal that DA neurons neighboring microglia-like cells were most affected by microglial secretions (Fig. S8A). These neurons have significantly higher PFF fluorescence than the control (i.e., neurons co-cultured with PBS-treated microglia-like cells; Fig. 4B). Media from microglia exposed to phosphate-buffered saline (PBS), PFF, LPS, and LPS + PFF were collected and analyzed for the presence of IFN-γ (Fig. 4C). Microglia-like cells treated with LPS, PFF, and LPS + PFF showed elevated levels of IFN-γ. It is, therefore, clear that IFN-γ are released by microglia and lead to inclusion formation in neurons. Consistent with previous findings, PFF treatment alone elicited a microglial response ^52^, suggesting a positive feedback loop might be at play.

**Figure 4.**
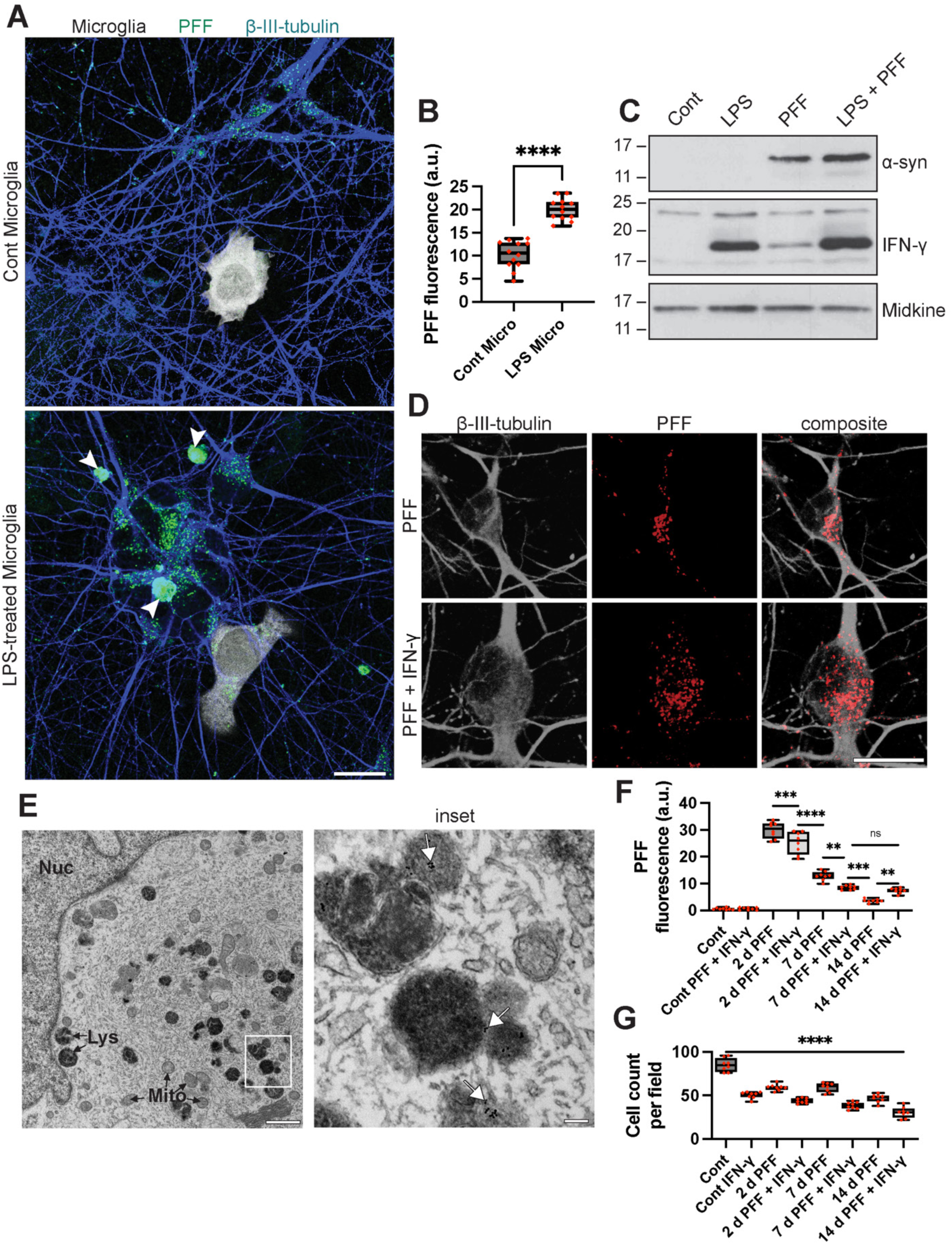
Microglial co-culture led to inclusions in DA neurons while cortical neurons did not form inclusions with IFN-γ treatment. (**A**) DA neurons previously exposed to PFF for 48 h and given fresh media for 72 h were co-cultured with LPS-treated and PBS-treated microglia (previously treated with Cell Tracker Violet dye). Microglia, HMC3 cell line, were co-cultured with neurons for 48 h prior to sample fixation and staining for β-III-tubulin. Neurons co-cultured with PBS-treated (Cont) Microglia exhibited small PFF puncta, while those co-cultured with LPS-treated microglia showed large PFF-positive inclusions. Scale bar = 20 µm. (**B**) PFF fluorescence of DA neurons co-cultured with microglia treated with LPS (or PBS for control: Cont) was quantified where n = 12 for each condition, and data was collected from three independent experiments. (**C**) Microglia were treated with PBS (Cont), LPS, PFF, and PFF + LPS for 24 h. They were then washed with trypsin to remove PFF from the cell surface. They were washed three times with PBS, given serum-free media, and incubated for 48 h. Media were then collected, concentrated, and secreted proteins were analyzed for each condition. Equal amounts of protein from the media were resolved by SDS-PAGE followed by a WB to detect secretory proteins. Microglia treated with LPS, PFF, and LPS + PFF secreted IFN-γ, with the LPS + PFF condition showing the most IFN-γ expression. Microglia treated with PFF also secreted α-syn, with the LPS + PFF treated microglia showing higher levels of α-syn release. Midkine secretion is shown as a loading control. (**D**) iPSC-derived forebrain cortical neurons, derived from iPSCs, underwent the 14 d dual hit treatment, with the control receiving PBS treatment while the IFN-γ condition was exposed to IFN-γ. Scale bar = 20 µm. (**E**) Samples were prepared in the same way described in **D** but given nanogold-labeled PFF and prepared for EM. Dark lysosomes and autolysosomes are scattered across the cytosol and not packed into inclusions; Lys (lysosomes and autolysosomes) and Mito (mitochondria). White arrows point to nanogold-labeled PFF. Scale bar = 1 µm and 100 nm for inset. (**F** and **G**) 14 d experiments were conducted to explore the effects of PFF and IFN-γ on cortical neurons with PFF fluorescence and cell count being quantified, n = 8 for each condition. Data were statistically analyzed using one-way ANOVA and PFF fluorescence data was further analyzed using *post-hoc* Tukey’s test.

### The formation of LB-like inclusions using the dual hit treatment is primarily restricted to DA versus other neuronal types

Although the substantia nigra is most studied for its degeneration in PD, cortical neurons also degenerate in this disease ^53^. We generated forebrain cortical neurons from the same lines used to produce DA neurons (Fig. S8B) and subjected them to the 14 d dual hit treatment regime. Cortical neurons did not form PFF-positive LB-like inclusions up to the 14 d time point (Fig. 4D). They did contain aberrant, dark, and large lysosomes (Fig. 4E), similar to the lysosomes observed within the LB-like inclusions of DA neurons; however, unlike DA neurons, the cortical neurons were unable to package these structures into LB-like inclusions, i.e., structures that are compact, tightly packed with multiple vesicular and organellar structures, and distinct in morphology. Neurogenin 2 (NGN2)-induced neurons, broad spectrum excitatory neurons also failed to form inclusions (Fig. S8C and D) but exhibited dark lysosomes filled with PFF throughout the cytoplasm. Quantification of PFF fluorescence in cortical neurons over 14 d (Fig. 4F) showed a similar trend to DA neurons (Fig. 1E); however, overall cell numbers were markedly decreased in the PFF and IFN-γ- treated samples in cortical neurons (Fig. 4G) relative to DA neurons (Fig. 1F). Since the reduction in cortical neurons occurs very quickly, we rule out the possibility that the formation of LB-like inclusions causes the death of the neurons, as the formation process takes some time to occur.

The specificity of inclusion formation in DA neurons was further explored by examining neuroblastoma (SH-SY5Y), glioblastoma (U87), and osteosarcoma (U2OS) cells. When treated with the dual hit regime, none of these cells formed inclusions (Fig. S9A and B); however, each cell line exhibited signs of lysosomal abnormalities: a loss of LAMP1 and an increase in lysosomal size (Fig. S9C and D). Thus, treatment of various cell types with PFF and IFN-γ causes lysosomal defects, but the formation of LB-like inclusions using the dual hit treatment is restricted to DA neurons.

### The formation of LB-like inclusions is dependent on the endogenous levels of α- syn

α-syn is present in LBs ^1,2,54,55^. The phosphorylation of α-syn is considered the best marker for α-syn aggregation and LB formation ^56^. To explore whether the inclusions formed in our treatment regime contain phosphorylated α-syn, DA neurons that underwent the dual hit regime were stained with phospho-α-syn antibody. These neurons formed phospho-α-syn-positive inclusions (Fig. 5A), and PFFs were localized in inclusions along with phospho-α-syn (Fig. 5B). Supplementary Table 1 displays all the attributes of the LB-like inclusions formed using the 14 day treatment regime.

**Figure 5.**
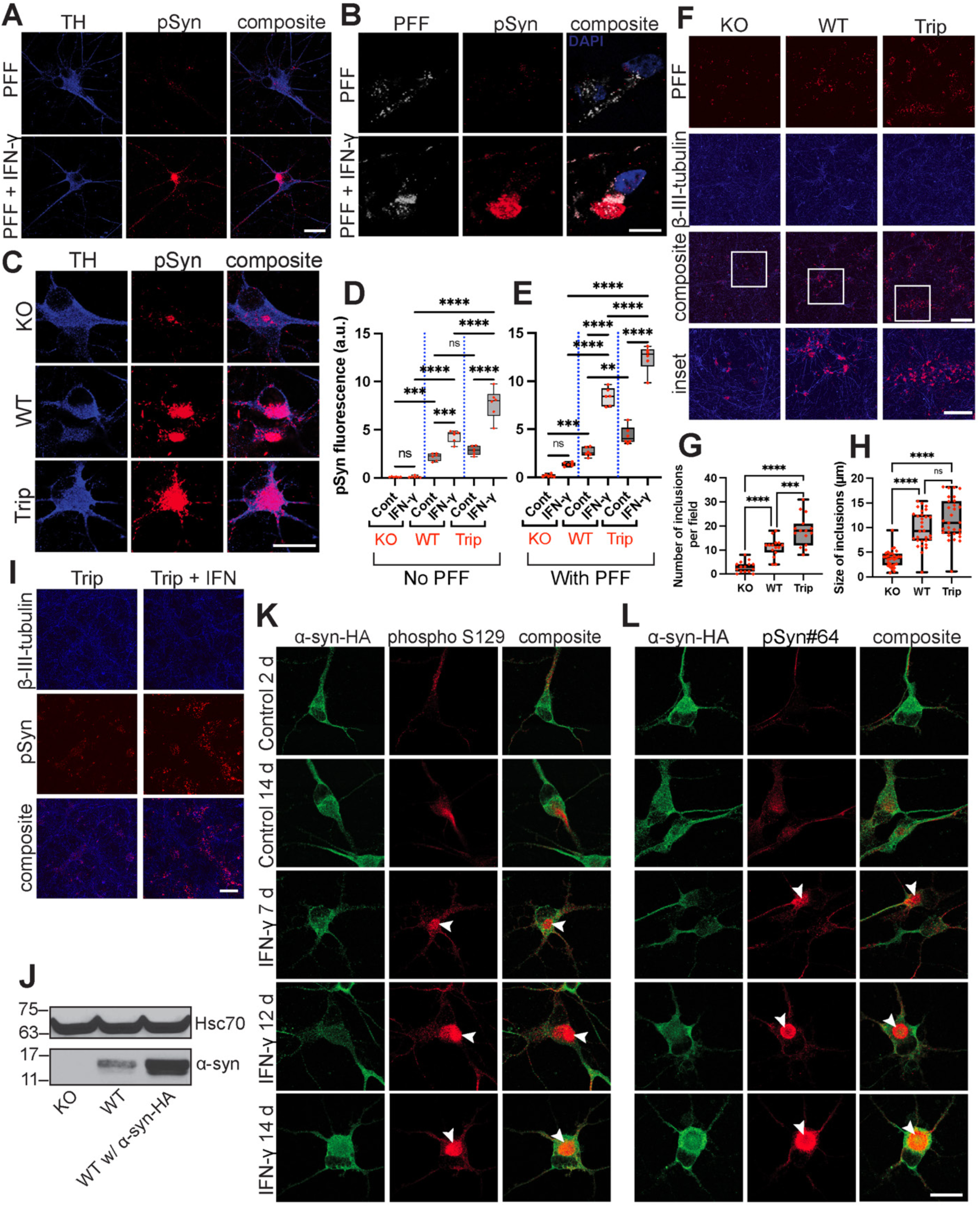
Phospho-α-syn is a marker for inclusions and is dependent on the endogenous pool of α-syn. (**A**) iPSC-derived DA neurons underwent the 14 d dual hit treatment regime, in which unlabeled PFFs were administered to neurons for 48 h, incubated in fresh media for 72 h, followed by the administration of IFN-γ (or PBS for control) for 24 h. Neurons were then incubated for 8 d in fresh media, fixed, and stained with tyrosine hydroxylase antibody (TH, blue) and phospho-α-syn (pSyn) antibody (red). Scale bar = 20 µm and 10 µm for insets. (**B**) Neurons underwent the same experiment as described in **A**, except fluorescent PFFs were used to ascertain localization. Both PFFs and phospho-α-syn localized to inclusions. Scale bar = 10µm. (**C**) DA NPCs derived from patient cells with *SNCA* triplication were knocked out to WT (two copies of *SNCA*) and KO (no *SNCA*). NPCs were then differentiated into neurons, and the dual hit treatment regime was carried out using unlabeled PFF. Scale bar = 20 µm. (**D**) Neurons with different levels of endogenous α-syn expression underwent the dual hit treatment regime without PFF treatment. (**E**) phospho-α-syn fluorescence was calculated in neurons that underwent the dual hit treatment regime with fluorescently labeled PFF, n = 6 for each condition. A threshold was set to exclude small and faint PFF puncta. (**F**) DA neurons with different number of copies of *SNCA* underwent the dual hit assay. Scale bar = 200 µm and 100 µm for insets. (**G**) The number of PFF-positive inclusions per field in experiment **F** were quantified, with only inclusions larger than 2.0 µm included in the calculation, n = 15 for each condition (**H**) Size of PFF-positive inclusions in the experiment described in **F**, were quantified, following thresholding. n = 30 for each condition, collected from three independent experiments. Data in **D**, **E**, **G**, and **H** were collected from three independent experiments, and one-way ANOVA and Tukey’s test were used to analyze the data. **** denotes that p < 0.0001, *** denotes p < 0.001, ** denotes p < 0.01, and ns = not significant. (**I**) Phospho-α-syn staining in SNCA triplication line (in absence of PFF) shows possible inclusions even without IFN-γ; however, number of inclusions is drastically increased with the treatment of IFN-γ for 24 h. Cells were maintained until 14 d and then fixed and stained. (**J**) WT DA neurons were transduced with α-syn-HA adenovirus DA for 48 h, given fresh media for 72 h, and collected for Western blotting. Transduced DA neurons showed higher levels of α-syn expression compared to WT and *SNCA* KO neurons. (**K** and **L**) WT neurons transduced with α-syn-HA adenovirus then underwent the dual hit treatment regime shown in Fig. S10. Neurons were then fixed and stained for phospho-α-syn antibodies. Neurons not treated with IFN-γ did not form inclusions at the earlier time point, immediately following α-syn-HA adenovirus transduction at day 2, nor did they show inclusions at day 14. Neurons treated with IFN-γ begin showing accumulation of phospho-α-syn in the cell body at day 7. This accumulation becomes more apparent and more prominent with an increase in incubation time, leading to large phospho-α-syn positive accumulations (white arrowhead). Scale bar = 20 µm.

To assess the influence of endogenous α-syn levels on LB-like inclusions, we used iPSCs from a patient with an *SNCA* triplication, in which we used a double knockout (KO) to generate wildtype (WT) cells and a quadruple KO to generate an *SNCA* null. These isogenic lines were differentiated into DA neurons and exposed to PFFs and IFN-γ. The level of phospho-α-syn fluorescence was influenced by α-syn expression levels more so than the addition of PFFs (Fig. 5C, D, and E). We observed a striking increase in phospho-α-syn fluorescence following treatment with IFN-γ. α-syn expression levels also influenced the formation of PFF-positive inclusions with higher levels leading to a progressive increase in the number and size of these structures (Fig. 5F, G and H). We noted, however, that SNCA triplication, alone, could form phospho-α-syn-positive inclusions, but at a much smaller level than when coupled with IFN-γ administration (Fig. 5I)

We next transduced WT neurons with α-syn-HA adenovirus and treated them with IFN-γ (Fig. S10 and Fig. 5J). We observed formation of phospho-α-syn-positive inclusions in the absence of PFFs (Fig. 5K and L). In summary, IFN-γ can also induce the formation of inclusions using endogenously expressed α-syn.

### Downregulation of lysosomal proteins and disruption in autophagic flux coincides with the formation of LB-like inclusions

Given the numerous qualitative defects in lysosomal morphology seen in cells treated with PFF and IFN-γ we sought to examine if this treatment alters proteins regulating lysosomal biogenesis and function. NRF2 and TFEB are transcription factors regulating the expression of lysosomal proteins and function ^57-61^. As LAMP1 and LAMP2, major lysosomal proteins have previously been implicated in PD ^19^, and both TFEB and NRF2 have been suggested to play a role in neurodegenerative diseases ^62,63^, we sought to examine the expression levels of these proteins. While LAMP1 and LAMP2 expression were higher in cortical neurons than DA neurons at baseline conditions, treatment with PFF and IFN-γ caused a strong reduction in LAMP1 and LAMP2 levels in DA neurons but not in cortical neurons. TFEB and NRF2 followed a similar trend and were downregulated in DA neurons but remained nearly unaffected in cortical neurons (Fig. 6A). Using CellROX and MitoSOX to look at generalized oxidative stress and superoxide production, respectively, we found that general oxidative stress levels were similar in both DA and cortical neurons (Fig. 6B); however, the production of superoxide species was found to be significantly different, as cortical neurons treated with PFF and those treated with PFF + IFN-γ exhibited significantly higher levels of MitoROX fluorescence compared to DA neurons (Fig. 6C).

**Figure 6.**
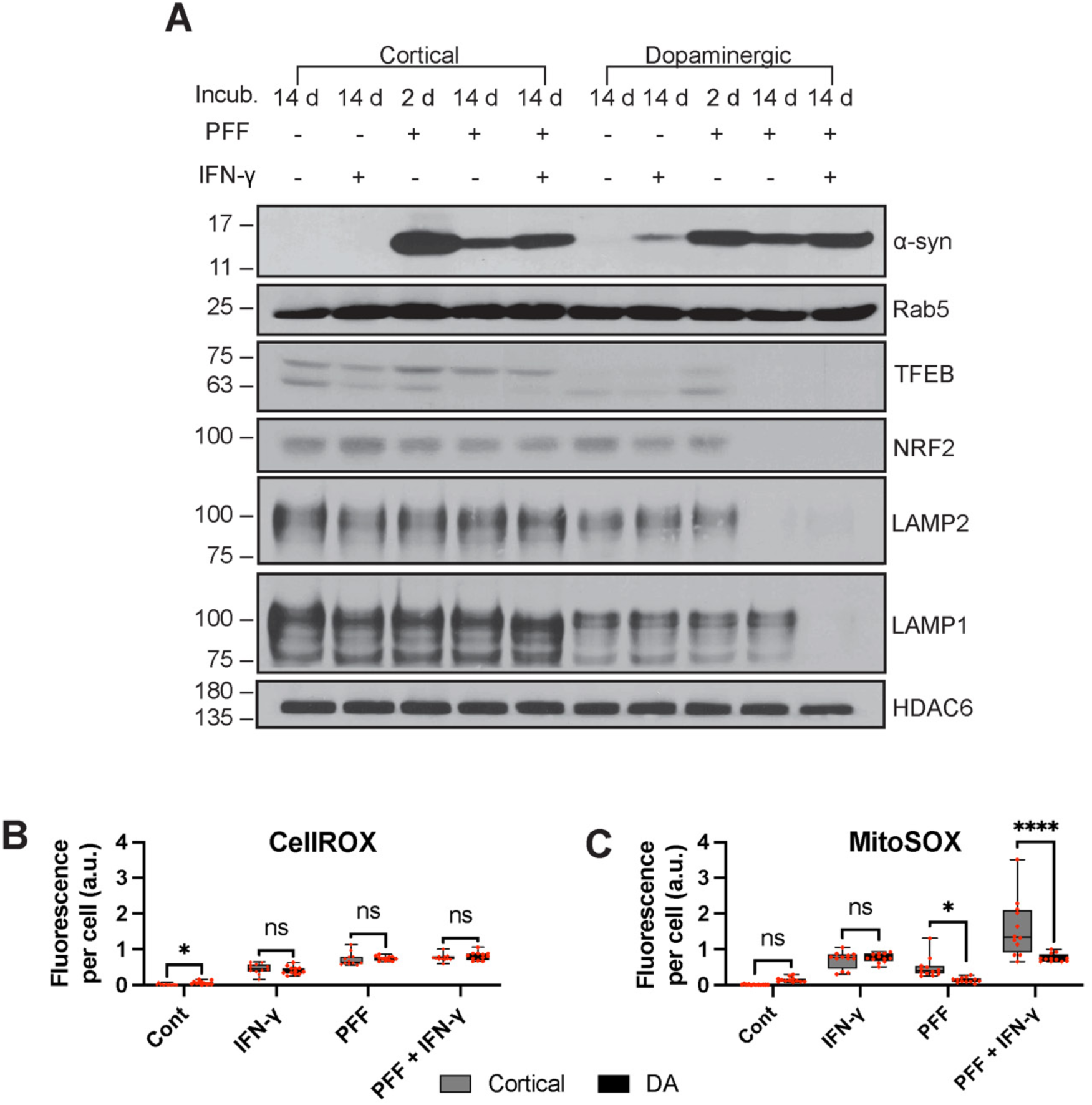
The dual hit treatment resulted in the downregulation of lysosomal proteins. (**A**) Protein expression in cortical and DA neurons were measured across five conditions. DA neurons show downregulation in LAMP1, LAMP2, NRF2, and TFEB, to a greater extent than cortical neurons in response to PFF and IFN-γ. At base line, LAMP1 and LAMP2 expression is much higher in cortical than in DA neurons. This is also the case with TFEB. Additionally, treatment with IFN-γ alone resulted in elevated α-syn expression in DA but not cortical neurons. Western blot is representative of three separate experiments done with neurons derived using the same iPSC NPCs. (**B** and **C**) Cortical and DA neurons were plated onto separate glass-bottomed dishes, given CellROX and MitoSOX, and imaged live at 14 d across four conditions. n = 12 for each condition, collected from three independent experiments. Two-way ANOVA and *post-hoc* Tukey’s test were done to compare each condition to the control. **** denotes p < 0.0001, ** denotes p < 0.01, and ns = not significant.

Due to their constant need to mitigate oxidative stress generated through the production and metabolism of dopamine ^64,65^, DA neurons are probably more adept at dealing with oxidative stresses, partly through the sequestration of damaged organelles. Should oxidative species impair organellar function, specifically mitochondria, the release of cytochrome c can lead to Caspase-dependent cell death ^66^. Furthermore, protecting the cell from leaky lysosomes, another consequence of oxidative stress ^67^, is also vital for cell survival ^68^. This is why the sequestration of damaged organelles is protective. Cortical neurons, unable to form LB-like inclusions, leave themselves vulnerable to the consequences of oxidative stress. This is perhaps why DA neurons can survive at much higher rates than cortical neurons when confronted with our treatment regime.

By tracking LC3B staining, a marker for autophagy ^69^, over the 14 d treatment regime, we found that large autophagosomes form, merge, and then eventually result in the even larger PFF-positive structures (Fig. 7A and B), similar to the inclusions we have repeatedly observed. These large LC3B-positive structures are only present in PFF + IFN-γ treated samples. The accumulation and perseverance of these large LC3B structures is indicative of autophagic dysfunction and the inability of the cell to clear the contents within these autophagosomes ^70^. This, along with their membrane-bound morphology, provides further proof that the LB-like inclusions that form are in fact dysfunctional autophagosomes.

**Figure 7.**
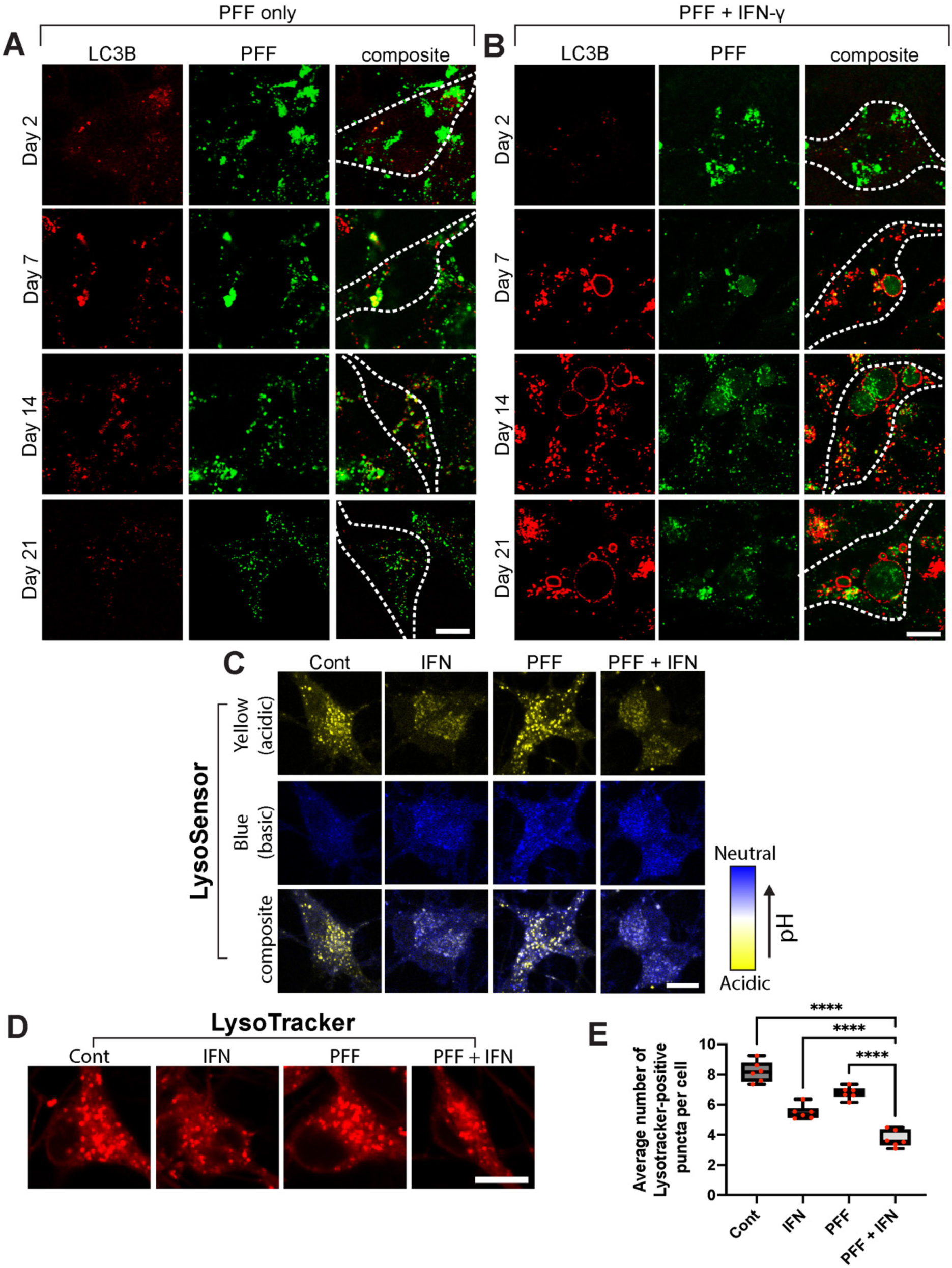
Inclusions forming in DA neurons are enveloped by LC3B, PFF and IFN-γ impair autophagic flux by altering lysosomal pH. (**A** and **B**) Neurons undergoing the 14 d treatment regime, were tracked overtime for their staining of LC3B. At day 2, just following the administration of PFF barely any LC3B staining can be seen. Following 48 h following IFN-γ administration on day 5, the PFF + IFN-γ sample show large LC3B positive structures. LC3B staining is also high in the PFF only samples, but the neurons in this condition do not possess large LC3B structures. While the LC3B staining abates in the PFF only sample following the passage of time, the LC3B staining in the PFF + IFN-γ sample remains high and large LC3B-positive autophagosomes still remain at 21 d. Scale bar = 10 µm. (**C**) Lysosomal acidity was ascertained in DA neurons at day 7 of the dual hit treatment regime using LysoSensor (Thermo Fisher Scientific). The IFN-γ-treated neurons showed much higher lysosomal pH (more blue staining) compared to the PFF-only and the control neurons. Scale bar = 10 µm. (**D**) Lysosomal pH was also tested using LysoTracker (Thermo Fisher Scientific) which fluoresces at a pH of ∼ 6. Scale bar = 10 µm. (**E**) LysoTracker-positive punctae were counted using ImageJ (NIH) and divided by the number of cells per field across conditions. Data was collected from three independent experiments, (n=6 for each condition). One-way ANOVA and Tukey’s test were used to analyze the data. **** denotes that p < 0.0001.

Furthermore, we analyzed lysosomal pH using LysoSensor and LysoTracker as done previously by Guerra et al. ^71^, in DA neurons following the 14 d dual hit treatment. We found that neurons treated with IFN-γ or PFF + IFN-γ presented with lysosomes with a higher pH compared to the control and PFF-treated samples (Fig. 7C). This was complemented by our Lysotracker staining, which showed that neurons treated with IFN-γ or PFF + IFN-γ had lower numbers of LysoTracker-positive lysosomes (Fig. 7D and E).

Lastly, the expression levels of lysosomal proteins were also examined in SH-SY5Y, U87, and U2OS (Fig. S11), cells that did not form inclusions. Although NRF2, LAMP1, and LAMP2 levels were decreased in samples treated with PFF + IFN-γ, TFEB was unaffected, indicating that impairment of lysosomal biogenesis is critical for a more detrimental impairment of lysosomal function, which may lead to the formation of inclusions.

### Formation of LB-like inclusions is prevented by Perillaldehyde (PAH)

As NRF2 and LAMP2 are highly downregulated in samples treated with both PFF and IFN-γ we searched to use a chemical activator for the lysosomal-inflammasome pathway and found PAH to be a perfect candidate. PAH is an activator of NRF2 and its ability to activate NRF2 and therefore increase lysosomal activity and degradation has been shown repeatedly in previous studies ^72-75^. To reduce/counteract the inclusion-forming effects in neurons caused by PFF and IFN-γ, PAH treatment experiments were conducted on neurons treated with PFF + IFN-γ (Fig. 8A and B). A cell survival assay indicated that PAH was cytotoxic at higher concentrations (Fig. 8C), therefore the concentration of 10 μM of PAH was used to minimize cell death related to PAH treatment. Samples treated with PAH 1 d following IFN-γ treatment, exhibited significantly less PFF-positive inclusions compared to those not treated with PAH (Fig. 8D). The 1 d samples also exhibited significantly smaller inclusions compared to the PAH-untreated condition (Fig. 8E).

**Figure 8.**
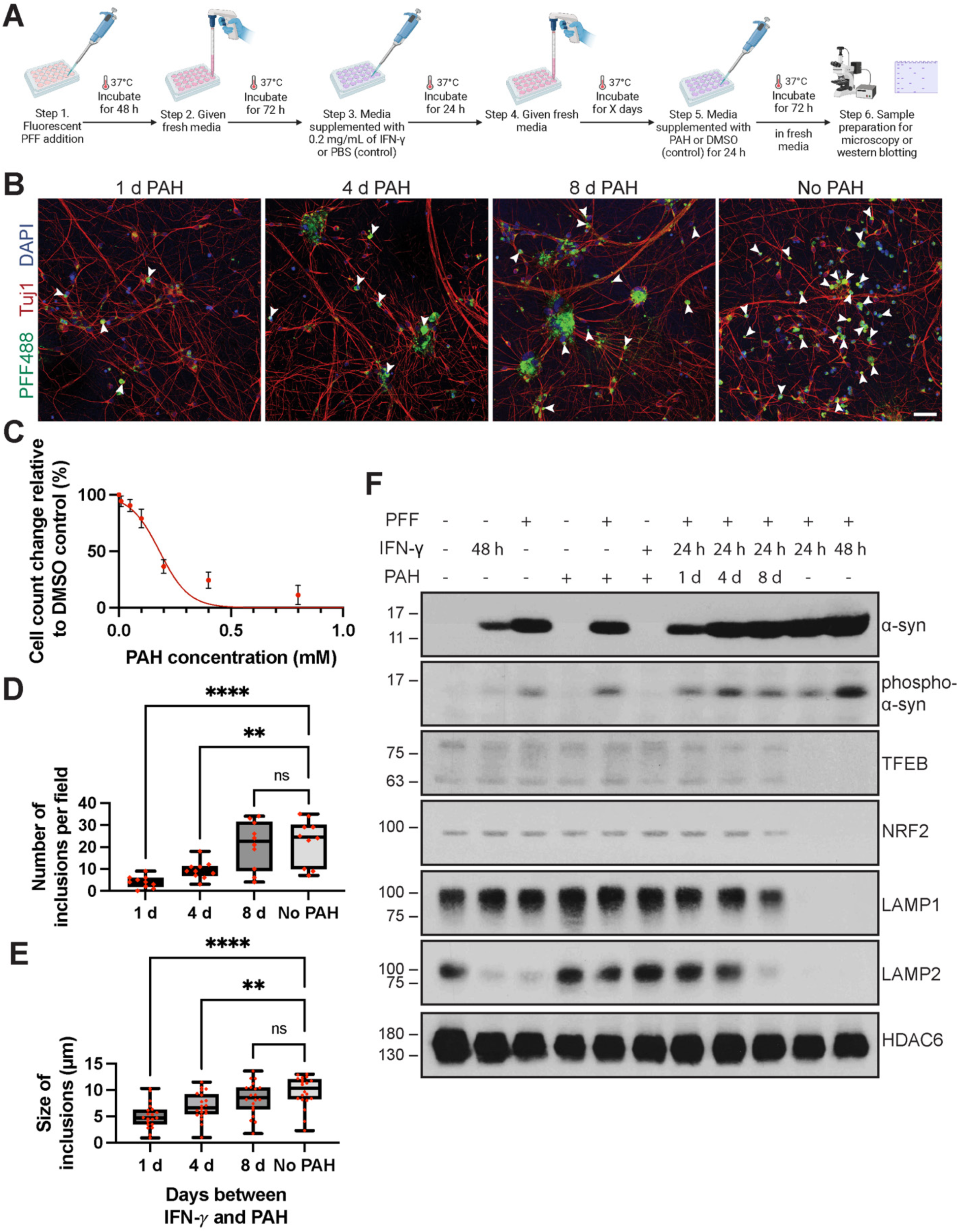
PAH treatment reduces inclusion formation and restores lysosomal protein expression. (**A**) An overview of the PFF, IFN-γ, and PAH treatment regime. DA neurons were given fluorescently labeled PFF for 48 h and then fresh media for 72 h, followed by treatment with IFN-γ for 24 h. All conditions were then given fresh media. PAH was administered at different time points following IFN-γ administration at a concentration of 10 µM. Some conditions received no PAH treatment (**B**) Neurons treated with PAH, or DMSO as a control, were fixed, stained with Tuj1 and DAPI, and prepared for fluorescence microscopy. Arrowheads point to PFF-positive inclusions. Scale bar = 80 µm. (**C**) PAH concentration was determined through a cell survival experiment, where DA NPCs were differentiated into neurons on 24-well plates. They were then treated with different concentrations of PAH for 24 h, stained with Hoechst, and the fluorescence of each well was calculated using a plate reader, with laser excitation set at 400 nm and emission detectors set to 450 nm. A rapid decline in neuronal survival occurs at concentrations higher than 0.2 mM. n = 6 for each condition. A log (agonist) vs. normalized response function was used to calculate the survival curve. (**D**) The number of inclusions per field (387.5 µm by 387.5 µm) was quantified with inclusions larger than 2.0 µm being incorporated in the calculation, n = 10 for each condition, and data was collected from five independent experiments. (**E**) The size of inclusions in each condition was calculated following thresholding of images to exclude faint and smaller PFF puncta, n = 20 for each condition, and data was collected from five independent experiments. Data in **D** and **E** were analyzed using one-way ANOVA and *post hoc* Tukey’s test. (**F**) Western blot was conducted to analyze protein expression changes in response to PAH exposure in DA neurons that underwent the treatment described in Fig. 8A. An increase in IFN-γ incubation time results in even more α-syn signal coupled with a higher phospho-α-syn signal. A decrease in the time between PAH treatment following IFN-γ results in the upregulation of LAMP1, LAMP2, TFEB, and NRF2. Western blot is representative of three separate experiments done with neurons derived using the same iPSC NPCs.

Lastly, we found that TFEB, LAMP1, LAMP2, and NRF2 protein expression levels were similar to control levels in the 1 d and 4 d PAH conditions (Fig. 8F). IFN-γ-only conditions exhibited a higher level of α-syn expression than control, as was observed previously (Fig. 6A). In addition, an increase in IFN-γ incubation time in PFF-treated neurons also resulted in higher levels of α-syn. Taken together, these results demonstrate that the DA neurons undergoing the dual hit treatment regime exhibit lysosomal impairment through the downregulation of LAMP1 and LAMP2, and the transcription factors NRF2 and TFEB, which creates an environment allowing for the formation of large LB-like inclusions. This phenotype can be partially prevented by recovering these lysosomal proteins after 1 d of PAH treatment.

## DISCUSSION

LBs are a hallmark of PD, LBD and other synucleinopathies. Although many studies have reported the ultrastructure of LBs, the reliance on post-mortem tissue has resulted in less-than-ideal analysis and an ongoing dispute regarding their origins. Many studies report that LBs are cytoplasmic structures, filamentous, and lacking a delineating membrane ^2,12,76-80^. However, a recent study using a shorter post-mortem interval and better processing of post-mortem tissues indicates that LBs contain filaments but also numerous organelles, vesicles, and membrane fragments ^10^. Despite this, limited information can be understood from post-mortem tissue samples, such as the formative stages involved in assembling LBs, their progression, and the incorporation of organelles and membranes. Here we determine that a dual treatment of a PD insult (fibrils of α-syn) and an immune challenge (IFN-γ or IL-1β) leads to structures with a remarkable similarity to LBs, and this occurs specifically in DA neurons. This has allowed us to study the stages involved in their formation and their morphology. The LB-like inclusions formed by this method are PFF-positive, phospho-α-syn-positive, LC3B-positive, membrane-enclosed, and contain a medley of organelles including lysosomes, mitochondria, MVBs, ER, and cytoskeletal filaments (Supplementary Table 1). Our findings bring together two separate fields of inquiry into the pathophysiology of PD: (1) the internalization, propagation, seeding, overexpression, and incorporation of α-syn in LBs ^81^; (2) the involvement of the immune system in PD ^23^.

IFN-γ is one of many proinflammatory cytokines released by microglia and other immune cells that can serve as an immune challenge. IL-1β used in this paper, tumor necrosis factor, and other cytokines are also released by immune cells ^82^. These cytokines could also contribute to an environment in which DA neurons can form inclusions in PD. In our co-culture experiments, we found that DA neurons readily formed inclusions within 48 h following co-culturing with microglia. Although IFN-γ secretion by microglia was detected and has been reported previously ^83,84^, it is likely a combination of microglia-secreted factors driving this effect *in vitro*, including IL-1β. In addition to microglia, we believe that neuronal exposure to IFN-γ may also occur through immune cell secretions, following their infiltration into the brain ^45^, as immune cells are robust producers and secretors of IFN-γ ^81^. In fact, aging microglia have been shown to facilitate immune system infiltration into the brain ^85^. Exposure to IFN-γ can induce antigen presentation in neurons ^81^, which may lead neurons to be further vulnerable to the immune system via autoimmune-mediated mechanisms ^86^, and perhaps more likely to form inclusions as a response to α- syn upregulation and oxidative stress ^87,88^. Overall, our goal is not to implicate any one specific cell type for the secretion of proinflammatory cytokines, or that one specific cytokine is responsible for inclusion formation. We simply present a model in which proinflammatory cytokines IFN-γ and IL-1β administered alongside a PD-insult, result in the formation LB-like inclusions.

Due to their formation in cultured cells, we can examine LB-like inclusions with ultrastructural integrity not currently possible with post-mortem tissue. This has allowed us to make a significant observation regarding the structure of these LB-like inclusions: they are membrane-bound. Whether LBs are membrane-enclosed has significant implications for their origin. If LBs are formed in the cytoplasmic compartment aided by the fibrilization of α-syn, which allows for the tethering and pulling of organelles and membranes together, then α-syn aggregation becomes the central event in inclusion formation and will lead to the formation of LBs given enough time ^84^. If LBs are membrane-bound, they may form within the lumen of the endo/lysosomal system. This would rely on impairments in autophagy and lysosomal function and would be less dependent on α-syn aggregation. Our *in vitro* LB-like inclusions were consistently membrane-bound across all our EM data, even when biochemically isolated. Coupled with our findings regarding the downregulation of lysosomal transcription factors and proteins, our data support the latter hypothesis that LBs are membrane-bound and ultimately form in the lumen of ever expanding, dysfunctional autophagosomes since the PFF-positive inclusions that form are LC3B-positive and enveloped by a double membrane.

As shown in our model (Fig. S12), α-syn aggregation occurs, for the most part, in the cytosol. This is initiated by cytosolic seeding of aggregation through the leaking of PFFs from lysosomes into the cytosol. Downregulation of LAMP2 results in impaired chaperone-mediated autophagy, resulting in a buildup of monomeric α-syn in the cytosol, contributing to aggregation. While aggregating in the cytosol, phosphorylation of α-syn also takes place. The cell, to degrade aggregates, induces autophagy, and takes up aggregated α-syn along with damaged organelles. Contents within autophagosomes are not degraded however due to dysfunctional lysosomal activity, and the lack of lysosomal fusion with autophagosomes, which is caused by the downregulation of LAMP1 and LAMP2. Autophagosomes continue to take up more aggregates from the cytosol but continue building in size as no degradation takes place. These events finally lead to the formation of membrane-bound inclusions.

In our study, PFF and IFN-γ cooperatively enable the downregulation of lysosomal transcriptional factors and lysosomal proteins, most notably LAMP2. Downregulation of LAMP2 inhibits the degradative activity of lysosomes by blocking chaperone-mediated autophagy and autophagosome-lysosomal fusion ^89^. LAMP2A, the isoform responsible for chaperone-mediated autophagy, has been previously studied for its potential role in PD ^89^. The mutant forms of α-syn are responsible for altering the chaperone-mediated autophagy activity of LAMP2A ^90^. LAMP2 downregulation in brain regions has also been associated with increased α-syn aggregation in those regions ^91^. Whether PFF and IFN-γ downregulate LAMP2 expression by affecting NRF2 expression, and, therefore, the transcription of LAMP2, or by affecting LAMP2 expression through other means remains to be studied. Still, the downregulation of NRF2 ensures that the cell cannot increase LAMP2 expression when needed as an inflammasome response ^92^. While EM evidence does not suggest that the number of lytic compartments decreases as a result of PFF and IFN-γ exposure, our Western blot and lysosomal pH data clearly indicate an impairment of the lysosomal compartments, with high pH and a decrease in lysosomal proteins. It is therefore important to note that while clear signs of lysosomal dysfunction and lysosomal protein downregulation are observed, the net lysosomal biogenesis – i.e., the number of electron dense vesicles – is not affected.

It is also worth noting that the lack of inclusion formation in cortical neurons in our treatment regime is not an indicator of LB formation in PD. Clearly, cortical neurons form LBs in PD as shown by Braak *et al.* ^3^; however, the mechanism by which these neurons are pushed into forming inclusions is perhaps different and may rely on α-syn overexpression as other models have suggested ^45^. In our model, we find that DA neurons respond to the dual hit treatment regime, as they have evolved mechanisms to mitigate the deleterious effects of oxidative stress response ^64,65^, which doesn’t occur with cortical neurons. Through sequestration of damaged mitochondria and leaky lysosomes into inclusions, LB-like inclusions protect DA neurons from harmful species that can play a role in neuronal apoptosis, including the release of cytochrome c from mitochondria and the release of degradative enzymes from lysosomes ^66,67^. This is perhaps why DA neurons survive at much higher numbers than cortical neurons. We believe that the dual hit treatment may, therefore, not facilitate the appropriate environment for the formation of LB-like inclusions in cortical neurons. Different neuronal subtypes probably require different conditions to induce the formation of LB-like inclusions, which needs to be explored in future research.

α-syn likely plays a large role in formation of inclusions, and our findings do not discount that. In patients with overexpression of α-syn due to locus duplication and triplication, and in models where overexpression of α-syn is used to drive inclusion formation, it is very likely that the tethering of organelles by α-syn fibrils does in fact result in inclusions. Our model aims to only suggest another possible way for the formation of inclusions, that has not been studied previously. We also don’t suggest that neuroinflammation is necessary for PD; however, our data suggests that it could be one of the ways in which DA neurons are pushed into making inclusions. It is very likely that multiple pathways are at play in PD: the overexpression of α-syn can drive inclusion formation, and the intercellular transport of α-syn oligomers coupled with immune system activation can also lead to inclusion formation. We believe our results, along with more recent work on inclusions formation shown by Mahul-Mellier *et al.* ^83^, Tanudjojo et al. ^93^, Iannielli et al. ^94^, and Gribaudo et al. ^95^, mark the beginning of new era, where the formation of LB-like inclusions can be studied and the mechanisms involved in the formation can be understood. We hypothesize that there are multiple mechanisms by which neurons can form inclusions, just as there are multiple genetic and environmental risk factors that lead to the onset of PD.

### Limitations

Our goal for this project was to find whether neuroinflammatory signals would influence DA neurons and whether the resulting dysfunctions would mimic those seen in PD. To this end, we were successful, and we clearly delineated a protocol in which LB-like inclusions could be formed and studied *in vitro*. However, future research using *in vivo* models, in which neuroinflammation occurs due to acute bacterial or viral infection, will need to be conducted to assess the physiological relevance of our model. Clearly, due to the *in vitro* nature of our study, the physiological relevance of our model is limited.

Additionally, this paper does not do an exhaustive investigation as to whether the phenotype observed here – i.e., the formation of LB-like inclusions – can only be caused by proinflammatory cytokines as the second hit. While this was not the project’s aim, this is a valid critique. Future research must determine whether toxins and metabolic stressors also push DA neurons to form LB-like inclusions. We hypothesize that this is the case: regardless of the stressor, cellular stress will affect autophagy-lysosomal pathways, and will result in the formation of aggregates and inclusions as the cellular machinery involved in degrading such structures will be impaired.

Lastly, whether the downstream effect of proinflammatory cytokines on the autophagy-lysosomal pathway is solely responsible for the formation of LB-like inclusions needs to be investigated further. As noted previously, proinflammatory cytokines have a multitude of effects on the cell. By knocking down important autophagy-lysosomal-related proteins or blocking the deleterious effects of proinflammatory cytokines on the autophagy-lysosomal pathway, future researchers can determine whether other pathways activated through exposure to proinflammatory cytokines are also responsible for the formation of LB-like inclusions in DA neurons, or whether this is solely due to the impairment of the autophagy and degradation machinery.

## Conclusion

The consistent formation of LB-like inclusions *in vitro* marks the next step in investigating the molecular underpinnings of LB formation and PD pathophysiology. Our *in vitro* treatment regime can serve as a starting point for future investigators to modify our protocol and explore the different stages of LB formation, changes in protein expression, and the onset of protein sequestration into LB-like inclusions. This treatment regime provides a reproducible and unlimited source of LB-like inclusions. It allows for a level of sample preservation and ultrastructural resolution that is currently impossible to attain with post-mortem tissue. Biochemical isolation of these LB-like inclusions at different time points and assessment of their protein contents over time will provide the field with a wealth of information regarding the molecular changes underlying LB formation. Lastly, adapting our protocol for use with midbrain organoids ^96^, brain assembloids ^97,98^, and other 3D models can allow for the recreation of PD *in vitro*, mapping the entire sequence of events in neurodegenerative disorders, ushering in the next era of research into neurodegenerative diseases.

## Acknowledgements

We acknowledge the Advanced BioImaging Facility and the Facility for Electron Microscopy Research at McGill University. We thank Dr. Kelly Sears for all his help with the EM. We thank Dr. Bruce Wright, the head of the Division of Medical Sciences at the University of Victoria. We also thank Drs. Sabatini and Zoncu for the HA-TMEM192 plasmids. AB is supported by Fonds de recherche du Québec doctoral award and a studentship from the Parkinson Society of Canada. This work was supported by financial support from the Canada First Research Excellence Fund, Healthy Brains, Healthy Lives, McGill University, awarded to TMD and PSM. EAF is supported by Foundation Grant from CIHR (FDN-154301) and a Canada Research Chair (Tier 1) in Parkinson disease. TMD is supported by a project grant from CIHR (PJT – 169095). PSM is a Distinguished James McGill Professor and Fellow of the Royal Society of Canada.

## Author Contributions

A.B. planned and conducted the experiments and wrote the manuscript. R.A. conducted inclusions isolation experiments, Western blot experiments, analyzed microglial media for secreted proteins, and aided with the writing of the manuscript. A.A. conducted Western blots for protein expression related to knockdowns and overexpression experiments and helped with the writing of the manuscript and organization of the figures. C.Z. provided us with NPCs and differentiated iPSCs and helped with the designing of the experiments. A.A. conducted Western blots for protein expression related to knockdowns and overexpression experiments and helped with the writing of the manuscript. C.H. provided NPCs and differentiated iPSCs into cortical and dopaminergic neurons. E.B. performed additional Western blotting experiments. E.N.R. helped generate all the dopaminergic NPCs and characterized the SNCA, AIW0020-02, and 3450 lines. W.L. produced PFFs and conducted fluorescent conjugation experiments, while I.S. characterized PFFs via dynamic light scattering. E.D.C.P. characterized and measured PFF size via EM. M.Y., E.A.F., J.A.S., and T.M.D. edited the manuscript and helped with the direction of the project. J.A.S. also aided in the writing of the manuscript and the designing of experiments. T.M.D. supervised and oversaw the production of PFFs. P.C.N. processed electron microscopy samples, helped with the identification of neuronal ultrastructure, and aided in writing the manuscript. P.S.M. funded and supervised the project and wrote the manuscript with A.B.

## Declaration of Interests

The authors declare no competing interests.

## Figures, Tables, and Legends

## STAR Methods

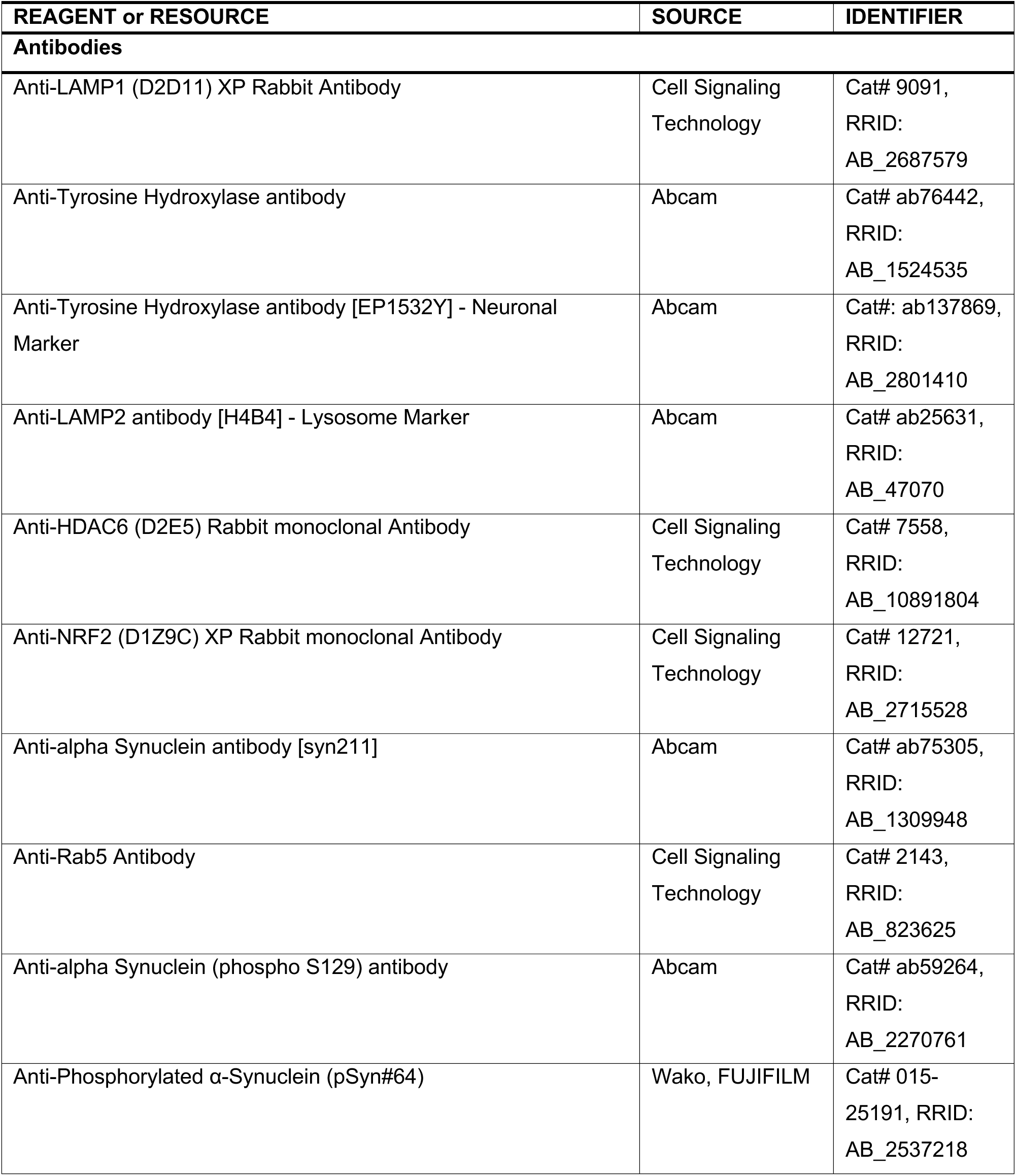
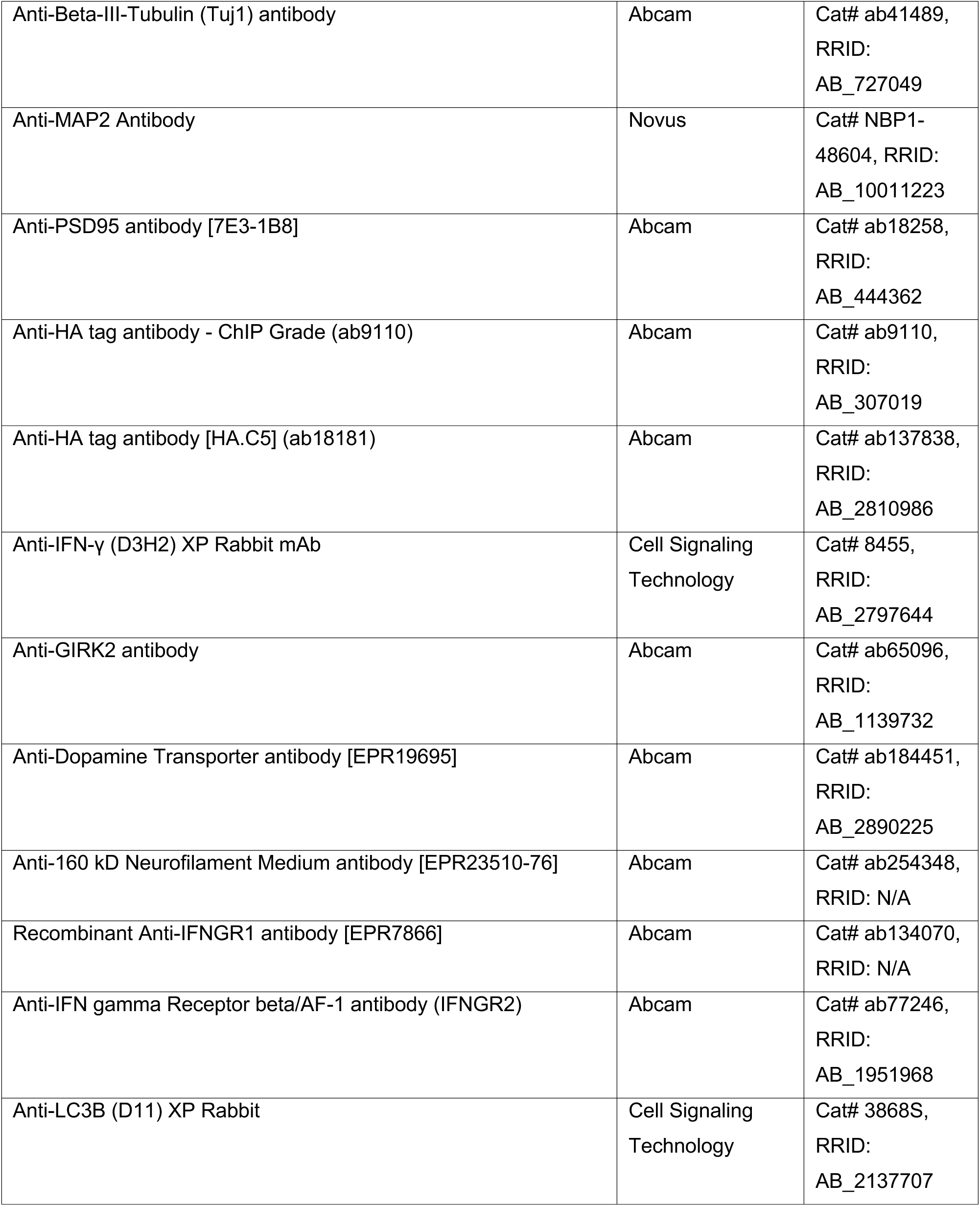
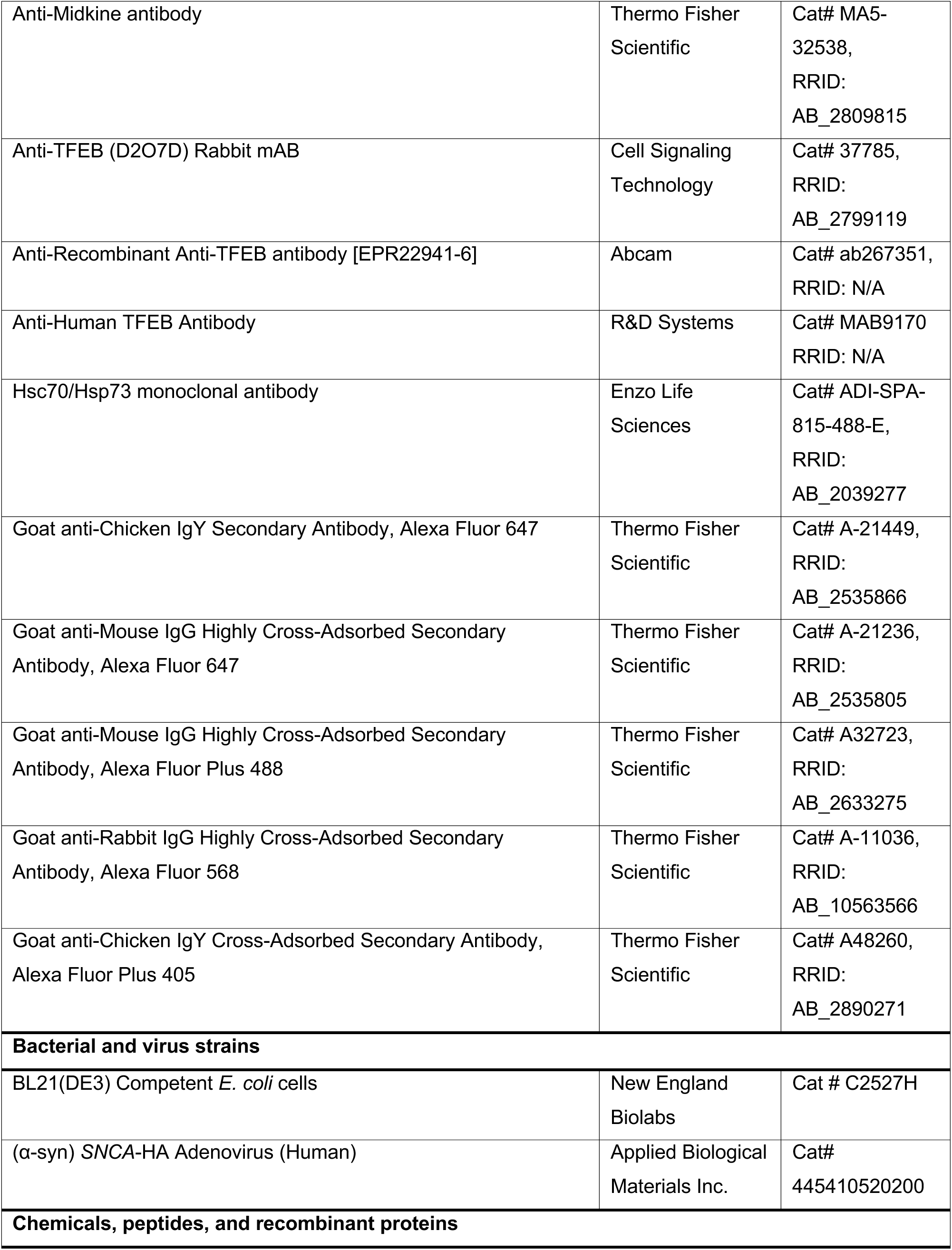
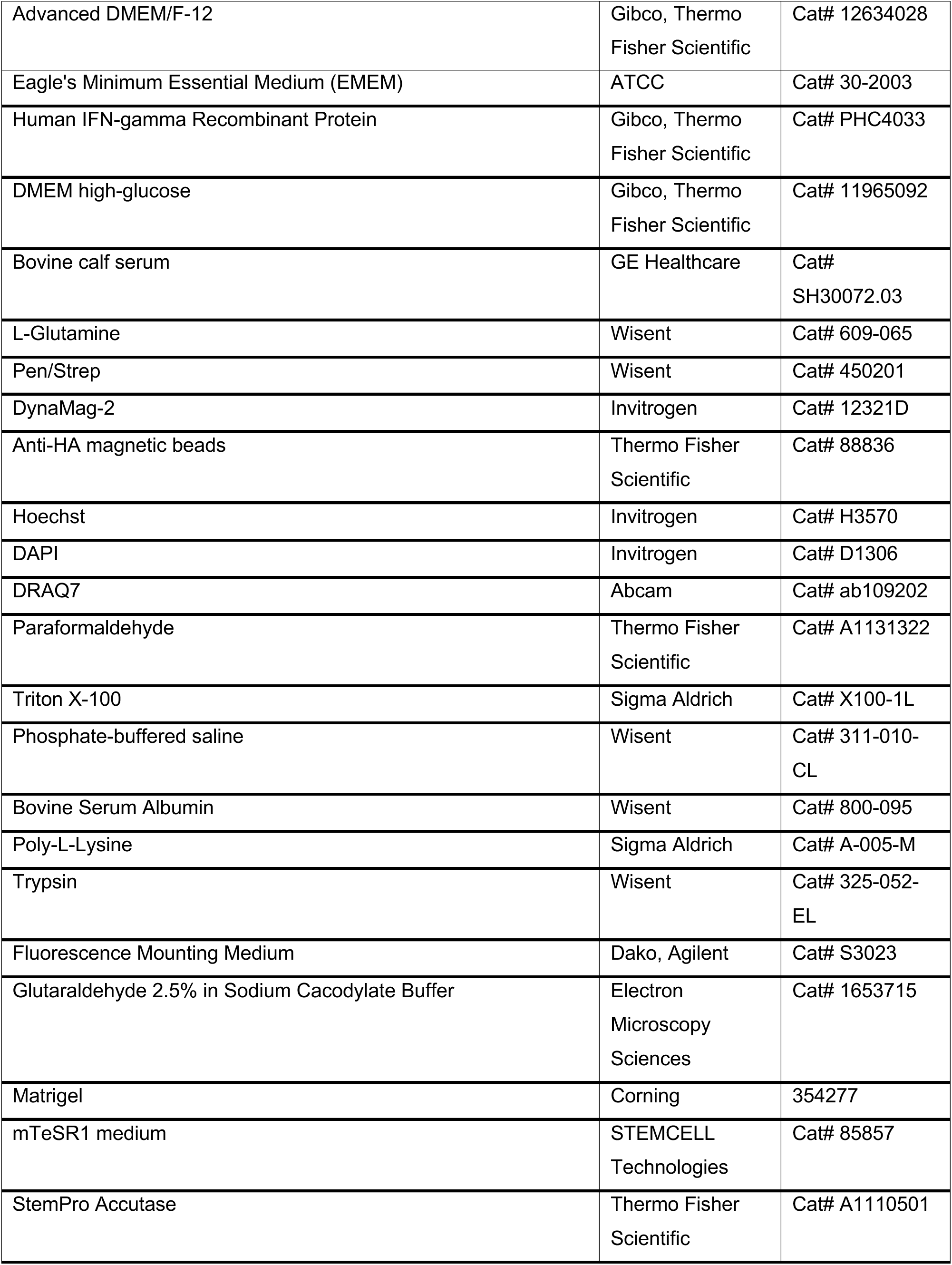
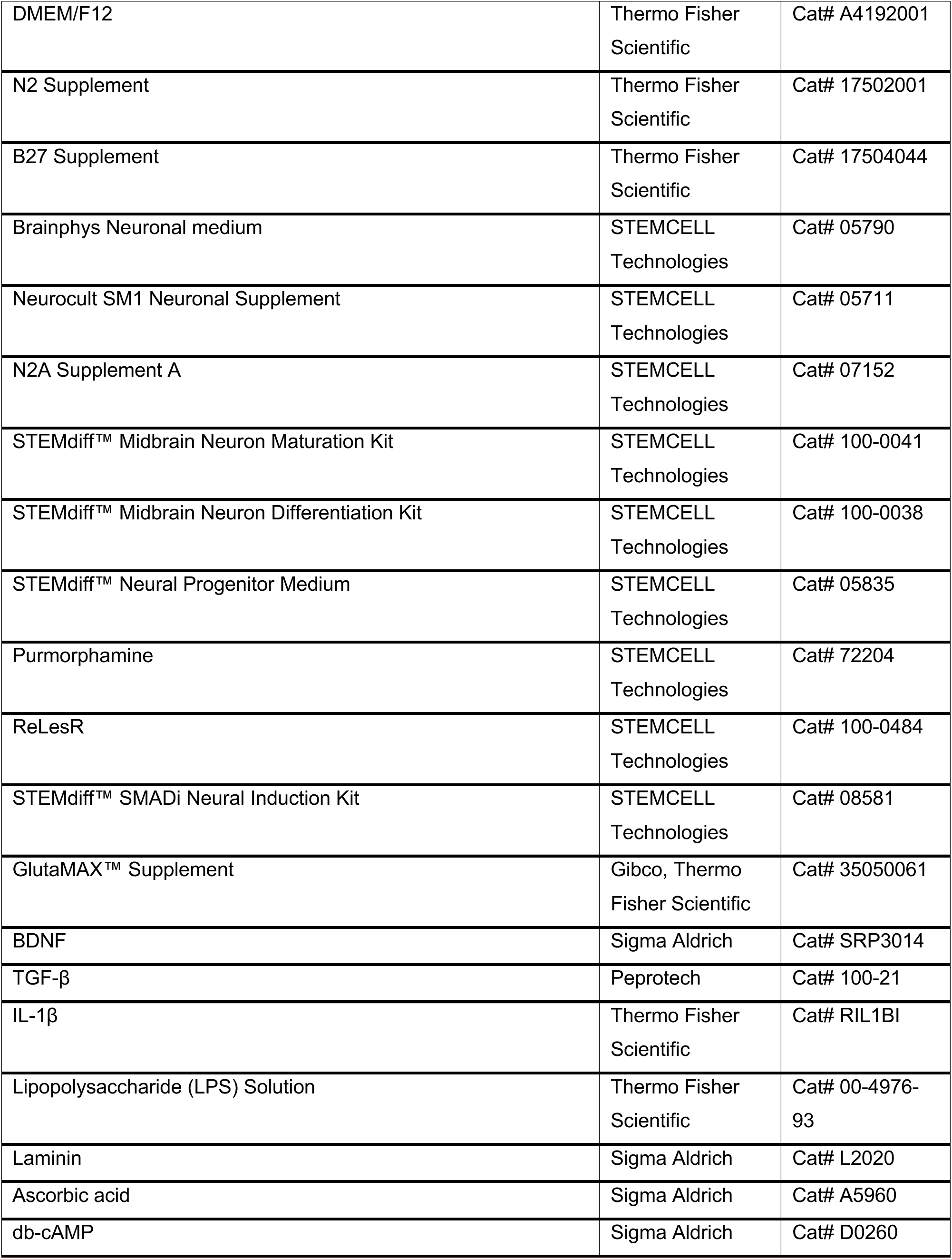
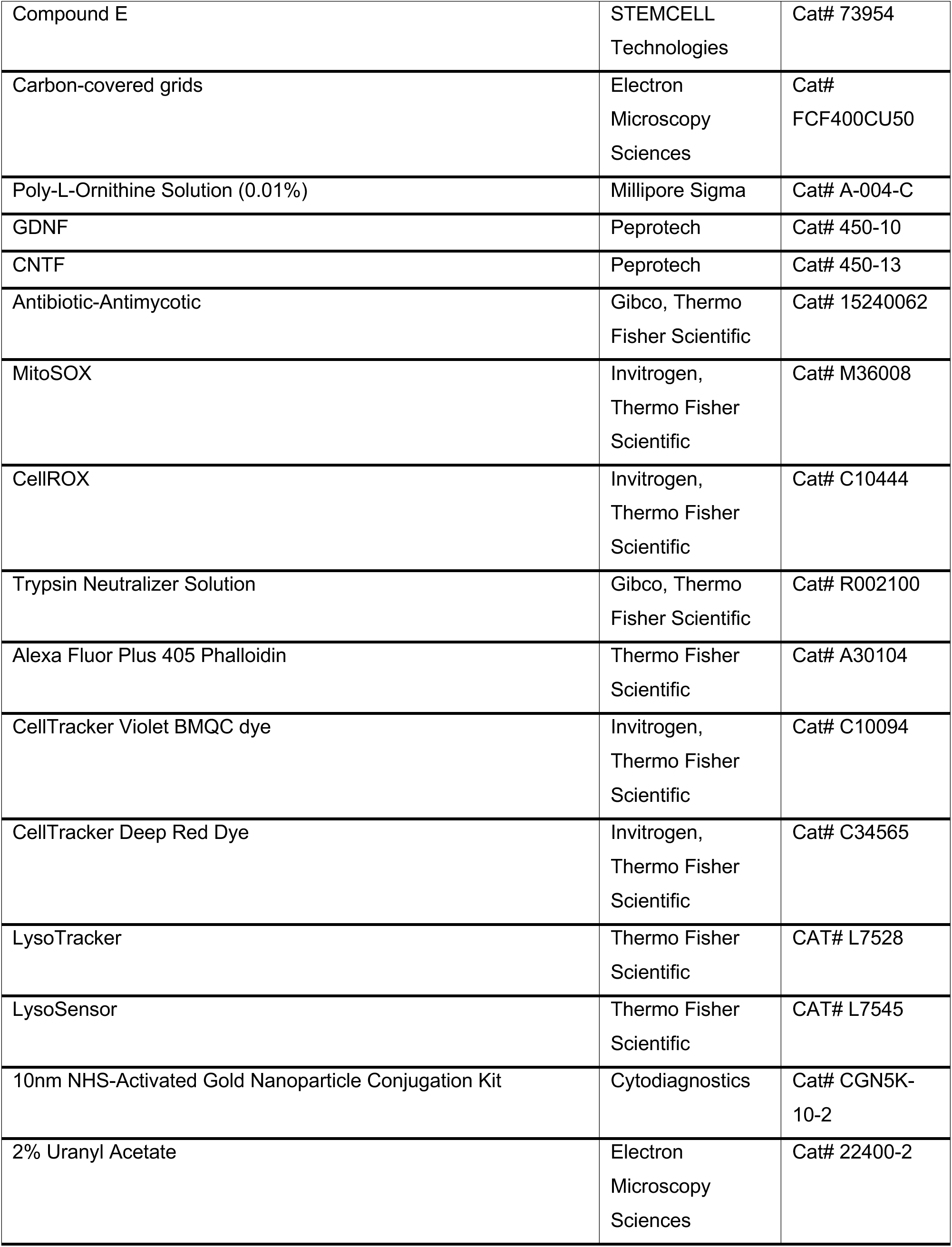
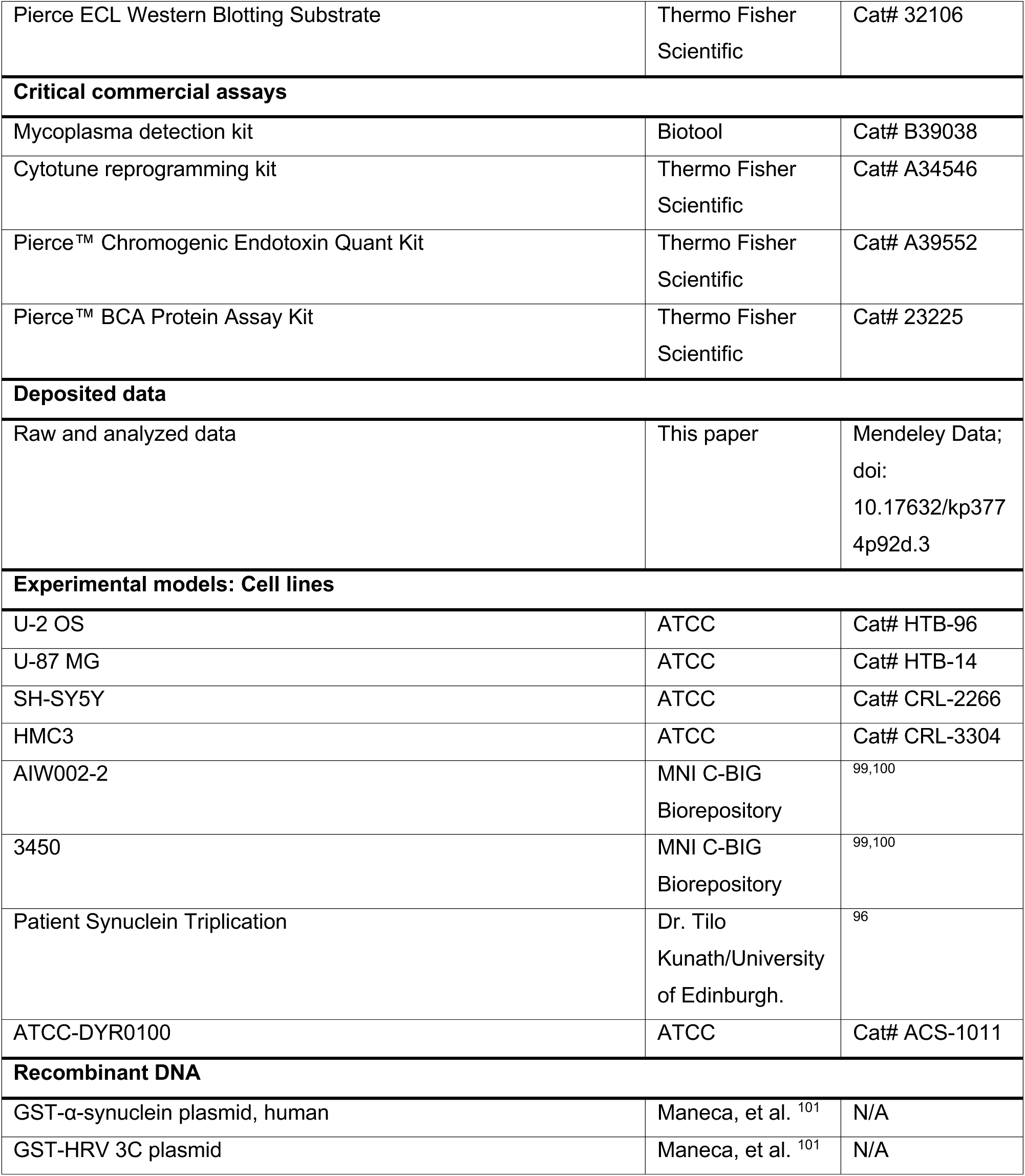
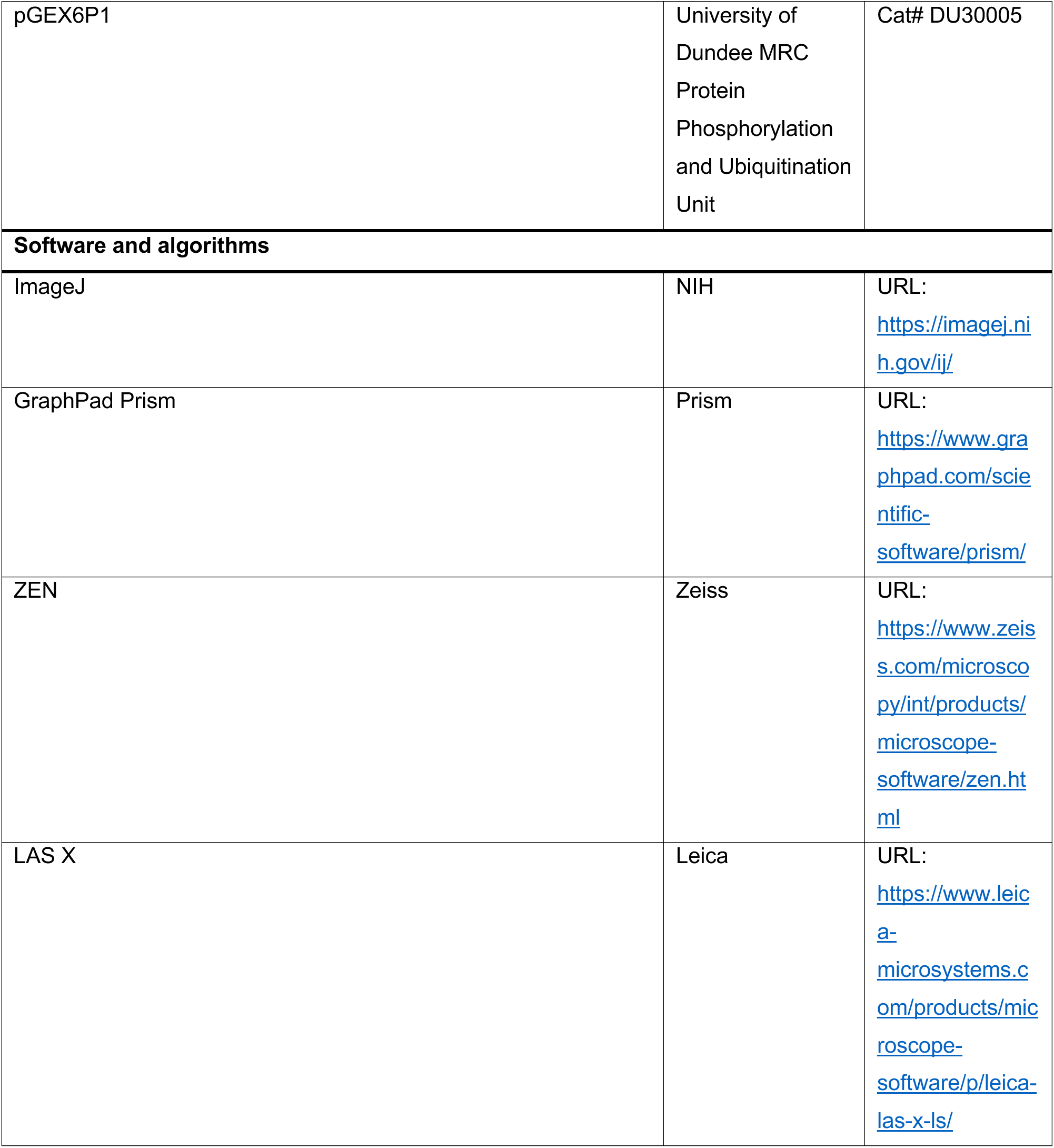
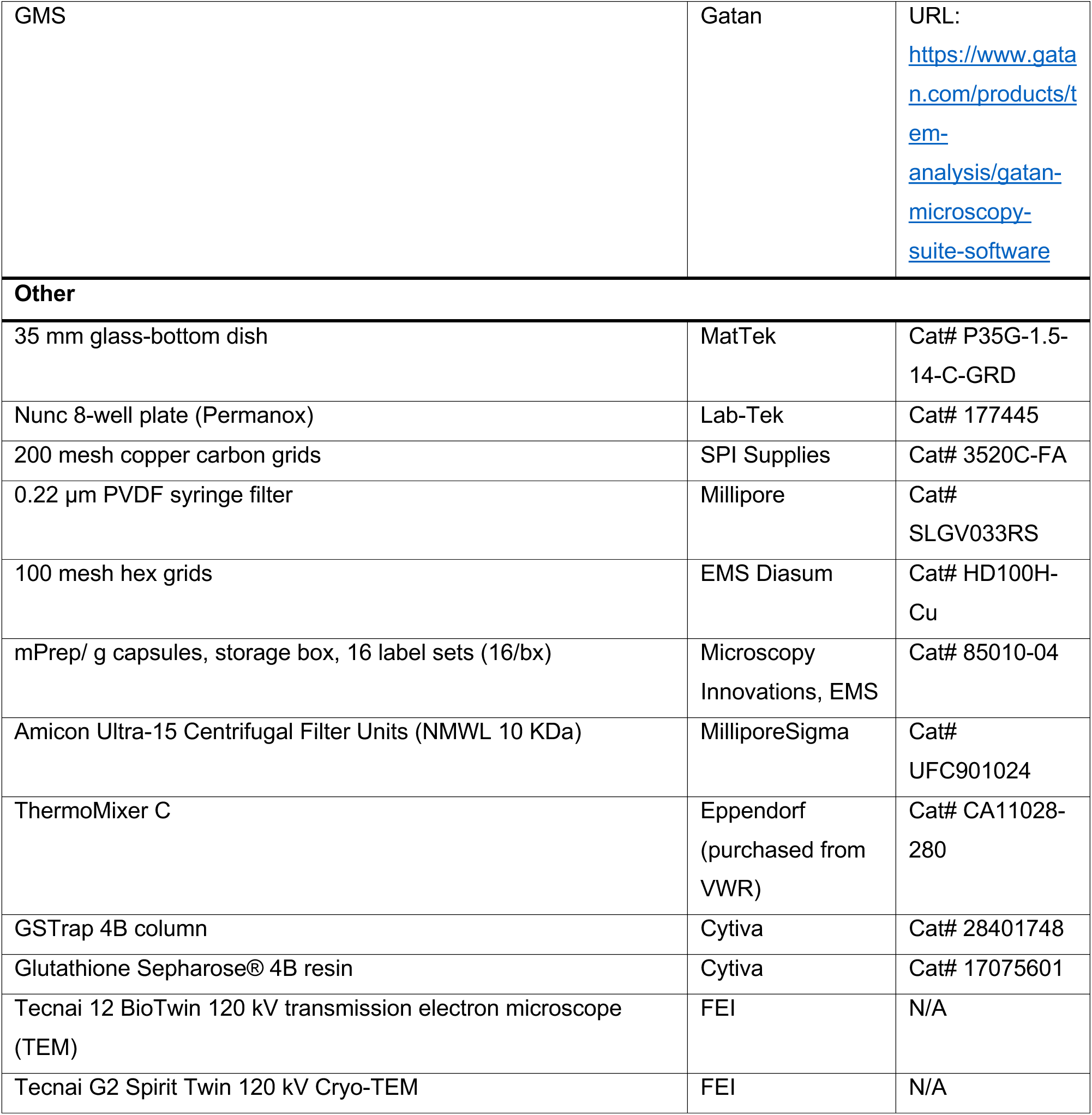
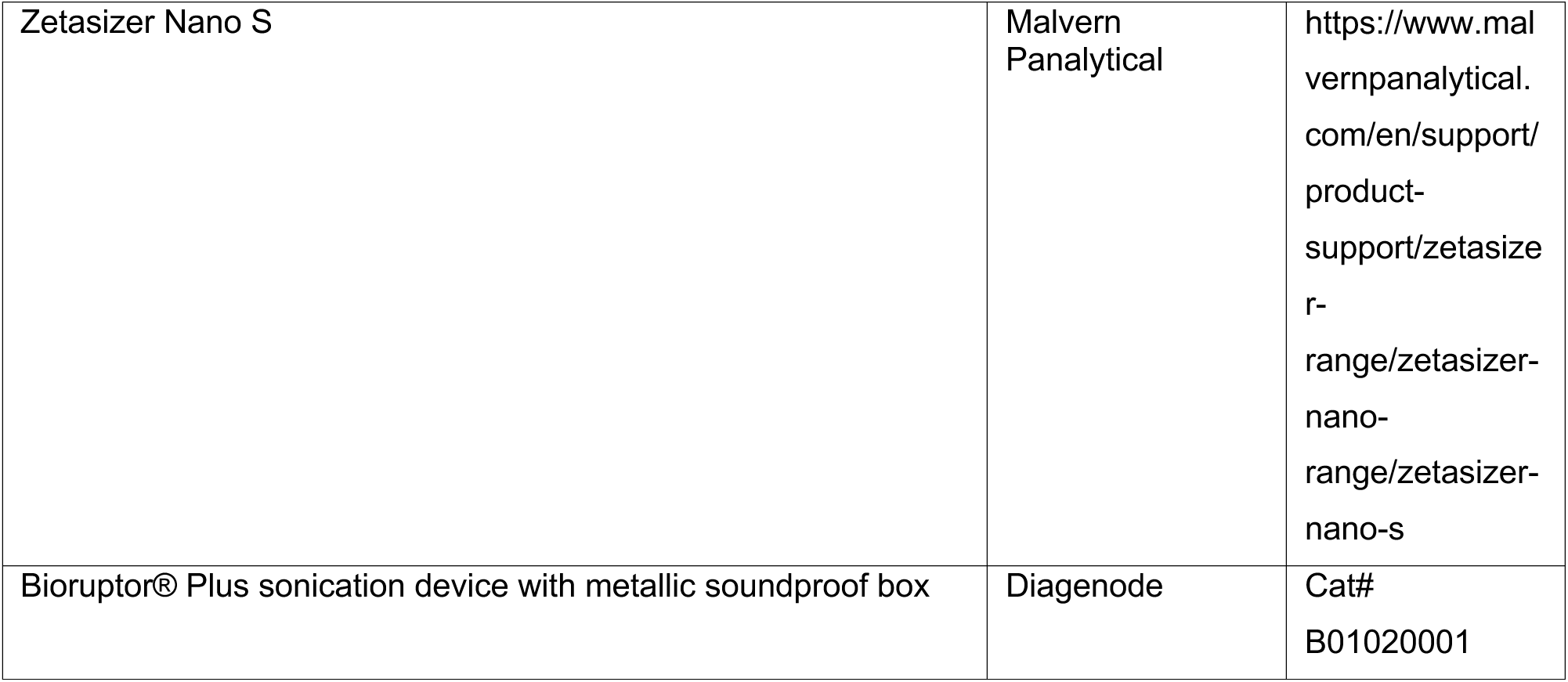
KEY RESOURCE TABLE.

## RESOURCE AVAILABILITY

### Lead Contact

Further information and requests for resources and reagents should be directed to Dr. Peter Scott McPherson. Further information regarding experimental protocols and procedures should be directed to Armin Bayati.

### Materials Availability

This study did not generate new unique reagents.

### Data and Code Availability

- All the raw data, along with statistical calculations used in this paper have been deposited at Mendeley Data. DOIs are listed in the key resources table.
- This paper does not report original code.
- Any additional information required to reanalyze the data reported in this paper is available from the lead contact upon request.

## EXPERIMENTAL MODEL AND STUDY PARTICIPANT DETAILS

### Cell Lines

U2OS, U87, SH-SY5Y, and HMC3 were obtained from American Type Culture Collection (HTB-96, HTB-14, CRL-2266, CRL-3304, respectively). For studies with iPSCs, we used the lines AIW002-2 and 3450 obtained from the Neuro’s C-BIG Biorepository, and ATCC- DYR0100 obtained from ATCC (ACS-1011). AIW002-2 was reprogrammed from peripheral blood mononuclear cells of a healthy donor with the Cytotune reprogramming kit (Thermo Fisher Scientific, A34546). 3450 was also generated from peripheral blood mononuclear cells, but episomal reprogramming was used as described by Wen et al. ^102^. The process of reprogramming and quality control profiling for these iPSCs was outlined in a previous study ^100^. The use of iPSCs in this project is approved by the McGill University Health Centre Research Ethics Board (DURCAN_IPSC / 2019-5374). The *SNCA* lines (originally named AST23, AST23-2KO, AST23-4KO) were provided and generated by Dr. Tilo Kunath from The University of Edinburgh.

Cell lines were routinely checked for mycoplasma contamination using the mycoplasma detection kit (Biotool, B39038).

### Production, Characterization, and Nano-Gold Labeling of PFFs

Production, characterization, and conjugation with nanogold beads of recombinant α-syn monomers and PFF were previously described ^101,103^.

Purification of α-syn was described previously by Feller et al. ^104^. In brief, the pGEX-6P-1 plasmid (University of Dundee MRC Protein Phosphorylation and Ubiquitination Unit, DU30005) cloned with full-length human wild-type α-syn was transformed to BL-21(DE3) *E.coli.* (New England Biolabs, C2527H) and over-expressed as GST-tagged α-syn in the presence of inducer IPTG. The protein was isolated by affinity binding to Glutathione Sepharose^®^ 4B resin (GE Healthcare, 17075601). GST tag was then cleaved off with GST-HRV 3C protease, and α-syn was purified by passing through a GSTrap 4B column (GE Healthcare, 28401748). The purified α-syn was evaluated to be of homogeneity on SDS-PAGE, adjusted to a final concentration of 5 mg/mL with PBS (pH 7.4), and filter-sterilized througth a 0.22-μm PVDF syringe filter (MilliporeSigma, SLGV033RS). The endotoxin level was less than 0.05 EU per mg of α-syn as assayed with a chromogenic endotoxin quant kit by following the manufacturer’s instruction (Thermo Fisher Scientific, A39552). Finally, α-syn was aliquoted and stored at -80°C.

PFF was prepared by shaking 0.5-mL of purified α-syn held in a 1.5-mL microtube on a ThermoMixer (VWR, CA11028-280) at 37°C and 1000 rpm for 5 days. Then PFF was sonicated at least 40 cycles of 30-sec on/30-sec off using a Bioruptor^®^ Pico sonication unit (Diagenode, B01020001). Samples (∼20 μL) were reserved for PFF quality control by thioflavin T assay, dynamic light scattering (using Zetasizer Nano S, Malvern, DLS assay) and TEM imaging as described ^104^.

PFFs were characterized using a negative staining protocol ^103^. PFFs were added to 200 mesh copper carbon grids (SPI Supplies, 3520C-FA), fixed with 4% PFA (Thermo Fisher Scientific, A1131322) for 1 min and stained with 2% uranyl acetate (Electron Microscopy Sciences, 22400-2) for 1 min. PFFs were visualized using a transmission electron microscope (FEI Tecnai 12 Bio Twin 120kV TEM) coupled to an AMT XR80C CCD Camera and analyzed with Fiji-ImageJ1.5 and GraphPad Prism 9 software.

Characterized PFF were conjugated with 10 nm gold beads (CytoDiagnostics, CGN5K- 10-2), immediately before experimental use, as described by Bayati et al. ^105^.

### Differentiation and Culture of AIW002-02 and 3450 DA Neurons

A previously reported differentiation protocol was used to generate DA neurons from two different iPSC cell lines: AIW002-02 (MNI, C-BIG) and 3450 (MNI, C-BIG) ^106^. hiPSCs were plated onto Matrigel (Corning, 354277)-coated plates in mTeSR1 medium (STEMCELL Technologies, 85857). The culture medium was changed daily until the cells reached ∼80% confluency. The cells were then passaged, frozen, or differentiated. A previously described protocol was used to generate ventral midbrain dopaminergic neural progenitor cells ^106^. Dopaminergic neural progenitor cells were dissociated with StemPro Accutase Cell Dissociation Reagent (Thermo Fisher Scientific, A1110501) into single-cell suspensions. Plating on coverslips was done using neural progenitor plating medium (DMEM/F12 supplemented with N2, B27 supplement from Thermo Fisher Scientific, A4192001, 17502001 and 17504044, respectively). To further differentiate into dopaminergic neurons, neural progenitor medium was switched to dopaminergic neural differentiation medium (Brainphys Neuronal medium from STEMCELL Technologies, 05790) supplemented with N2A Supplement A (STEMCELL Technologies, 07152), Neurocult SM1 Neuronal Supplement (STEMCELL Technologies, 05711), BDNF at 20 ng/mL (MilliporeSigma, SRP3014), GDNF at 20 ng/mL (Peprotech, 450-10), Compound E at 0.1 μM (STEMCELL Technologies, 73954), db-cAMP at 0.5 mM (MilliporeSigma, D0260), Ascorbic acid at 200 μM (MilliporeSigma A5960) and laminin at 1 μg/mL (MilliporeSigma, L2020).

### Differentiation and Culture of DYR0100 DA NPCs

The monolayer method was used for the differentiation of DYR0100 iPSCs into DA NPCs as described by Chen *et al*. ^107^. Frozen DYR0100 NPCs were thawed and allowed to recover for one week prior to seeding for downstream assays.

### Generation and Differentiation of NGN2-induced Neurons

AIW002-02 iPSCs expressing doxycycline-inducible *NGN2* were generated as described previously ^108^. Briefly, the parental AIW002-02 iPSC line was split with ReLesR (STEMCELL Technologies, 100-0484) and seeded at a density of 135,000 cells per well in a 6-well plate in mTesR1 media plus Y27632 one day prior to simultaneous transduction with separate lentiviruses encoding Ngn2 and rtTA. The media was changed to mTesR1 only 1 h prior to transduction at a dilution factor of 1/100 for each virus. This resulted in a MOI of one. The transduction was carried out in mTesR1 media plus 4ug/ml of polybrene over a period of 24 h prior to puromycin selection at 1ug/ml for a total of 48 h. Throughout the transduction, selection and iPSC expansion steps, the media was changed daily. The AIW002-02 Ngn2 rtTA line was tested for pluripotency by immunofluorescence for Nanog, Tra-1-80, Oct3/4 and SSEA (positive for all markers, data not shown) and for mycoplasma (negative, data not shown). Subsequently, AIW002- 02 iPSCs were differentiated into Ngn2 neurons based on a protocol adapted from Zhang *et al.*^109^ and Meijer *et al.*^110^ and characterized by immunofluorescence for neuronal markers (MAP2 and PSD95) at the differentiation endpoint.

### iPSC Culturing of Forebrain Cortical Neurons

AIW002-02 iPSCs were differentiated into forebrain cortical neurons according to a protocol based on EB formation combined with dual SMAD inhibition (STEMCELL, 08581) ^100^. In summary, AIW002-02 iPSCs were dissociated into single cells and allowed to form EBs on low-attachment plates for one week in DMEM/F12 media supplemented with N2, B27, 10 µM SB431542, and 2 µM DMH1. This was followed by neural rosette formation onto polyornithine- and laminin-coated (MilliporeSigma, A-004-C and L2020, respectively) plates in the same media, which were selected semi-manually after 7 days. The rosettes were dissociated with Gentle Cell Dissociation Reagent for 5 min at room temperature (RT) and cultured on polyornithine- and laminin-coated plates in DMEM/F12 media supplemented with N2 and B27 to generate NPCs. The NPCs were passaged every 5-7 days until day 25. Final differentiation, was carried out in Neurobasal media supplemented with N2, B27, 1 µg/mL laminin, 500 µM db-cAMP, 20 ng/mL BDNF, 20 ng/mL GDNF, 200 µM ascorbic acid, 100 nM Compound E, and 1 ng/mL TGF-β (Peprotech, 100-21).

### *SNCA* Triplication, Double KO, and Quadruple KO iPSCs

*SNCA* lines (originally named AST23, AST23-2KO, AST23-4KO) were generated and provided by Dr. Tilo Kunath from The University of Edinburgh according to methodology described previously ^96,111^. The AST23 line carries a triplication of the *SNCA* gene, while the AST23-2KO line has been corrected by CRISPR/Cas9-mediated deletion of 2 copies of the *SNCA* gene to create the isogenic control for the former. The AST23-4KO line is a complete *SNCA* knockout.

### PFF + IFN-γ treatment Regime

The 14 d PFF and IFN-γ treatment was described for confocal and Western blot analysis in Fig. 1A. For EM, the regime was described in Fig. S1A. The earlier 2 d time point, used in Western blots and quantifications, was described in Fig. S1B. Briefly, differentiated dopaminergic neurons, grown on 15 mm coverslips, were administered PFFs at 1 µg/mL. The cells were incubated for 48 h with PFF, in Brainphys Neuronal media (STEMCELL Technologies, 05790), and then given fresh Brainphys media and incubated for 72 h. Neurons were then given media supplemented with 0.2 µg/mL of IFN-γ (Thermo Fisher Scientific, PHC4033) or PBS control. Cells were incubated for 24 h, and then given fresh media. 6 d neurons were collected at this point. 7 d neurons were collected 24 h after fresh media was given to cells. 10 d neurons were collected 72 h following the collection of 7 d neurons, and so on. 14 d neurons were collected 8 d following fresh media being given to neurons.

For 2 d neurons, PFF was administered for 24 h, followed by the addition of IFN-γ (or PBS for control) to the PFF containing media for 24 h. A graphical representation of the 2 d treatment is shown in Fig. S1B.

The staggered nature of the PFF and IFN-γ treatment was to enable neurons to survive and form inclusions. In earlier iterations of this protocol, PFF and IFN-γ were treated simultaneously. This led to massive (over 70%) neuronal cell death, and left very few cells for imaging and analysis, and very few inclusions to study. By increasing PFF exposure to 48 h, we were able to increase PFF fluorescence in samples at 14 d and ensure nanogold-PFF could be easily found in EM samples. Incubation in fresh media allowed for neuronal recovery and greatly enhanced neuronal survival. IFN-γ treatment was then limited to 24 h to enhance survival while allowing for the formation of PFF-positive inclusions to occur as early as 7 d into our treatment. Overall, our treatment aimed to increase cell survival while creating the environment necessary for the formation of inclusions in DA neurons. All fixed samples were imaged on an SP8 microscope (Leica). The same protocol was used for the administration of IL-1β (Thermo Fisher Scientific, RIL1BI), where IL-1β was administered at 50 ng/mL instead of IFN-γ. This concentration was used previously by Tong et al. ^112^.

### Maintenance of iPSC-derived Neurons

We note here, that if stressed, iPSC-derived neurons can present with structures that are ultrastructurally somewhat similar to the inclusions we have observed. These structures can even appear in control cells, if care is not taken to 1) Incubate iPSCs at the back of the incubator (most stable temperature), 2) Avoid the use of cold media when changing cell media (temperature shock related stress), 3) ensure cells are never dry while changing media, and 4) avoid contamination. All these factors can lead to cellular stress and cell death, which when viewed under an electron microscope, may lead to cells with dark lysosomes and disorganized cytoskeletal elements, which can be mistaken for Lewy bodies. Fortunately, these apoptotic features in cells lack major compaction and organization when compared to LB-like inclusions, which make them easy to differentiate from bonafide inclusions. Apoptotic cells (or remnants of cells) also lack PFF-gold staining and mitochondrial incorporation.

The utmost importance is to check the quality of the cells and the level of neuronal stress in cultured iPSCs in the control conditions before starting these experiments. If cells are regularly present with pyknotic nuclei, contain dark lysosomes, or unclear mitochondrial cristae in control conditions, measures should be taken to improve cell culturing techniques to ensure the most minimal amount of cellular stress. This will lead to less confusion as to what structures are inclusions and what structures are not when looking at their cellular ultrastructure.

### Reactive Oxygen Species Experiment

In order to quantify the amount of reactive oxygen species (ROS) that were forming as a result of our 14 d treatment, DA neurons and forebrain cortical neurons were plated on live imaging chambers and underwent our 14 d treatment in the presence or absence of IFN-γ, along with an IFN-γ only condition and a control condition where no PFF or IFN-γ was administered (PBS was used instead). Cells were then stained with CellROX and MitoSOX (Thermo Fisher Scientific, C10444 and M36008, respectively) for 1 h prior to live imaging. MitoSOX and CellROX fluorescence was then quantified for both cortical and DA neurons, across two independent experiments. Images were captured on an LSM 880 microscope (Zeiss).

### EM Processing

Neurons and cell lines were plated in 8 well chamber slides (Lab-Tek, Nunc, Thermo Fisher Scientific, 177445) and were put through the dual hit treatment. After treatment, the media was removed and cells were fixed with 2.5% glutaraldehyde in 0.1 M sodium cacodylate buffer (Electron Microscopy Sciences, 1653715) supplemented with 2 mM calcium chloride, washed in buffer, and then post-fixed with 1% osmium tetroxide and 1.5% potassium ferrocyanide in sodium cacodylate buffer. Cells were then *en bloc* stained with 4% uranyl acetate (Electron Microscopy Sciences, 22400-2) and dehydrated in ascending ethanol concentration to 100% and infiltrated with a 50:50 mixture of 100% ethanol and Spurr’s resin, followed by pure Spurr’s resin overnight. After a fresh addition of resin, samples were polymerized overnight at 70°C in an oven. Ultrathin sections were cut on a Leica Ultracut E using a 45° diamond knife and collected on 100 mesh copper hex grids (Electron Microscopy Sciences, HD100H-Cu). Some sections were stained with uranyl acetate for 5 min for enhanced membrane staining. Grids were examined with a FEI Tecnai Spirit 120 kV transmission electron microscope and images captured using a Gatan Ultrascan 4000 camera.

### Plasmids and Lentivirus

α-synuclein-HA-tagged adenovirus was obtained from Applied Biological Materials Inc. (44541051). Protocols provided by the manufacturer were followed for transduction.

### Fixation and Antibody Staining

Cells were fixed with 4% paraformaldehyde (PFA; Thermo Fisher Scientific, A1131322) for 10 min at 4°C, washed with PBS and then permeabilized for 30 min with 0.05% Triton X-100 (MilliporeSigma, X100-1L) in PBS (Wisent, 311-010-CL) along with 5% bovine serum albumin (BSA, Wisent, 800-095). Coverslips were then transferred into a wet chamber and incubated for 2 h at RT with a 1:500-1:1000 dilution of the primary antibody diluted in PBS containing 0.01% Triton X-100 and 5% BSA (dilutions varied depending on the antibody). Coverslips were then gently washed twice with PBS, and a 1:1000 dilution of secondary antibody diluted in the same buffer was added and incubated for 2h. Cells were then washed with PBS and, in some situations, stained with 1 µg/ml Hoechst 33342 (Fisher Scientific, H3570) diluted in PBS for 10 min at RT. Coverslips were mounted on a glass slide with Fluorescence Mounting Medium (Dako, Agilent, S3023). Samples were then imaged using a Leica TCS SP8 confocal microscope.

### LB-like Inclusion Extraction (Western Blot and EM)

Neurons undergoing the dual hit treatment regime were collected at day 7 and 10 for the extraction of inclusions. This was done since previous attempts at extraction of inclusions from 14 d samples resulted in very little isolated material, perhaps due to cellular stress and cell death common with that time point. In brief, cells were washed three times in PBS 1x, pH 7.4 and scraped off in a hypotonic lysis buffer (20 mM HEPES, 1 mM EDTA, pH 7.4) supplemented with protease and phosphatase inhibitor cocktail mix. Cells were kept on ice for 30 min then passed 8 times through a 27G needle until homogenized. Homogenates were kept on ice for 30 min and then 10% of the homogenate was transferred into a fresh tube (total homogenate sample), and the remainder was centrifuged at 720 x *g* for 10 min at 4°C using a tabletop centrifuge. The post-nuclear supernatant (PNS sample) was transferred into a fresh tube and the pellet (P sample) containing large LB-like inclusions and nuclei was suspended in lysis buffer. For Western blot, all samples were sonicated for 10 sec at 40% amplitude and lysates were prepared in Laemmli sample buffer (1x) followed by SDS-PAGE. For EM, the pellet was kept at - 80°C until processed.

### Western blot protocol

Cells were washed three times in PBS (pH 7.4) and then scraped off in RIPA buffer: 50 mM TRIS-HCl pH 7.4, 150 mM NaCl, 1mM EDTA, 0.5% sodium deoxycholate, 0.1% SDS (pH 7.4). RIPA lysis buffer was supplemented with a protease and phosphatase inhibitor cocktail mix. Lysates were sonicated for 10 sec at 40% amplitude, kept on a nutating rocker for 30 min at 4°C and centrifuged at 21,000 x *g* for 15 min at 4°C using a tabletop centrifuge. Supernatants were collected in a new tube and protein concentrations were measured using a BCA protein assay kit (Thermo Fisher Scientific, 23225).

Equal amounts of protein from each condition tested were prepared in Laemmli sample buffer prior to SDS-PAGE. Samples were resolved on large gels with the appropriate gradient of polyacrylamide. Proteins were transferred to nitrocellulose membrane and labelled appropriately. For Western blot, membranes were blocked in 5% milk in TBST (20 mM TRIS, 150 mM NaCl, 0.1% Tween20, pH 7.4). Primary antibodies were diluted in 5% BSA in TBST and membranes incubated overnight at 4°C with constant rocking. Membranes were washed three times (10 min each) in TBST and then incubated in the corresponding secondary antibody for 1 h at RT. Membranes were then washed three times in TBST with constant rocking. For signal exposure, membranes were incubated with the Pierce ECL Western Blotting Substrate (Thermo Fisher Scientific, 32106) for 1 min and developed on autoradiography film in a dark room.

All Western blots were done three times, from three separate iPSC frozen batches, prior to being incorporated into the manuscript.

### IFN-γ and PAH Treatment in DA Neurons and Cell Survival

DA NPCs were thawed and cultured on 24 well plates, pre-treated with PO and laminin. NPCs were then matured into neurons using the Midbrain Neuronal Maturation Kit (STEMCELL Technologies, 100-0041) for 14 d. Following differentiation, neurons were treated with varying concentrations of IFN-γ or PAH for 24 h. Cells were then stained with Hoechst 33342 for 20 min at RT, washed with PBS and nuclear fluorescence quantified using Infinite 200 Pro Tecan plate reader. Six measurements for each concentration of PAH were conducted and used in the quantification. Quantification was done by setting the control condition’s nuclear fluorescence to 100% and dividing all subsequent cell counts by the control, resulting in the percentage of cell count compared to control.

### PAH Treatment in DA Neurons for Western Blot Experiment

DA NPCs were thawed and cultured in 6-well plates in STEMdiff™ Neural Progenitor Medium (STEMCELL Technologies, 05835). The cells were then differentiated into neurons using Midbrain Neuron Maturation Kit (STEMCELL Technologies, 100-0041) for 14 d. Following differentiation, 6 wells were established as various controls: (1) null condition with PBS and DMSO treatment, (2) IFN-γ only condition in which neurons were treated with IFN-γ for 48 h and then incubated in fresh Brainphys Neuronal media (STEMCELL, 05790) and maintained for 14 d, (3) PFF only condition in which neurons were only treated with PFF for 48 h and then incubated in fresh media and maintained for 14 d, (4) PAH only condition in which cells were treated with 10 µM PAH for 24 h, given fresh media and maintained for 14 d, (5) PFF and PAH condition in which neurons were incubated with PFF for 48 h given fresh media for 72 h, and then treated with PAH for 24 h, (6) IFN-γ and PAH condition, in which neurons were treated with PBS (control for PFF) for 48 h, given fresh media, incubated with IFN-γ for 24 h, given fresh media for 24 h, and then given 0.025 mM of PAH for 24 h, then given fresh media and maintained until 14 d. 5 wells were then designated as experimental conditions: (7) cells were given PFF for 48 h, fresh media for 72 h, IFN-γ for 24 h, 24 h of fresh media, 24 h of media supplemented with 0.025 mM PAH, following by 72 h of incubation with fresh media before cell collection, (8) same as condition 7, except that 96 h elapsed between IFN-γ treatment and PAH treatment, (9) same as condition 7 except that 8 d elapsed between IFN-γ and PAH treatment. (10) same as condition 7 except that no PAH treatment was done and samples were maintained until 14 d, (11) same as condition 10, except that IFN-γ treatment was done for 48 h, followed by incubation in fresh media until 14 d. All cells were fixed 72 h after the addition of PAH.

### Endogenous α-syn expression in KO and WT *SNCA* DA Neurons

KO and WT *SNCA* DA NPCs from Mohamed *et al.* ^96^ were differentiated into neurons as described above. Neurons were then treated for 48h with α-syn-HA adenovirus (Applied Biological Materials, 445410520200) at concentration of 2 x 10^5^ IFU/PFU (infectious unit)/ml. Neurons were then collected, lysed, and processed for Western blotting.

For imaging experiments, neurons grown on coverslips, then underwent the protocol described above and in Fig. S10, in which they are given 72 h of rest in fresh media, followed by administration of IFN-γ for 24 h, followed by administration of fresh media. Neurons were then fixed at various timepoints, stained with HA and phospho-α-syn antibodies.

A 2 d shortened treatment was also conducted to serve as controls for the longer timepoint experiments (Fig. S1B).

### Microglia Media Analysis

HMC3 cells were grown on 150 mm plates using EMEM (ATCC, 30-2003) supplemented with 10% bovine calf serum (GE Healthcare, SH30072.03) and Pen/Strep (Wisent, 450201). Once cells reached 80% confluency, cells were given PBS, LPS, PFF, and LPS + PFF. LPS (Thermo Fisher Scientific, 00-4976-93) was administered at 10 µg/mL concentration, while PFF was administered at 1 µg/mL concentration. Cells were incubated for 24 h in the treatment media then trypsin washed three times to remove PFF on the cell surface. Cells were passaged onto new plates and incubated in complete media for 24 h. Cells were washed four times with PBS 1x and incubated in serum-free media (EMEM + Pen/Strep) for 48 h. Media from each condition were collected, centrifuged for 10 min at 500 x *g* to eliminate any cells or large contaminants, then for 10 min at 4000 x *g* to eliminate any small contaminants. Media were concentrated by centrifuging at 4000 x *g* for 30min using Amicon Ultra-15 Centrifugal Filter Units with a membrane NMWL of 10kDa (MilliporeSigma, UFC901024). Concentrations of protein in the media were quantified and equal amounts of protein from each condition were resolved by SDS-PAGE followed by the corresponding Western blot to detect each protein.

### LysoTracker and LysoSensor Experiment

To assess the lysosomal pH and health of neurons undergoing the 14 d dual hit treatment regime, LysoTracker (Thermo Fisher Scientific, L7528) and LysoSensor (Thermo Fisher Scientific, L7545) were used. LysoTracker was added to cells 15 min prior to imaging, while LysoSensor was added 5 min prior to imaging. Neurons were plated on 35 mm glass bottomed dishes (MatTek, Cat# P35G-1.5-14-C-GRD). Cells were imaged using LSM 880 (Zeiss), equipped with a live imaging chamber.

LysoTracker lysosomal count was done by taking thresholded images (50% threshold) and loading them into ImageJ (NIH) and using the “3D Object Counter” function. The software then outputted the number of lysosomes in each image. Cell count was done by taking the Cell tracker (Thermo Fisher Scientific, C10094) channel and loading it into ImageJ and using the “Find Maxima” function to count the number of cells. The number of lysosomes was then divided by the number of cells to attain the “Number of Lysosomes per Cell” quantity. This was averaged across three independent experiments, with 3 images from each experiment being taken.

### Co-culture experiments

HMC3 cells were grown in EMEM media supplemented with 10% of bovine calf serum and 1x of Gibco Antibiotic and Antimycotic (Thermo Fisher Scientific, 15240062). Glioblastoma, U87 cell line were grown in DMEM media (Gibco, Thermo Fisher Scientific, 11965092) supplemented with 10% of bovine calf serum, 2 mM of L-Glutamine (Wisent, 609065), and 100 IU Penicillin 100 μg/ml Streptomycin. Cells were grown and expanded on 150 mm plates and were passaged onto 6-well plates prior to the beginning of the experiment. HMC3 (microglia) were treated with LPS at 10 µg/mL concentration or given PBS of equal volume (for control). Cells were also given 5 µL of CellTracker Violet dye (Thermo Fisher Scientific, C10094). 24 h following LPS treatment, cells were trypsinized, washed and pelleted three times, and added to coverslips containing differentiated DA neurons previously exposed to PFFs (5.0 x 10^4^ cells per coverslip). Microglia were incubated with DA neurons, in Neurobasal media for 48 h prior to fixation. Samples were then stained with β-III tubulin and prepared for microscopy.

### Cell Line Panel Experiments

U87 and U2OS cells were grown in 500 mL DMEM media, supplemented with 10% bovine calf serum, 2 mM of L-Glutamine, and 100 IU Penicillin 100 μg/ml Streptomycin. SH-SY5Y cells were grown in 500 mL of Advanced DMEM1/F-12 (Gibco, Thermo Fisher Scientific, 12634028) supplemented with 10% bovine calf serum, 2 mM of L-Glutamine, and Antibiotic-Antimycotic at 1x. Cells were grown and expanded on 150 mm plates and were passaged onto 6-well plates. Cells were treated with PFF for 48 h, trypsin washed on ice for 30 s, trypsinized, plated onto new 6-well plates and incubated for 48 h. Cells were then plated at low confluency onto poly-L-lysine (MilliporeSigma, A-005-M)-treated coverslips and incubated in fresh media for another 24 h. Cells were then treated with IFN-γ (or PBS for the PFF-only condition), for 24 h. Cells were then given fresh media, and incubated for 8 days. The cells in the PFF-only condition had to be passaged once more 96 h later, due to overconfluency. Cells were then fixed, stained with LAMP1 (Cell Signaling Technology, 9091S) and Phalloidin (Thermo Fisher Scientific, A30104) and prepared for imaging.

### Graphical Abstract and Graphical Representation of treatment regimes

Graphical abstract, graphical representation of treatments, and models were created using Biorender.com.

### Cell and Inclusion Count

An automated approach was taken for the cell and inclusion counts as described by Labno ^113^. Cells and inclusions were counted by loading single-channel images into ImageJ (NIH) and performing the “analyze particles” function, following thresholding of images. The same threshold was used for all the conditions within each experiment (thresholds were different for different experiments). For cell count, the nuclear staining channel was used, and for inclusions the PFF channel was used. For inclusions, particles smaller than 2.0 µm in diameter were not included in our count.

### Inclusion Size

Using the “analyze particles” function in ImageJ, thresholded single channel images were loaded into ImageJ and particles of high circularity and 0-infinity in size were selected. The “show” and “outline” options were also chosen. From the outlines produced by ImageJ, the diameters of the inclusions were then measured manually. For the images in which inclusion size was measured, a high threshold was set to exclude most lysosomal-PFF but include smaller puncta with high intensity levels. This was done so that a measure of the more intensely fluorescent puncta was incorporated in our size calculation. Although some puncta measured were under 2.0 µm in diameter, which is below our inclusion definition set above, we believe that this provides a better representation of where most PFFs are located, and what sort of structures they are localized to. In samples where PFFs are less aggregated, the brightest puncta were smaller, whereas in 14 d samples, more of the brightest PFF puncta will be localized to inclusions and therefore the PFF puncta will be of larger size. In short, we wanted to measure the size of the PFF puncta with the highest fluorescence intensities.

### Lysosomal Count, Lysosomal Size, Fluorescence Intensity

To count the number of lysosomes per cell, results shown in Fig. S9, the MorphoLibJ plugin ^114^ was used, using fluorescent microscopy data. Single plane, single cell, single channel images previously thresholded were then loaded into ImageJ, and the number of puncta was calculated according to the methods outlined by Peters et al. ^115^. Lysosomal size was calculated using electron micrographs, and the lysosomes were measured manually using GMS 3.0 (Gatan). Fluorescence intensity was calculated using LAS X (Leica) software, where channel intensities are quantified using the “Quantification” function.

### Quantification Analysis

Measurements, including cell/inclusion count, inclusion size and puncta number, were all inputted into Graphpad Prism 9.0. Quantification data was statistically analyzed, using Student’s t-test when only two conditions were being compared, one-way ANOVA for multiple conditions (along with post hoc Tukey’s test), and two-way ANOVA when comparing multiple independent variables and multiple dependent variables (multiple comparisons was done within each column across different rows). All raw data and statistical analyses (not multiple comparisons analyses) have been deposited into Mendeley data online repository.

## Supplementary Table 1.

**Supplementary Figure 1.**
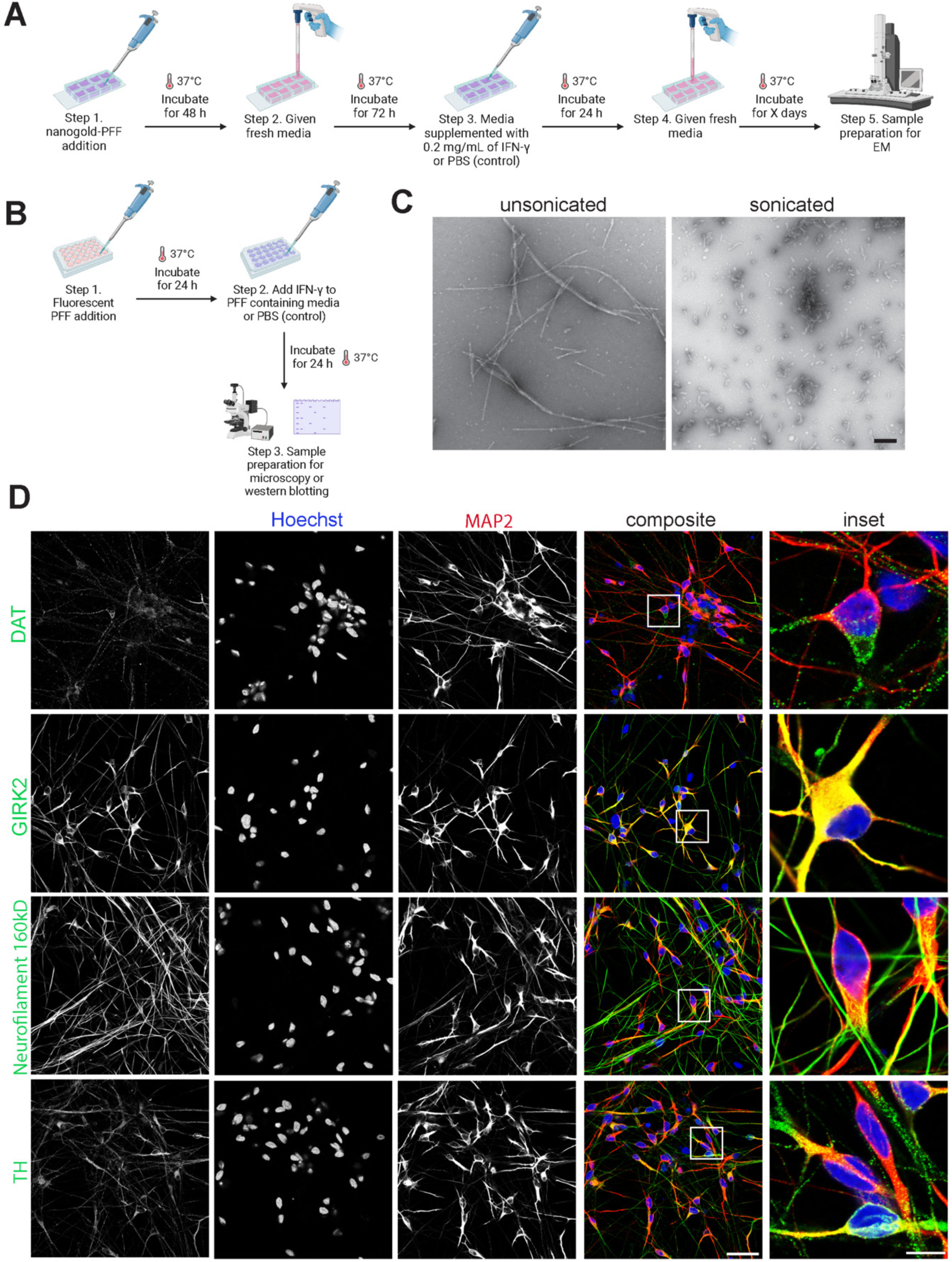
Dual hit treatment, PFF characterization, and neuronal characterization. (**A**) Graphical representation for the dual hit treatment regime used for DA neurons grown and differentiated in 8 well permanox chambers. (**B**) Graphical representation of the shortened dual hit treatment protocol for the 2 d time point. (**C**) Characterization of unsonicated and sonicated α-syn fibrils. Scale bar = 200 nm. (**D**) DA neurons that were differentiated and matured were characterized using the following antibodies: Dopamine Transporter (DAT), GIRK2, Neurofilaments (160kD), and Tyrosine Hydroxylase (TH). Neurons were also positive for MAP2. While DAT and TH confirm the dopaminergic identity of the neurons, GIRK2 specifies the regional specificity of these DA neurons to the substantia nigra. MAP2 and Neurofilaments confirm the neuronal maturity of the neurons.

**Supplementary Figure 2.**
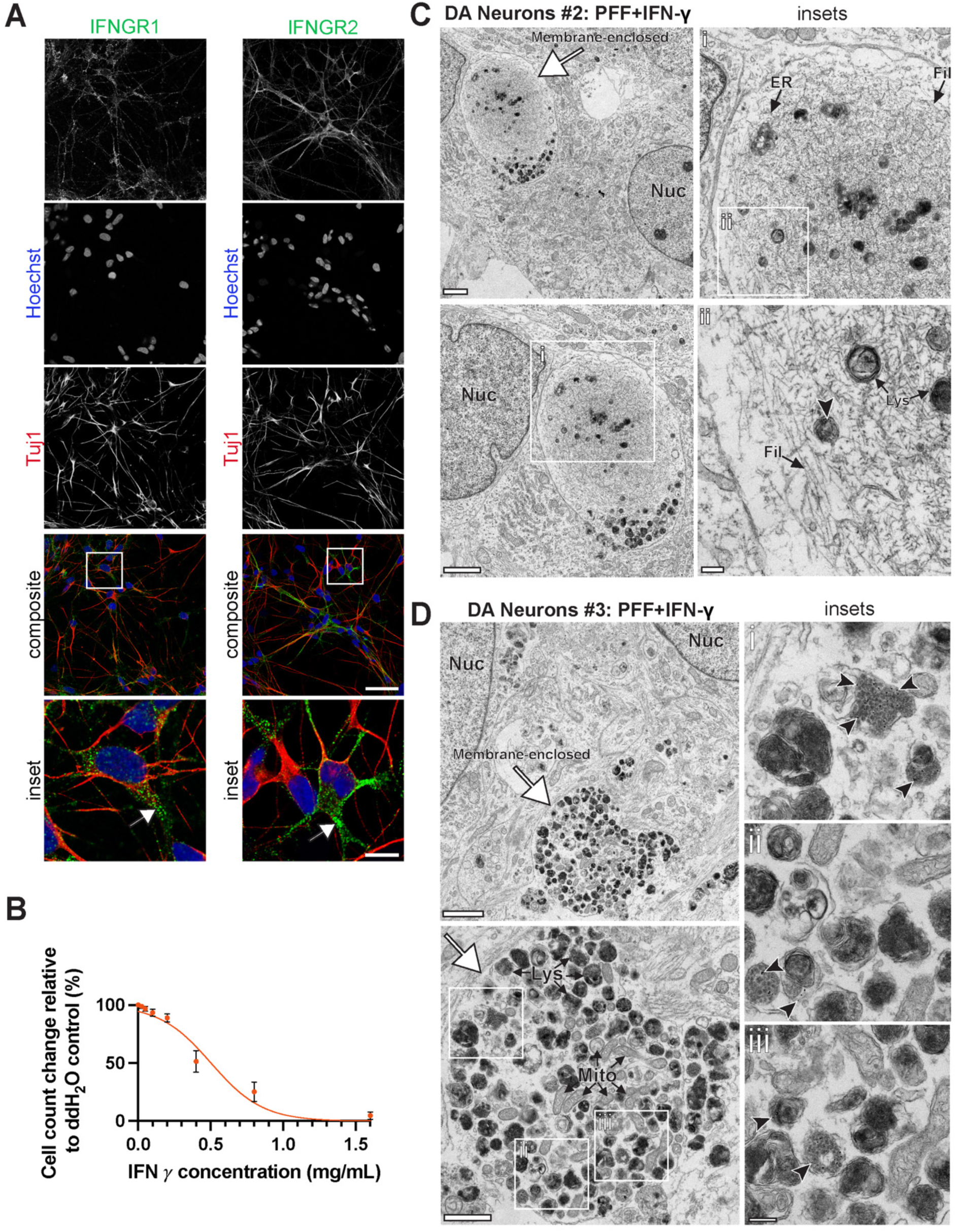
IFN-γ receptor expression, IFN-γ dose-response curve, and inclusions in DA neurons from other iPSC cell lines. (**A**) Antibodies against IFN-γ receptors 1 and 2 were used to illustrate the expression of these receptors in DA neurons. Scale bars = 40 µm and 10 µm for insets. (**B**) The dose-response curve for the concentration of IFN-γ used and the corresponding cell count, using Hoechst to stain nuclei. Cells were counted using plate reader (Tecnai). (**C**) Inclusions found within DA neurons generated from the DYR-0100 line (cell line #2). An exceptionally large inclusion that is mostly filled with filamentous materials, with islands of organelles, can be observed. Scale bars = 2 µm, 1µm and 200 nm, respectively from lowest to highest magnification images. (**D**) A mostly lysosomal- and mitochondrial-filled inclusion generated in 3450 iPSC DA neurons (cell line #3). Scale bar = 2 µm for large field, 1 µm for inclusion, and 200 nm for insets. Lysosomes/autolysosomes (Lys), mitochondria (Mito), filaments (Fil), endoplasmic reticulum (ER), white arrows point to the membrane surrounding the inclusions, and dark arrowheads point to a few examples of nanogold-labeled PFF inside lytic vesicles.

**Supplementary Figure 3.**
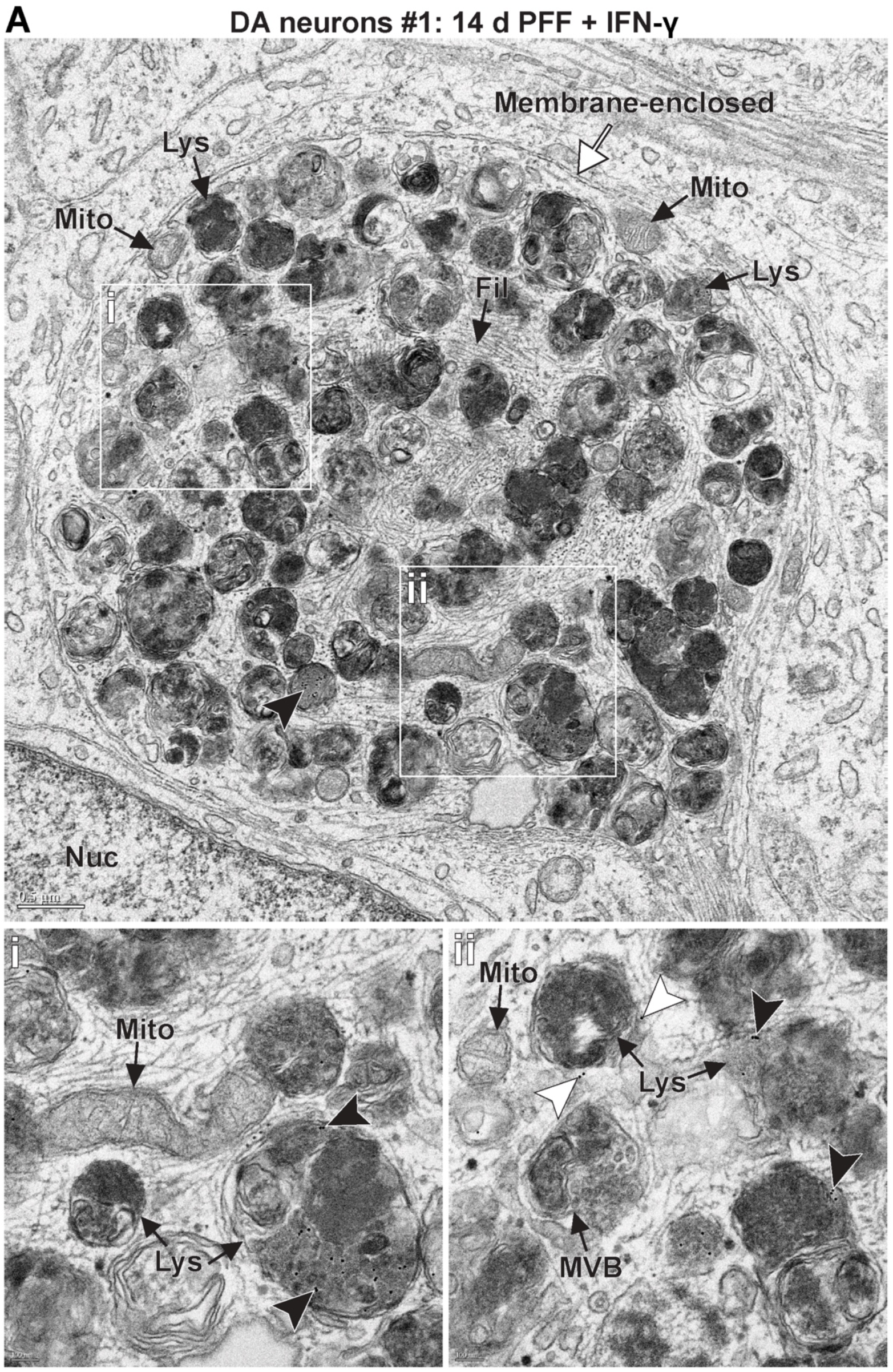
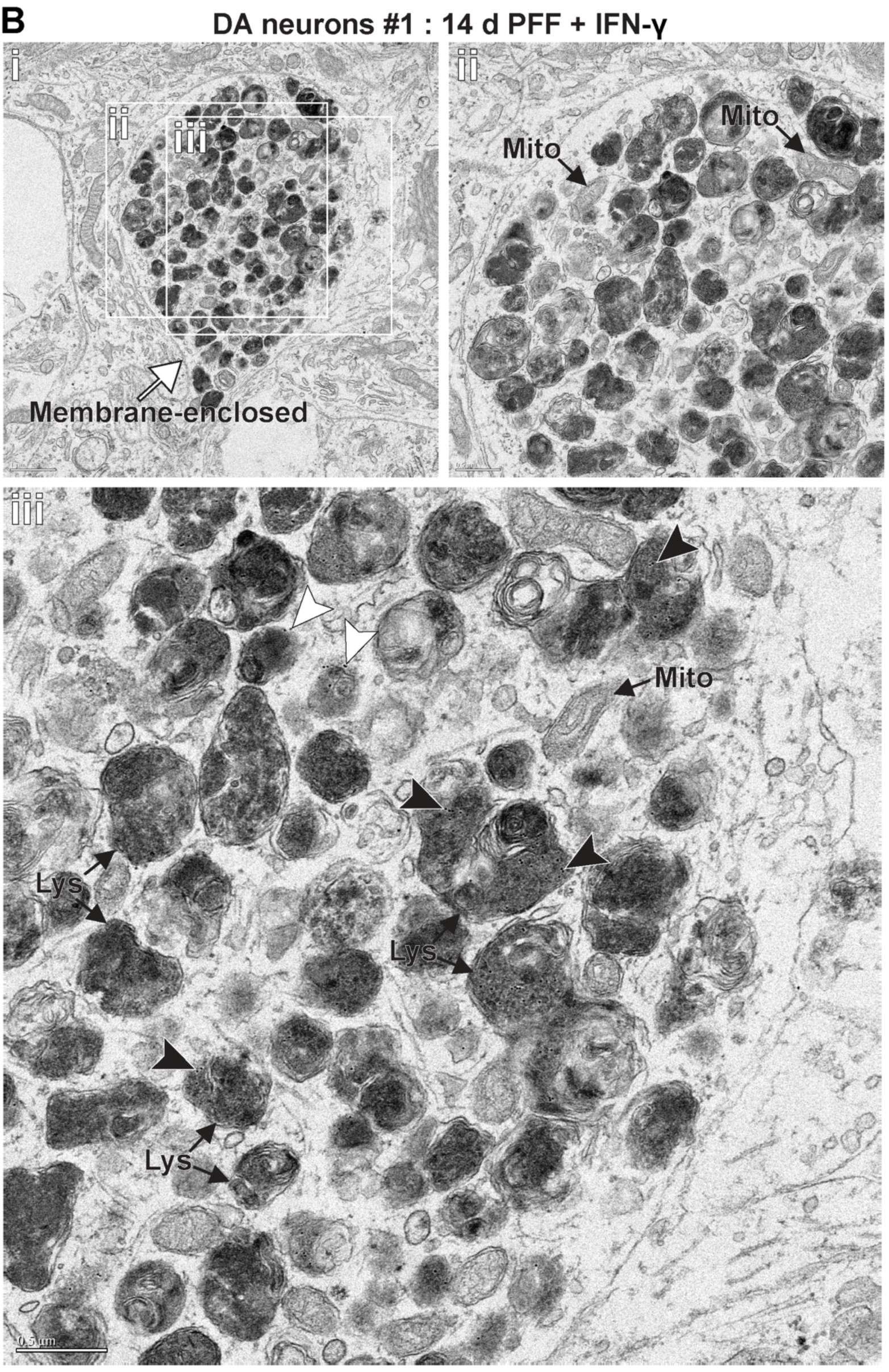
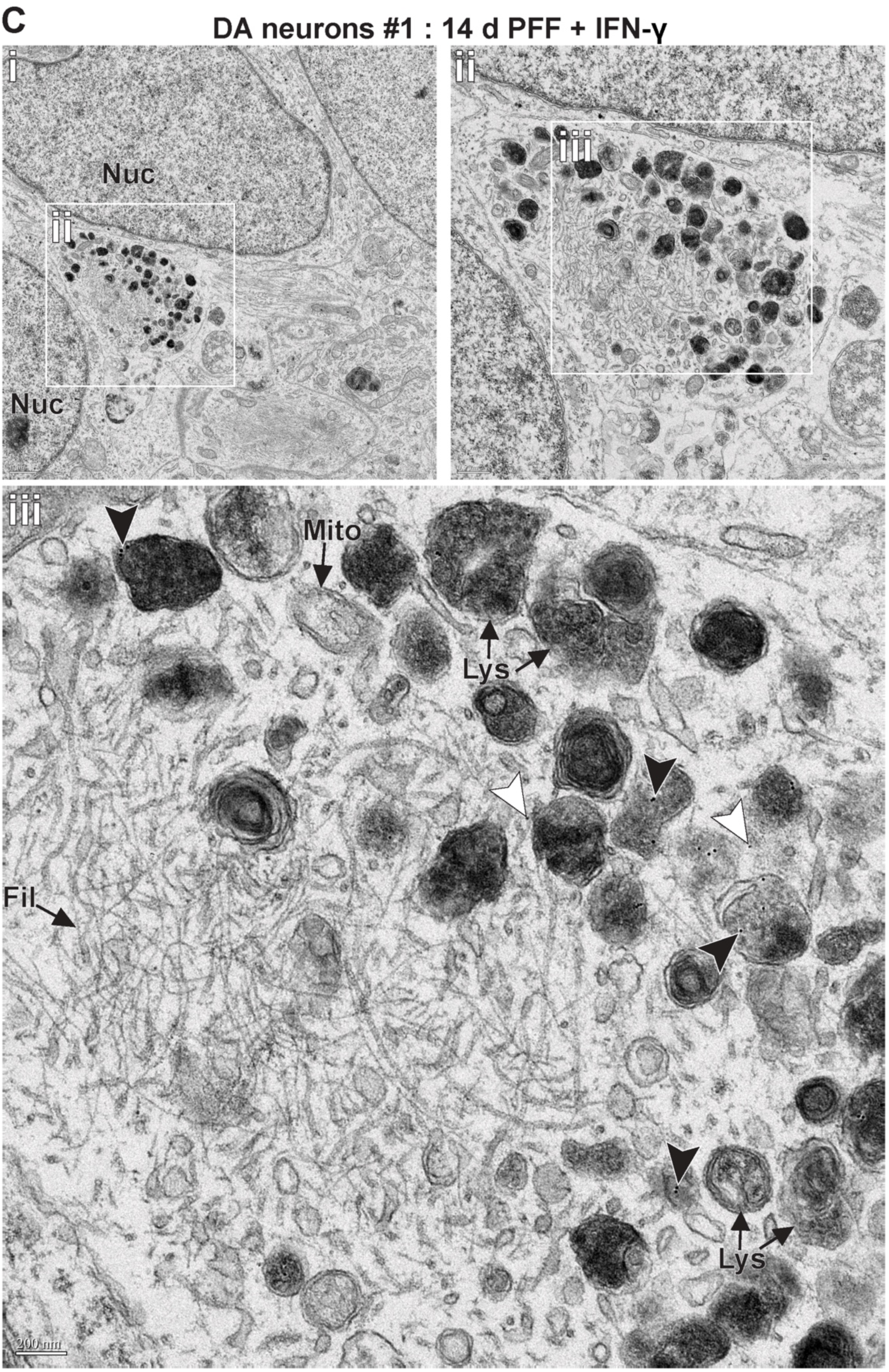
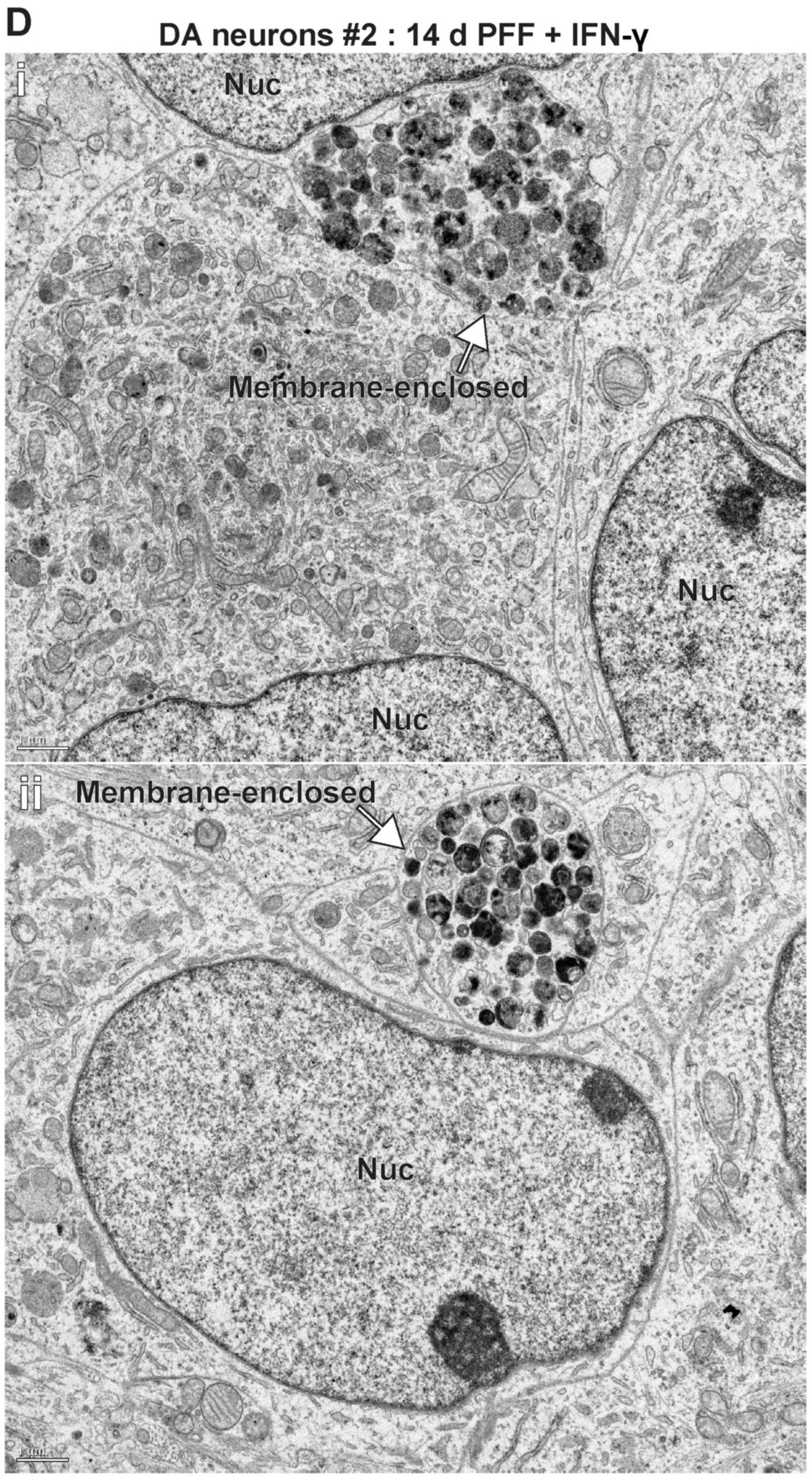
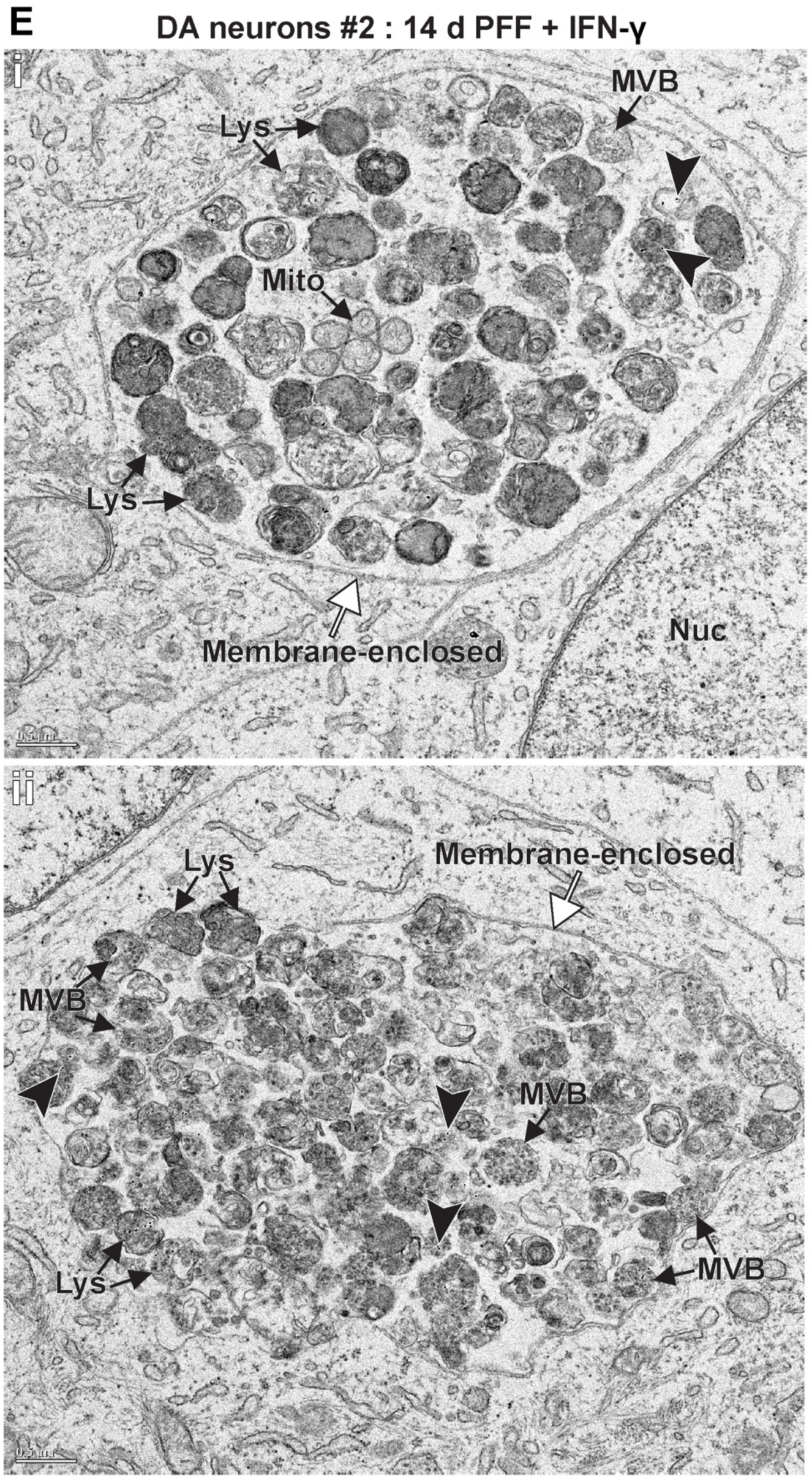
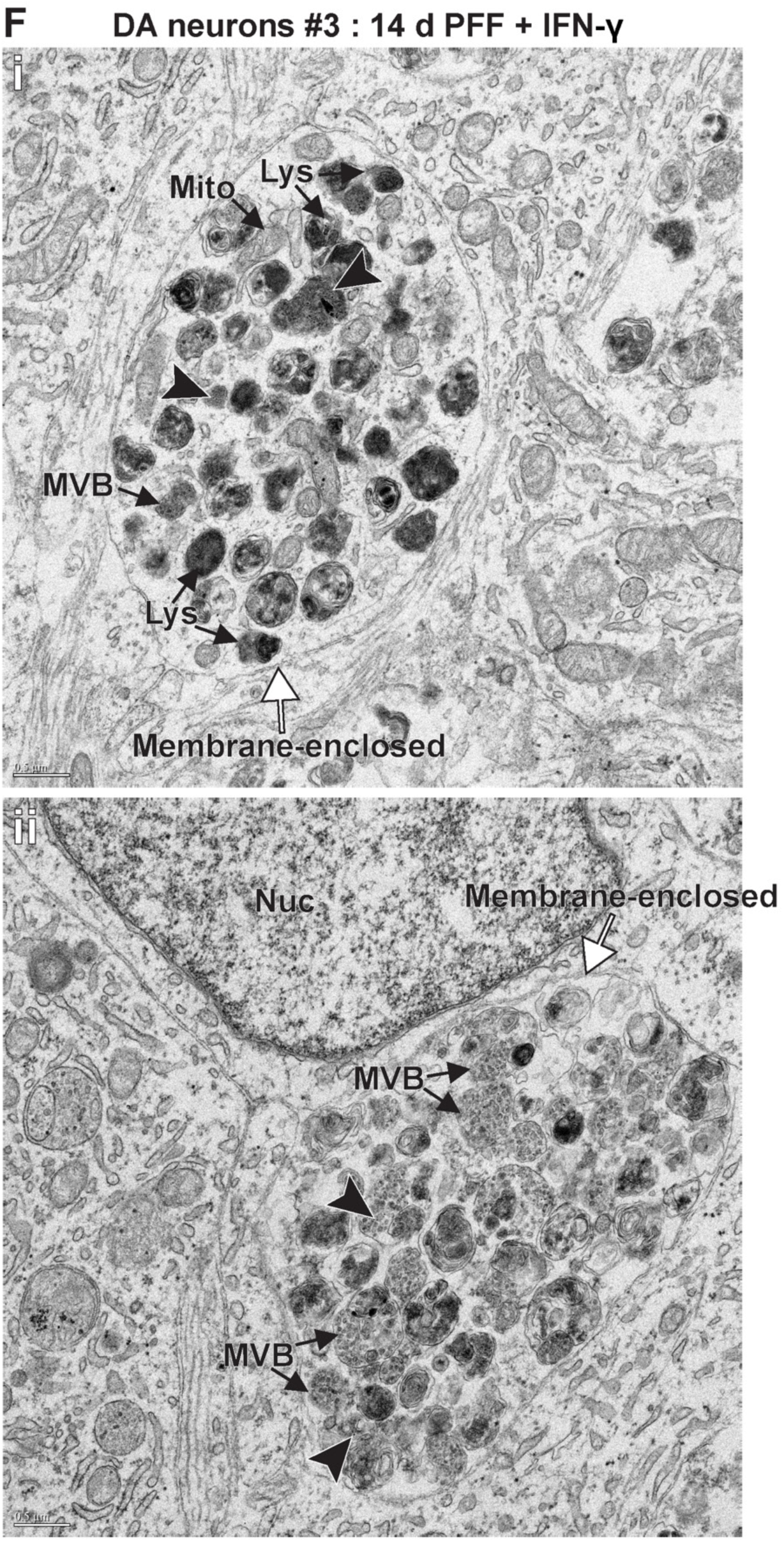
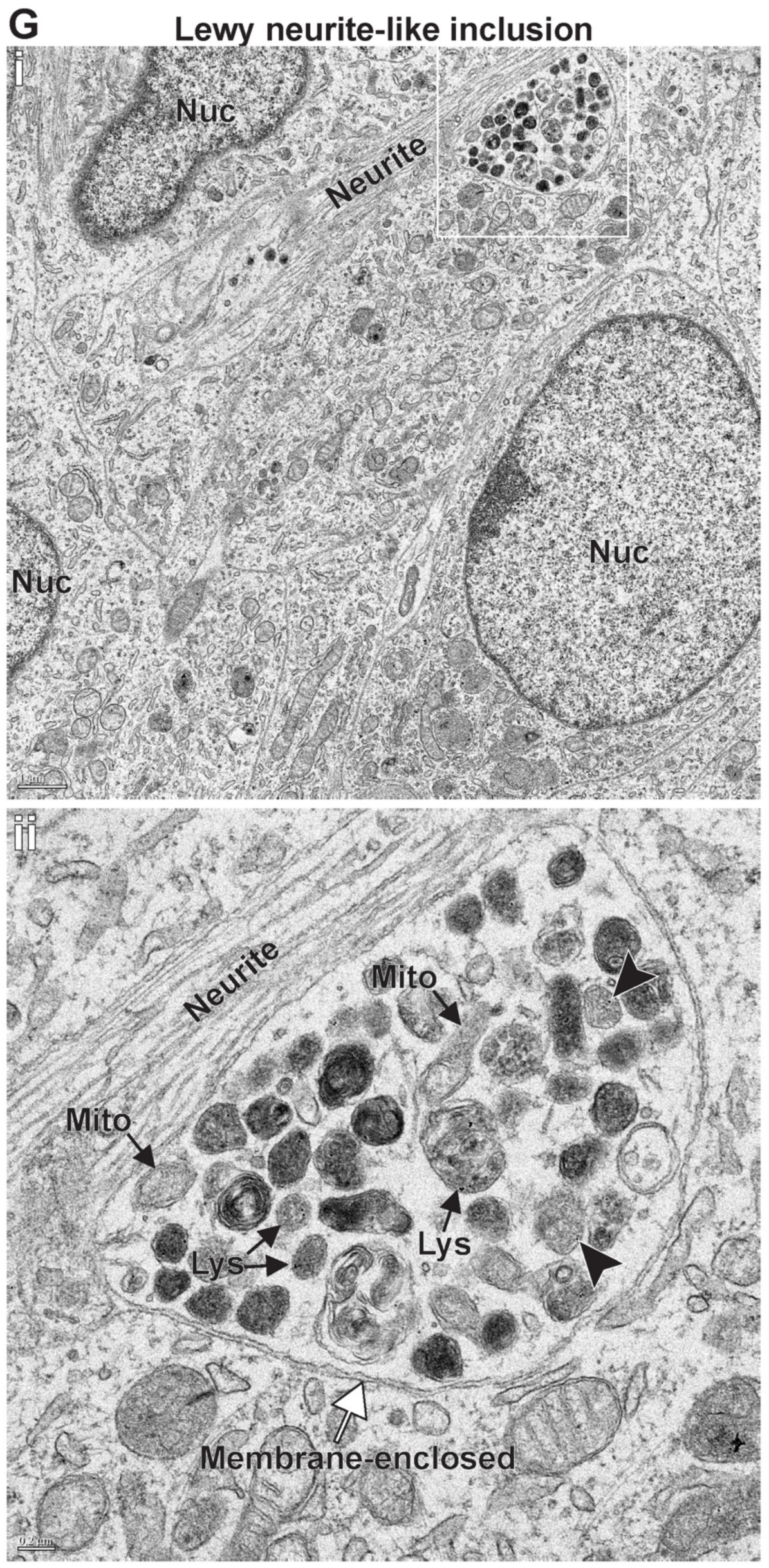
Collection of inclusions formed in DA neurons. (**A**-**C**) Inclusions formed within DA neurons generated from the AIW002-02 iPSC cell line. Inclusion showed in **A** contains a variety of organelles and filamentous (cytoskeletal) materials. Inclusion showed in **B**, consists of a collection of organelles but does not show a lot of filaments. In **C**, we see a mostly filamentous (a variety of filaments with different thickness) inclusion that contains islands of organelles at its edge. (**D** – **E**) shows inclusions, representative of the type of inclusions commonly found in the DYR-0100- derived DA neurons. (**F**) Inclusions formed within DA neurons generated from the 3450 iPSC cell lines. Similar to **E**, the inclusion shown in **i** is filled with lytic compartments (lysosomes and autolysosomes); however, **ii** shows a membrane-enclosed inclusion mostly filled with electron-dense MVBs. (**G**) Lewy neurite-like inclusion formed in DA neurons generated from AIW002-02. Lysosomes/autolysosomes (Lys), mitochondria (Mito), multivesicular bodies (MVB), filaments (Fil), white arrows point to the membrane surrounding the inclusions, white arrowheads point to nanogold-labeled PFFs found outside of lytic compartments, and dark arrowheads point to a few examples of nanogold-labeled PFF inside lytic vesicles.

**Supplementary Figure 4.**
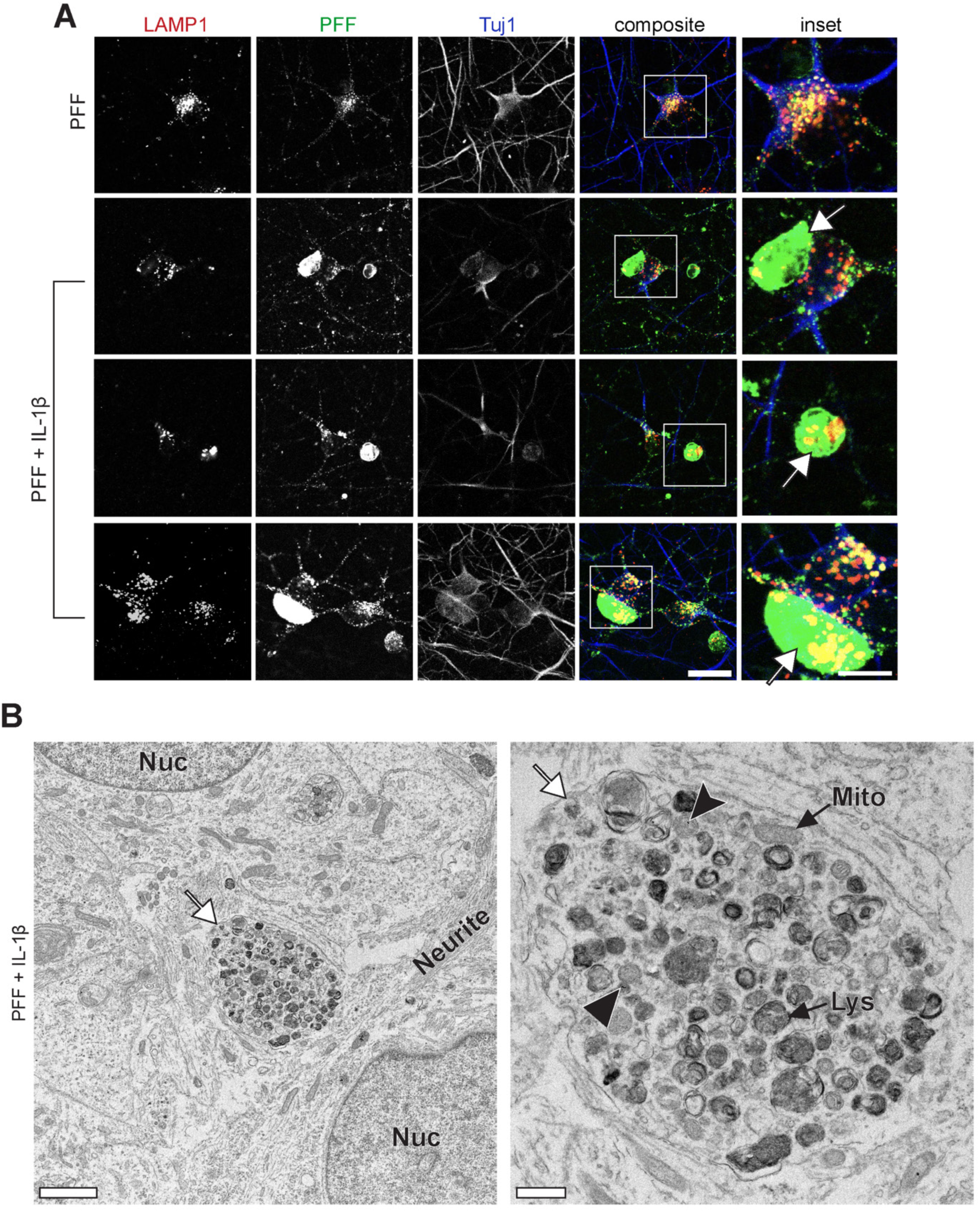
IL-1β administration following PFF treatment also leads to the formation of PFF-positive inclusions. (**A**) DA neurons were treated with PFF for 48 h, followed by 72 h rest following administration of 50 ng/mL of IL-1β for 24 h. The cells were then incubated in fresh media until day 14. Neurons were stained for LAMP1 and Tuj1. IL-1β data treated cells formed PFF-positive inclusions. Scale bar = 20 µm and 10 µm for insets. (**B**) Inclusions formed in neurons following dual hit treatment regime using IL-1β. Lysosomes/autolysosomes (Lys), mitochondria (Mito), white arrows point to the membrane surrounding the inclusions, and dark arrowheads point to a few examples of nanogold-labeled PFF inside lytic vesicles. Scale bar = 2 µm and 0.5 µm for insets.

**Supplementary Figure 5.**
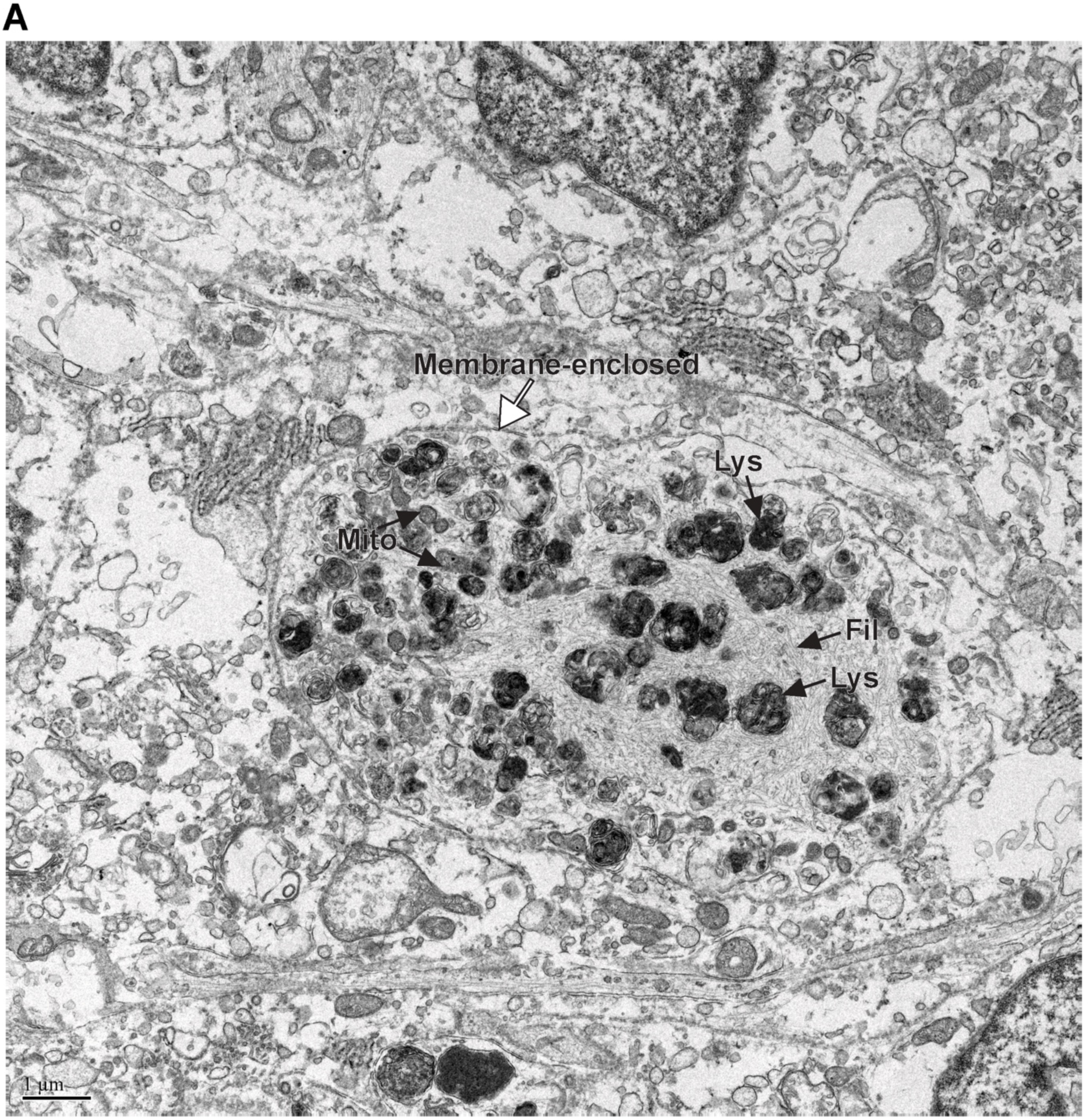
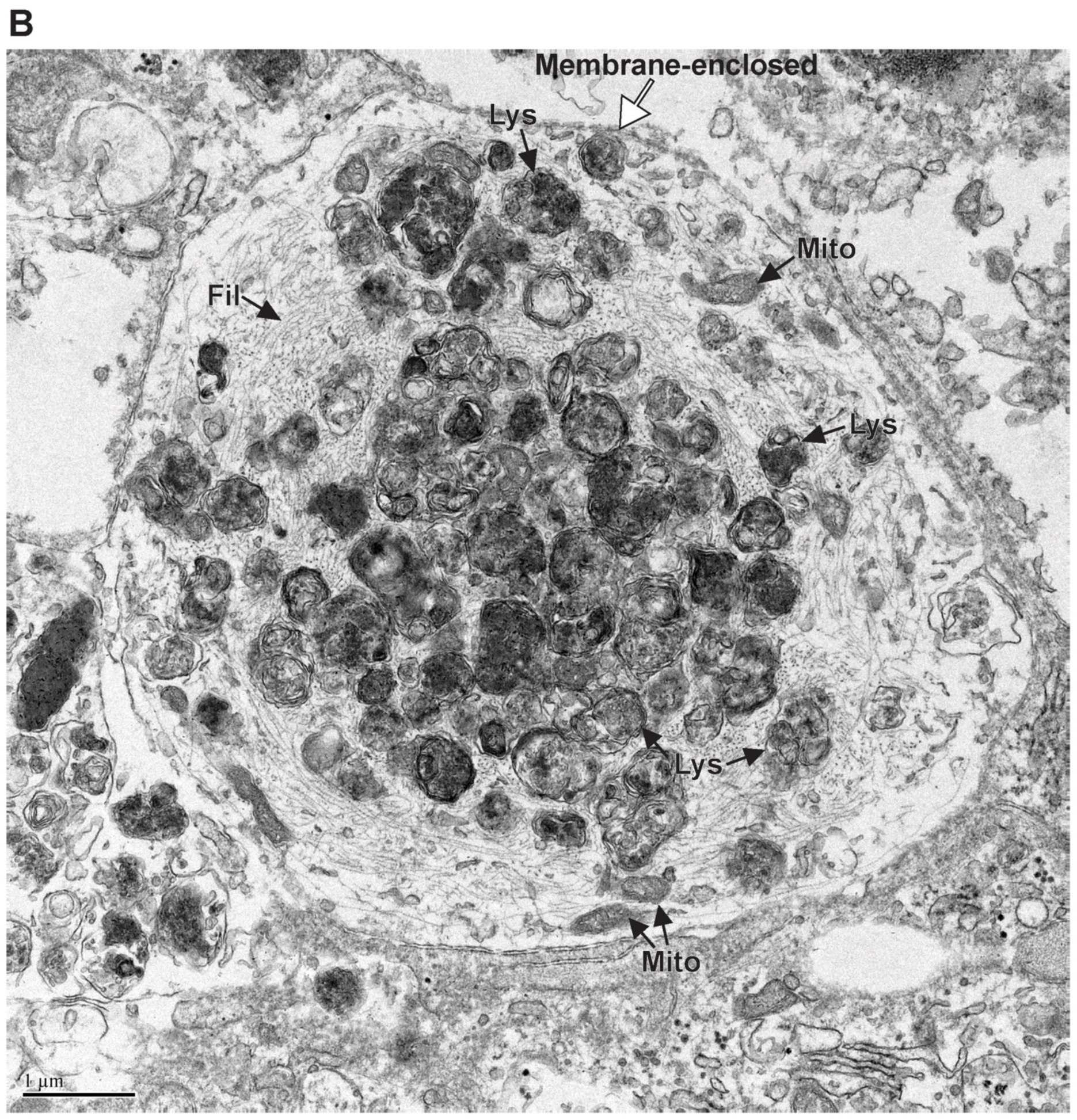
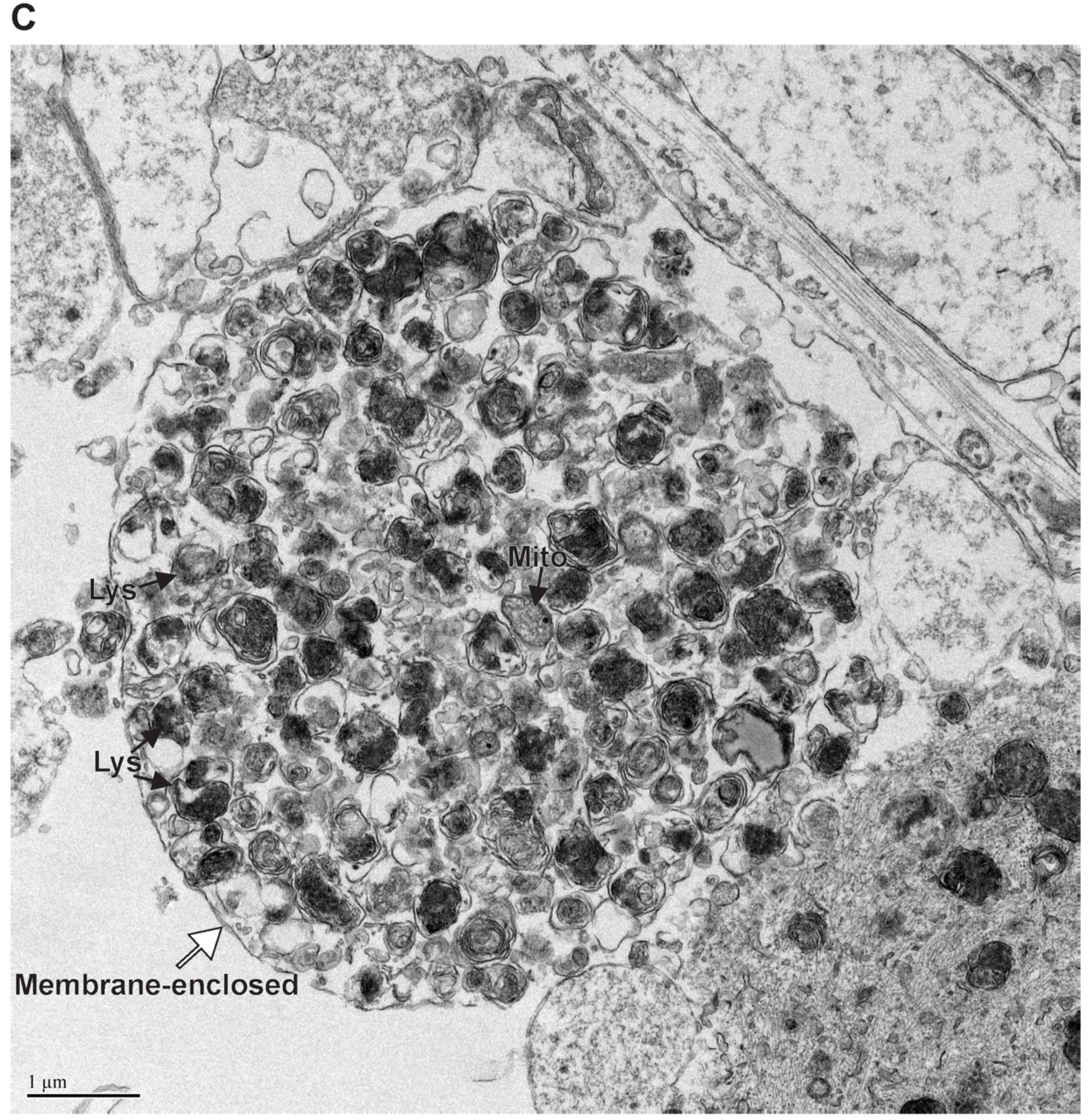
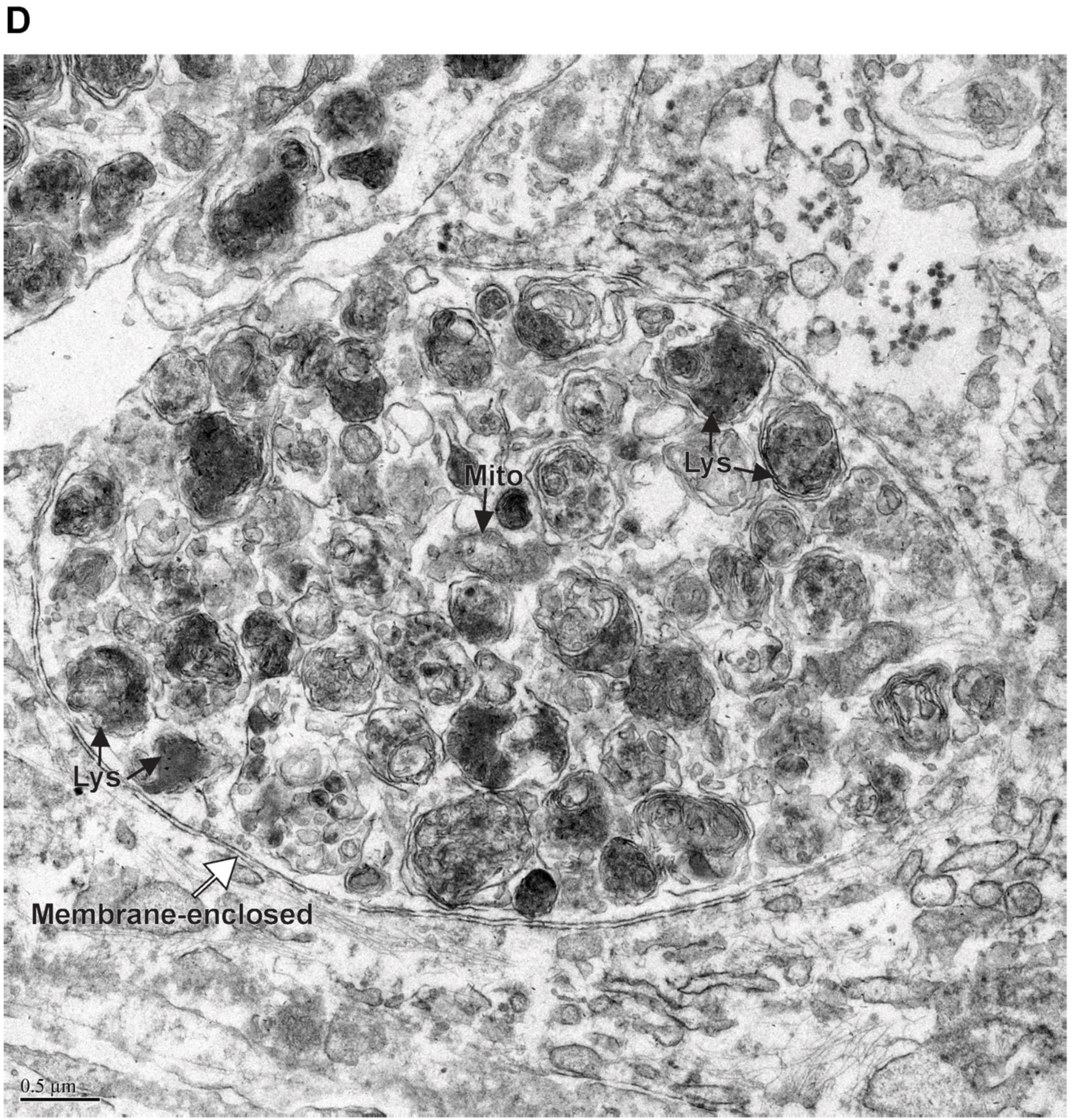
Collection of inclusions formed in DA neurons after 30 d of incubation. (**A**-**D**) Inclusions formed in DA neurons generated from the 3450 iPSC line. Neurons underwent the 14 d treatment regime, with PFF + IFN-γ treatment, and were then allowed to incubate an additional 16 d, for a total of 30 d in culture since the beginning of the treatment regime. The resulting inclusions are very tightly packed and compressed. Some inclusions contain a highly dense array of filaments (cytoskeletal), as seen in **A** and **B**. In contrast, others contain large amounts of seemingly dysfunctional organelles, as seen in **C** and **D**. Lysosomes/autolysosomes (Lys), mitochondria (Mito), multivesicular bodies (MVB), filaments (Fil), white arrows point to the membrane surrounding the inclusions.

**Supplementary Figure 6.**
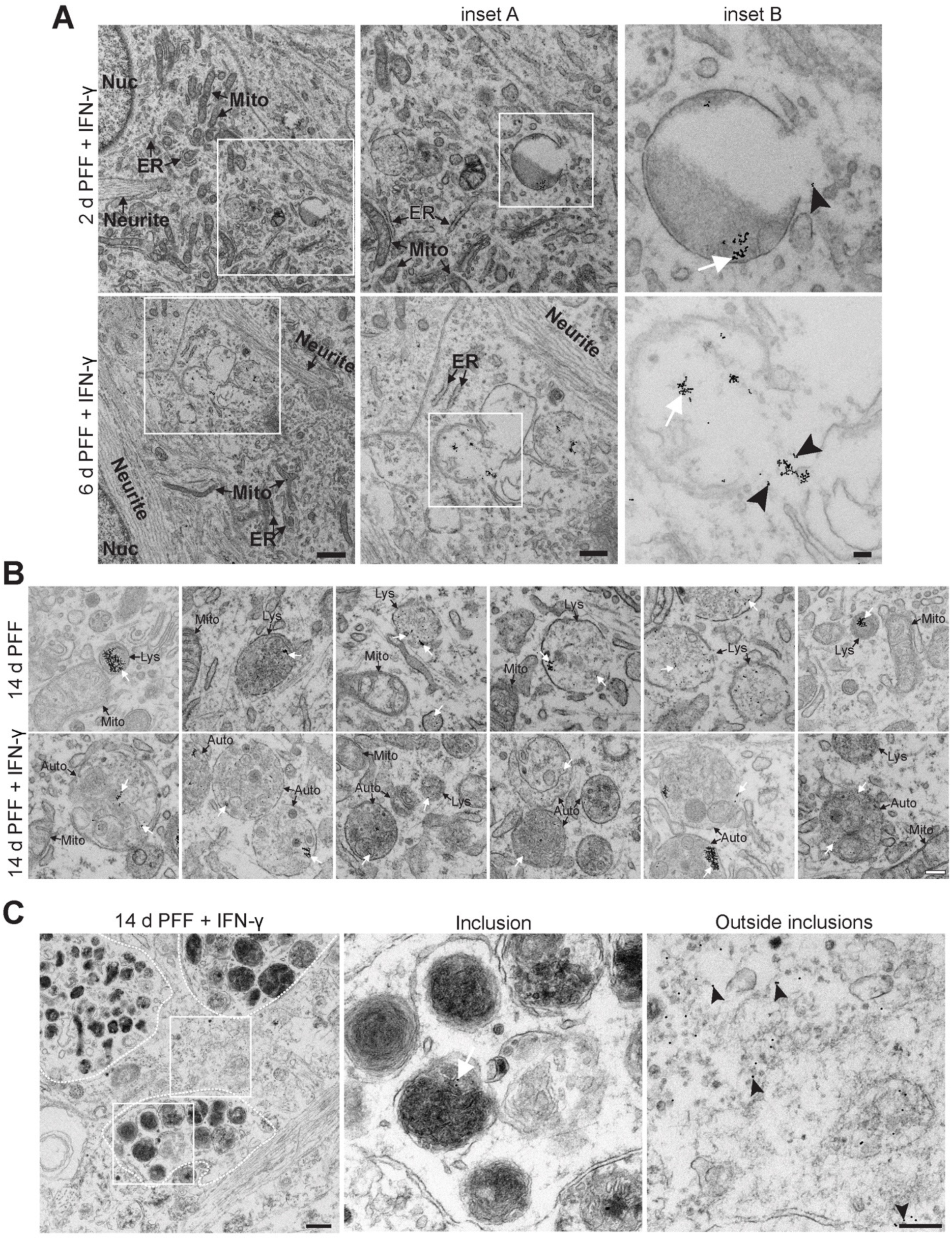
Lysosomal leakage of PFFs occurs in PFF + IFN-γ treated samples. **(A)** DA neurons that underwent the dual hit treatment regime and the more rapid 2 d treatment were fixed and processed for EM. PFFs can be seen accumulating in lysosomes, and some can be spotted outside of lysosomes as early as 2 d. More evidence for the leaking of PFFs into the cytosol can be seen with the 6 d samples. White arrows point to nanogold-labeled PFF inside lysosomal/autolysosome compartments, and arrowheads point to nanogold-labeled PFF outside lysosomes and in the cytosol. Few examples of endoplasmic reticulum (ER), mitochondria (Mito), and nuclei (Nuc) have been indicated. Scale bar = 1 µm, and 0.5 µm for inset A, and 100 nm for inset B. **(B)** Lytic compartments showing nanogold-PFF localization differ in the 14 d PFF-treated and the 14 d PFF + IFN-γ-treated samples. In the 14 d PFF-treated samples, very few nanogold-PFFs can be seen inside autolysosomes/autophagosomes, and lysosomes contain most of the internalized pool PFFs. In the 14 d PFF + IFN-γ-treated samples, PFFs are mostly localized in autolysosomes and autophagosomes. An accumulation of autolysosomes and autophagosomes can be seen in the PFF + IFN-γ-treated samples which are not present in the PFF-only samples. White arrows point to nanogold-labeled PFF inside lysosomal/autolysosome compartments. Scale bar = 200 nm. **(C)** In neurons where inclusions have formed, clear evidence of PFF leakage into the cytosol can be seen. Arrowheads point to cytosolic PFF. Scale bar = 500 nm and 200 nm for inset.

**Supplementary Figure 7.**
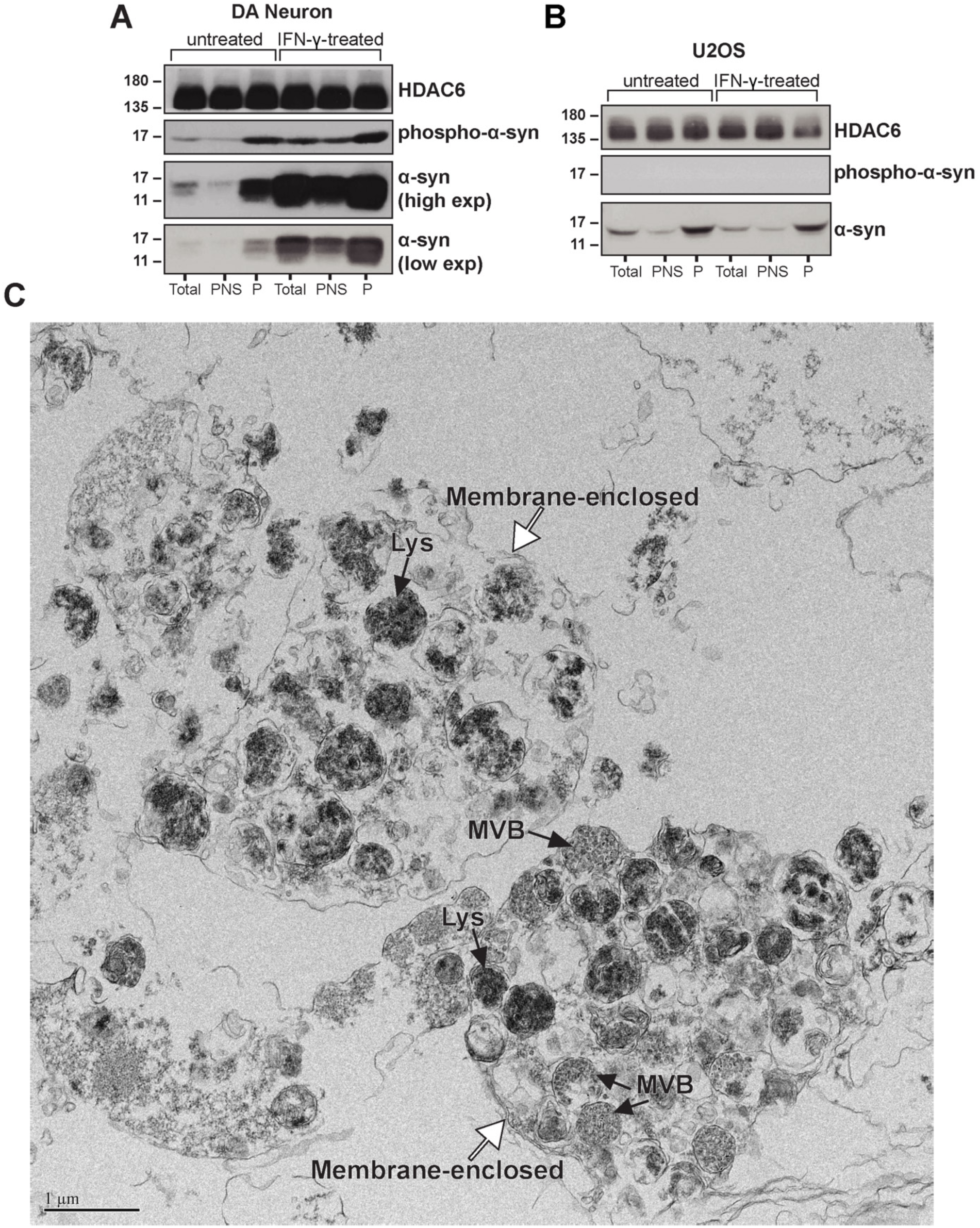
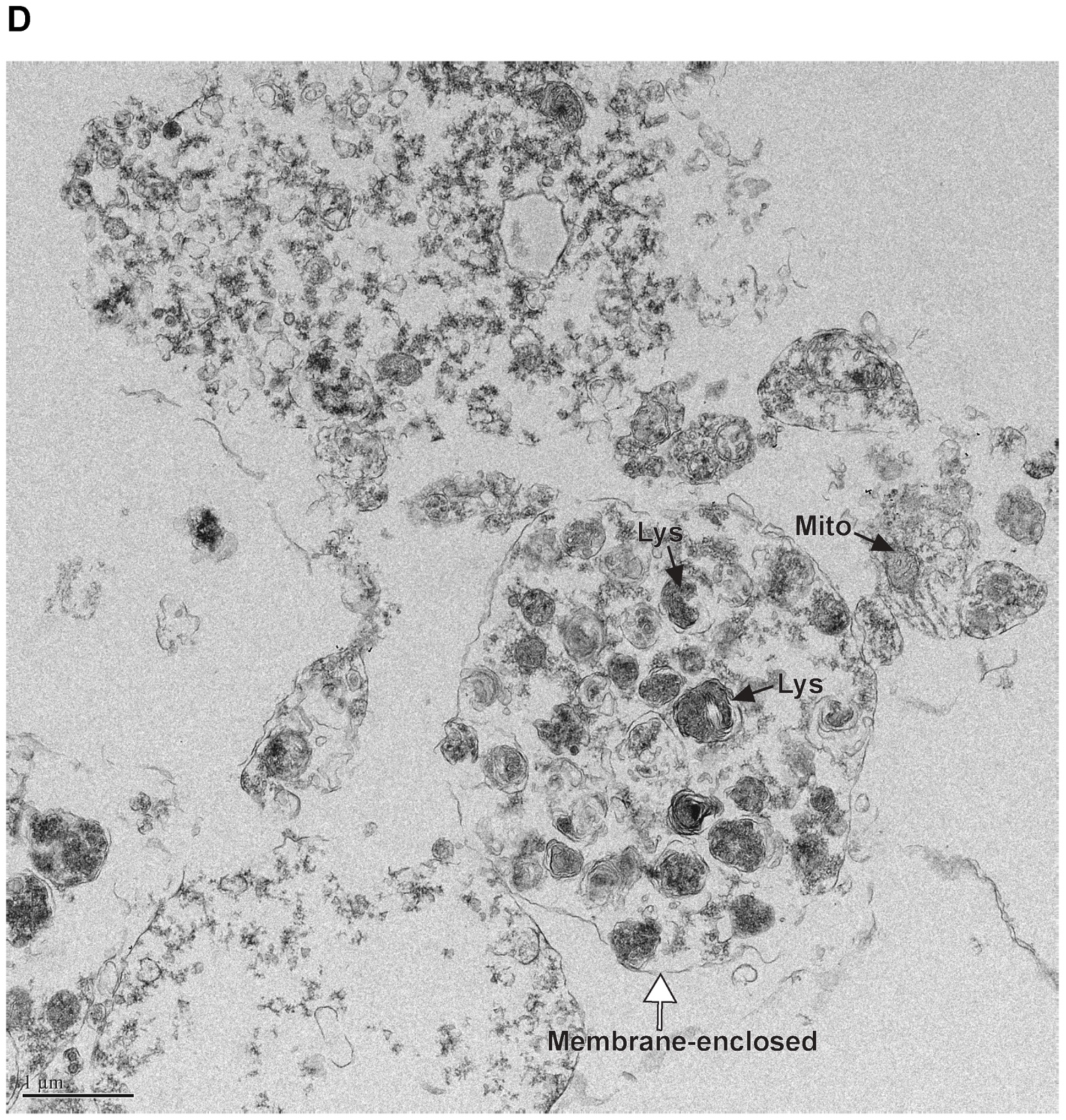
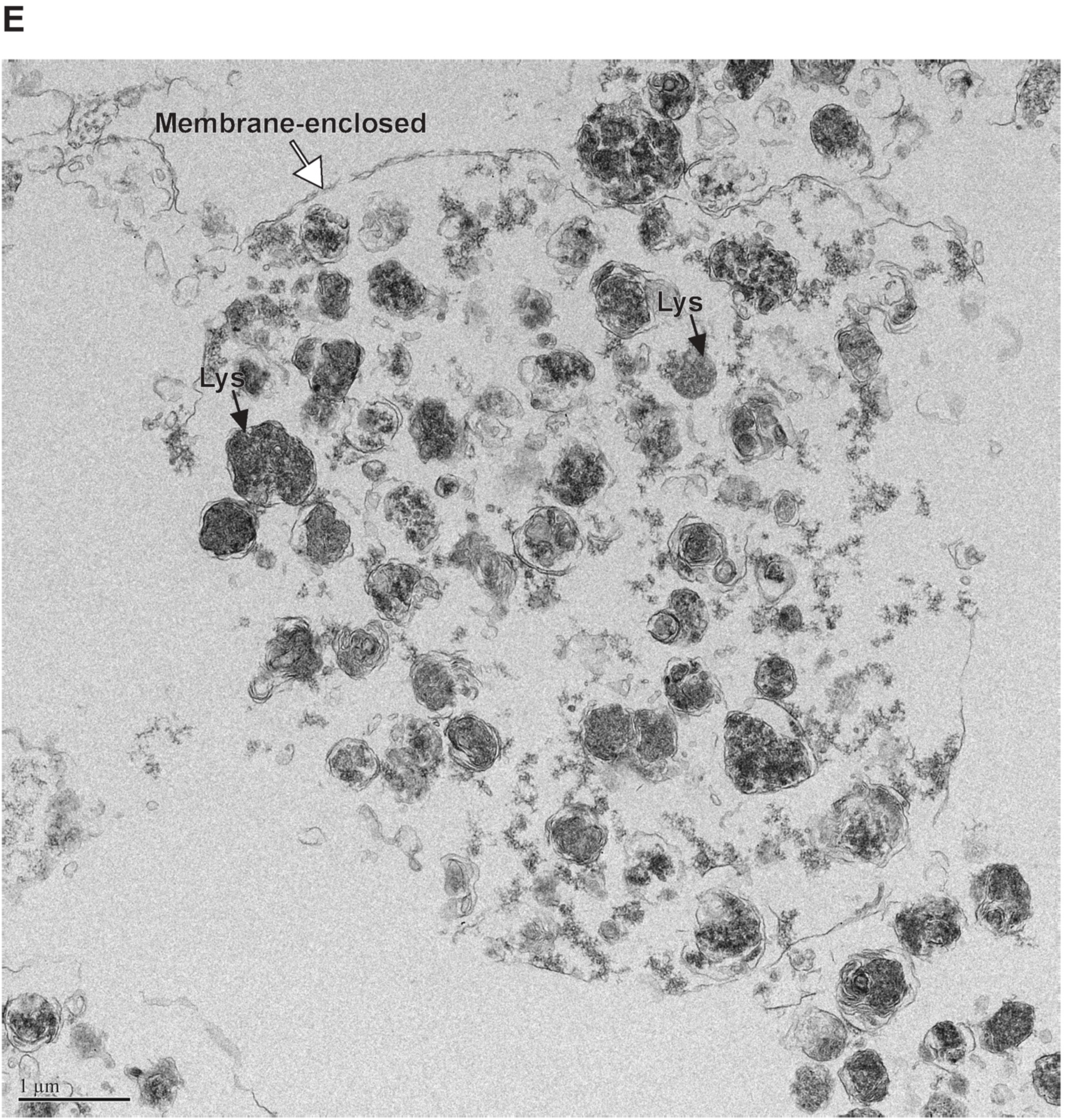
Isolated inclusions formed in DA neurons. (**A** and **B**) α-syn WB in DA neurons and U2OS in different subcellular fractions. In DA neurons, the LB-like inclusions and some enlarged lysosomes were pelleted at low speed after cellular homogenization in a detergent-free buffer. Control DA, where only PFF is added, shows the enrichment of α-syn in large organelles found in the pellet (P) compared to the total homogenate (total) and post-nuclear supernatant (PNS). IFN-γ-treated DA shows a much greater α-syn enrichment in the pellet, where more LB-like inclusions are collected compared to the control. In U2OS, where no LB-like inclusions are formed, no significant enrichment of α-syn was seen in the pellet, and no difference was observed between control and IFN-γ-treated cells. HDAC6 WB is shown as a loading control. Note that the difference in the total α-syn signal in the PFF condition compared to the PFF + IFN-γ-treated samples indicate a much more active degradation system in the PFF-only neurons compared to the PFF + IFN-γ-treated ones. Also note the double band in the α- syn signal, present (**C**-**E**) Inclusions isolated by subcellular fractionation were fixed, embedded, and sectioned for EM. Inclusions imaged were membrane-enclosed with the same morphology as those found in DA neurons. Lysosomes/autolysosomes (Lys), mitochondria (Mito), multivesicular bodies (MVB), and white arrows point to the membrane surrounding the inclusions.

**Supplementary Figure 8.**
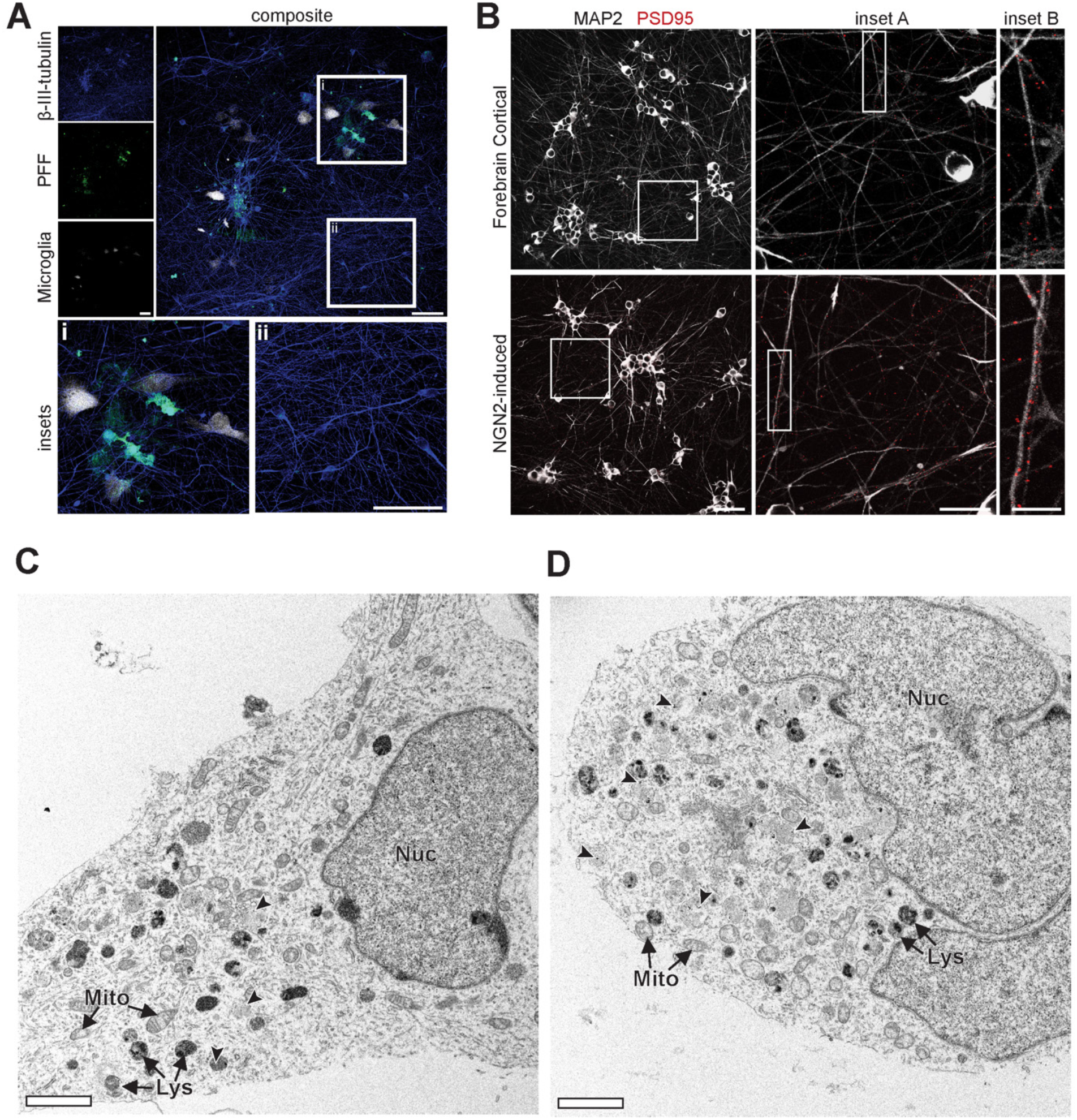
Microglia-neuron co-culture, characterization of forebrain and NGN2-induced neurons, and NGN2-induced neurons did not form inclusions. **(A)** Microglia, treated with LPS, were co-cultured with differentiated DA neurons with previous exposure to fluorescently labeled PFFs. Neurons near microglia were seen to form PFF-positive inclusions, as shown in i. Neurons not in proximity to microglia did not form inclusions and showed lower PFF fluorescence, as shown in ii. Scale bar = 80 µm and 50 µm for insets. **(B)** Cortical and NGN2-induced neurons were characterized using microtubule-associated protein 2 (MAP2) and post-synaptic density 95 (PSD95). Using a broad-spectrum induction process, forebrain cortical neurons derived from iPSC include excitatory and inhibitory neurons. NGN2-induced neurons are exclusively excitatory neurons and hence showed much higher levels of PSD95 fluorescence. Scale bar = 80 µm, 20 µm for inset A, and 5 µm for inset B. **(C and D)** EM images showing NGN2-induced excitatory neurons filled with swollen mitochondria and dark lysosomes; however, these compartments are not packaged into inclusions. Lysosomes/autolysosomes (Lys), mitochondria (Mito), and dark arrowheads point to examples of nanogold-labeled PFF inside lytic vesicles. Scale bar = 2 µm.

**Supplementary Figure 9.**
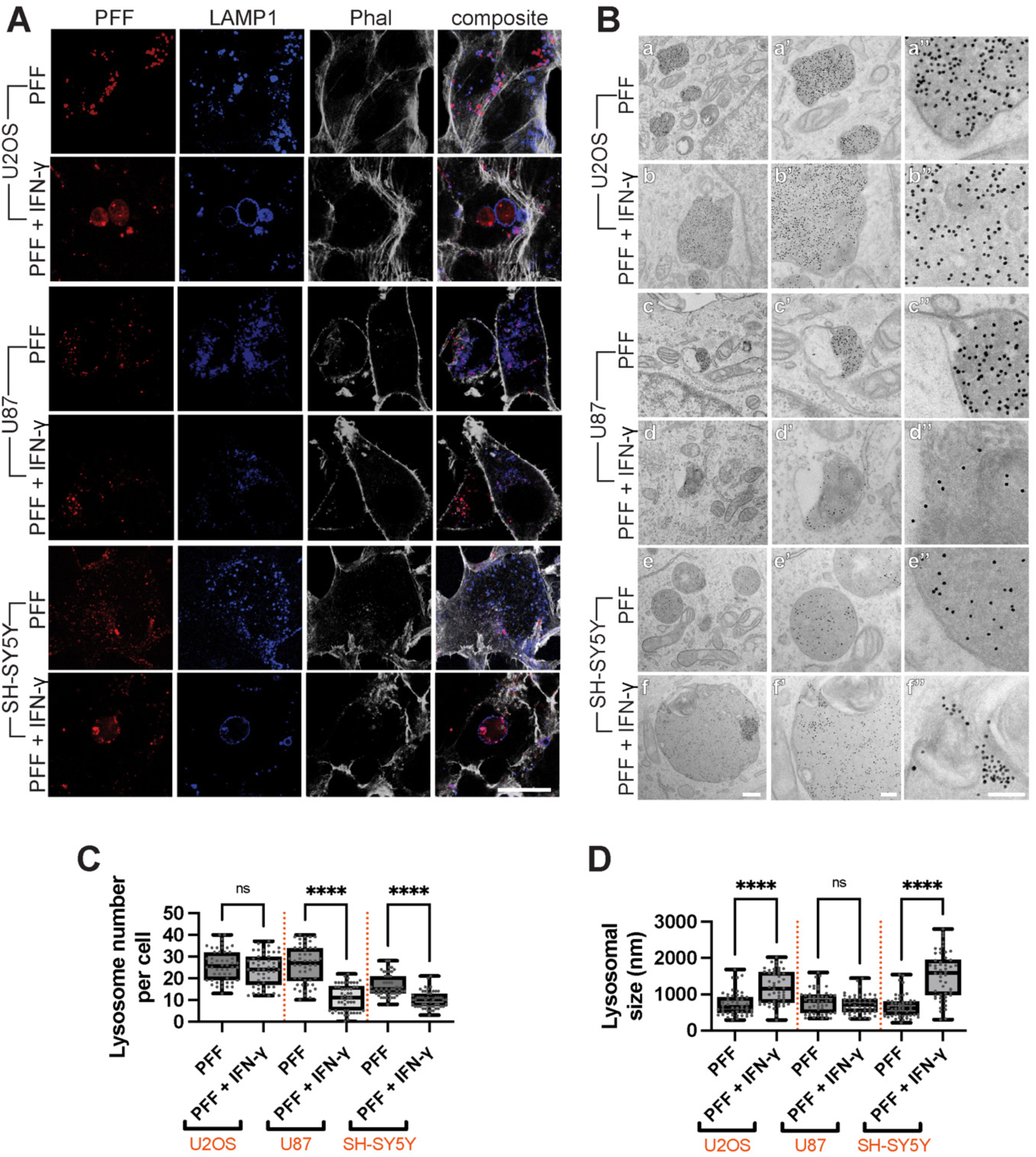
Cell lines do not form LB-like inclusions but show signs of lysosomal dysfunction. (**A**) U2OS, U87, and SH-SY5Y cells were given PFF and grown on 6-well plates before being passaged onto coverslips at low confluency to be treated with IFN-γ or PBS (control). Cells were then incubated for 8 d prior to fixation. Cells only given PFF showed LAMP1-PFF colocalization. U2OS and SH-SY5Y cells treated with PFF + IFN-γ formed large lysosomal structures, while U87 cells exhibited a loss in LAMP1 staining. Scale bar = 25 µm. (**B**) Cells underwent the 14 d dual hit treatment regime, described in **A**, except that they were given nanogold-labeled PFF and grown on 6-well plates prior to being passaged onto 8-well permanox chambers. They were then treated with IFN-γ or PBS and prepared for EM. U2OS and SH-SY5Y treated with PFF showed an accumulation of nanogold-PFF in electron-dense lysosomes. U2OS and SH-SY5Y cells treated with PFF + IFN-γ showed large lysosomal structures containing high amounts of nanogold-PFF. U87 cells exhibited abnormal lysosomal morphology in both IFN-γ-treated and untreated samples. Scale bar for a-f is 500 nm, for a’ to f’ is 200 nm, and for a’’ to f’’ is 100 nm. (**C**) The number of lysosomes (LAMP1-positive puncta) was quantified. U87 and SH-SY5Y showed a significant decrease in lysosomal number, n = 50 for each condition, and data was collected from five independent experiments. (**D**) The lysosomal size was calculated using electron micrographs collected following the dual hit treatment. U2OS and SH- SY5Y showed a significant increase in lysosomal size with IFN-γ treatment, n = 50 for each condition. Data were collected from two independent experiments. Both data collected in **C** and **D** were analyzed using one-way ANOVA with *post-hoc* Tukey’s test to compare means.

**Supplementary Figure 10.**
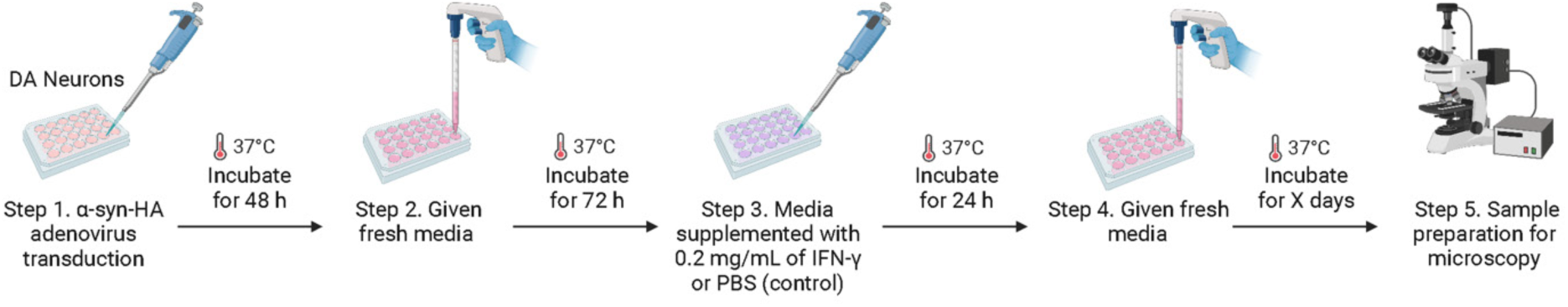
Protocol used to form phospho-α-syn positive inclusions through α-syn overexpression.

**Supplementary Figure 11.**
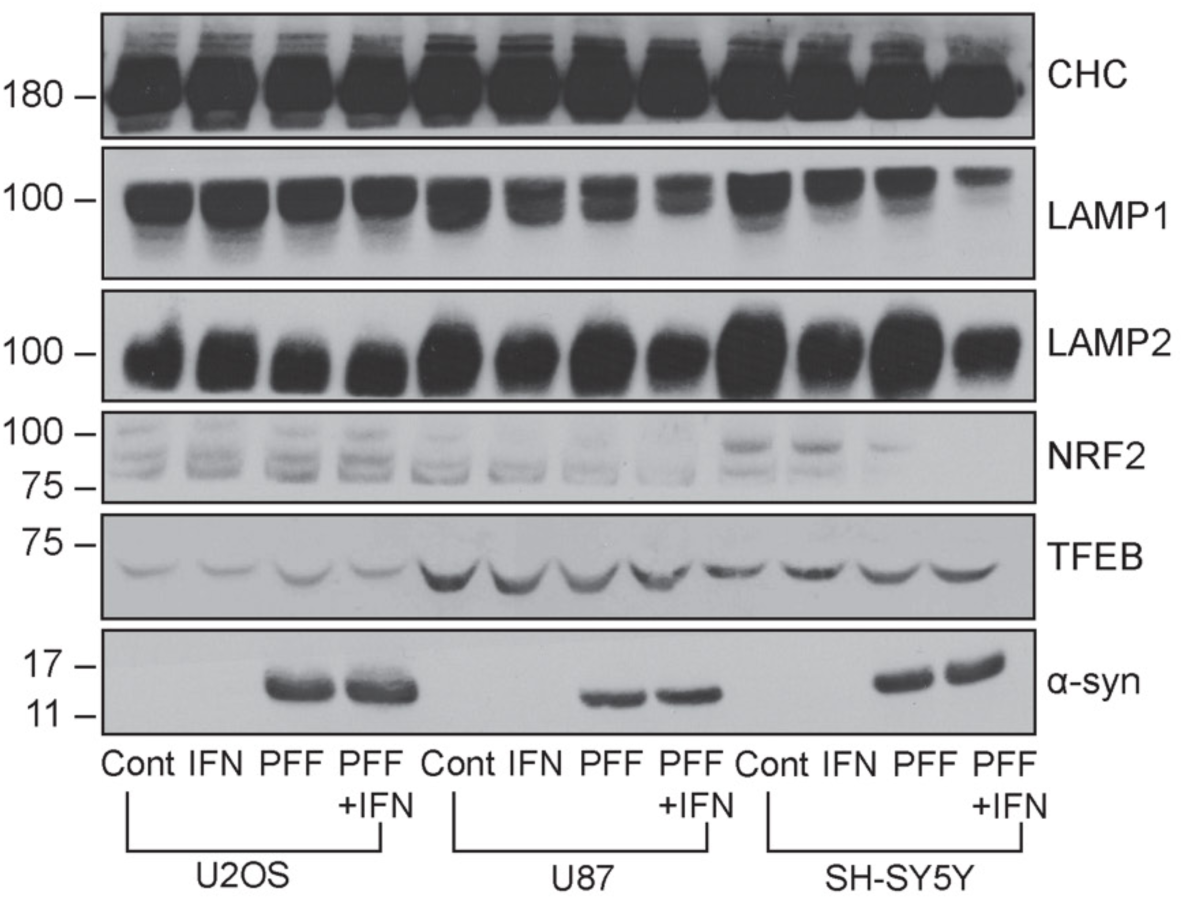
Lysosomal proteins are also affected in cell lines undergoing the dual hit treatment. U2OS, U87, and SH-SY5Y cells were given PFF and grown on 6-well plates at low confluency to be treated with IFN-γ or PBS (control). Cells were then incubated for 8 d prior to collection. Protein expression in cells that underwent the 14 d dual hit treatment regime (with or without IFN-γ treatment), along with control (labeled as Cont, No IFN-γ + No PFF) and IFN-γ were examined via WB. LAMP1, LAMP2, and NRF2 expression decreased with exposure to PFF and IFN-γ in U87 and SH-SY5Y, while TFEB was unaffected for all cell lines.

**Supplementary Figure 12.**
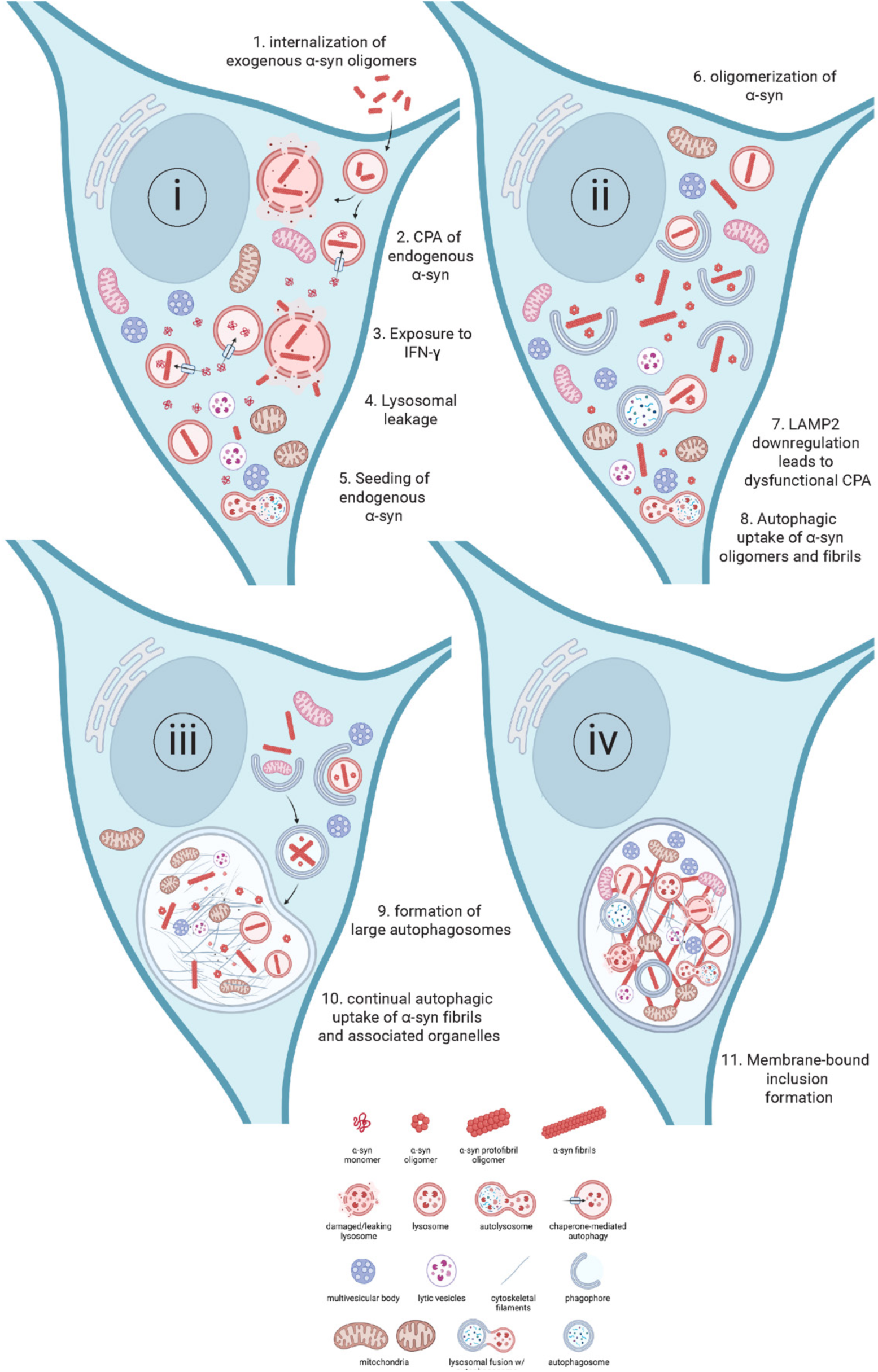
Possible mechanism for the formation of membrane-bound, membranous, organelle filled, LB-like inclusions. The internalization of oligomeric forms of α-syn, released by neighboring neurons serves as the first insult to lysosomal activity (1). The transport of monomeric α-syn into lysosomes through chaperone-mediated autophagy (CPA) leads to aggregation of α-syn inside lysosomes (2). Further lysosomal stress caused by IFN-γ results in lysosomal leakage (3 and 4). Leaking of oligomeric/misfolded α-syn leads to seeding and oligomerization of endogenous α-syn into aggregates (5 and 6). Downregulation of LAMP2 leads to dysfunctional CPA, leading to a buildup of endogenous α-syn, allowing more α-syn to be available for aggregation (7). To clear aggregates from the cytosol, autophagosomes form and take up α-syn aggregates and damaged organelles (8). Dysfunctional lysosomes are unable to fuse with autophagosomes, and even when fused with autophagosomes, are unable to degrade organelles and aggregates. This leads to the enlargement of autophagosomes (9). α-syn aggregation continues within the lumen of the autophagosomes (10), leading to the formation of membrane-bound LB-like inclusions (11).

**Supplementary Table 1.** Characteristics of LB-like inclusions.

| Characteristic | Method | Figure |
| --- | --- | --- |
| Filamentous | EM | Fig. 1C; Fig. 2B; Fig. 3A-C; Fig. S2C; Fig. S3A and C; Fig. S5A, B, and D; |
| Organellar Medley: Mitochondria | EM | Fig. 1D; Fig. 2B; Fig. 3B and C; Fig. S2C and D; Fig. S3A, B, C, E, F, G; Fig. S4B, Fig. S5A, B, D; |
| Organellar Medley: Lysosomes | EM, Isolated inclusions | Fig. 1C and D, Fig. 2B, Fig. 3A-C, Fig. S2C-D; Fig. S3A-G; Fig. S4C; Fig. S5A-D; Fig. S6C; Fig. S7C-E; |
| Organellar Medley: MVBs | EM, Isolated inclusions | Fig. 2B; Fig. 3C; Fig. S2D; Fig. S3A, E, and F; Fig. S7C; |
| Organellar Medley: ER | EM | Fig. 3C; Fig S2C |
| PFF-positive | EM, Immunofluorescence, Western blot of isolated inclusions | Fig. 1B-D; Fig. 2; Fig. 4A; Fig. 5B and F; Fig. 7B; Fig. 8B; Fig. S2C and D; Fig. S3A-G; Fig. S4A and B; Fig. S6C; Fig. S7A |
| Phospho-α-syn-positive | Immunofluorescence, Western blot of isolated inclusions | Fig. 5B, C, I, K, and L; Fig. S7A; |
| LC3B-positive | Immunofluorescence | Fig. 7B |
| Membrane-enclosed | EM | Fig. 1D; Fig. 3; Fig. S2C and D; Fig. S3A-G; Fig. S4B; Fig. S5A-D; Fig. S7C-E |

## Notes

http://dx.doi.org/10.17632/kp3774p92d.1

### Competing Interest Statement

The authors have declared no competing interest.

### Summary of Updates

Based on new submission

## References

1. Spillantini, M.G., Schmidt, M.L., Lee, V.M., Trojanowski, J.Q., Jakes, R., and Goedert, M. (1997). Alpha-synuclein in Lewy bodies. Nature 388, 839–840. 10.1038/42166.

2. Baba, M., Nakajo, S., Tu, P.H., Tomita, T., Nakaya, K., Lee, V.M., Trojanowski, J.Q., and Iwatsubo, T. (1998). Aggregation of alpha-synuclein in Lewy bodies of sporadic Parkinson’s disease and dementia with Lewy bodies. The American journal of pathology 152, 879–884.

3. Braak, H., Tredici, K.D., Rüb, U., de Vos, R.A.I., Jansen Steur, E.N.H., and Braak, E. (2003). Staging of brain pathology related to sporadic Parkinson’s disease. Neurobiology of Aging 24, 197–211. 10.1016/s0197-4580(02)00065-9.

4. Rajput, A.H., Offord, K.P., Beard, C.M., and Kurland, L.T. (1984). Epidemiology of parkinsonism: incidence, classification, and mortality. Annals of neurology 16, 278–282. 10.1002/ana.410160303.

5. Braak, H., Sandmann-Keil, D., Gai, W., and Braak, E. (1999). Extensive axonal Lewy neurites in Parkinson’s disease: a novel pathological feature revealed by alpha-synuclein immunocytochemistry. Neurosci Lett 265, 67–69. 10.1016/s0304-3940(99)00208-6.

6. Wakabayashi, K., Tanji, K., Odagiri, S., Miki, Y., Mori, F., and Takahashi, H. (2013). The Lewy body in Parkinson’s disease and related neurodegenerative disorders. Mol Neurobiol 47, 495–508. 10.1007/s12035-012-8280-y.

7. Kim, W.S., Kagedal, K., and Halliday, G.M. (2014). Alpha-synuclein biology in Lewy body diseases. Alzheimers Res Ther 6, 73. 10.1186/s13195-014-0073-2.

8. Engelhardt, E., and Gomes, M.D.M. (2017). Lewy and his inclusion bodies: Discovery and rejection. Dement Neuropsychol 11, 198–201. 10.1590/1980-57642016dn11-020012.

9. Glausier, J.R., Konanur, A., and Lewis, D.A. (2019). Factors Affecting Ultrastructural Quality in the Prefrontal Cortex of the Postmortem Human Brain. J Histochem Cytochem 67, 185-

10. 202. 10.1369/0022155418819481.

10. Shahmoradian, S.H., Lewis, A.J., Genoud, C., Hench, J., Moors, T.E., Navarro, P.P., Castano-Diez, D., Schweighauser, G., Graff-Meyer, A., Goldie, K.N., et al. (2019). Lewy pathology in Parkinson’s disease consists of crowded organelles and lipid membranes. Nat Neurosci 22, 1099–1109. 10.1038/s41593-019-0423-2.

11. Soper, J.H., Roy, S., Stieber, A., Lee, E., Wilson, R.B., Trojanowski, J.Q., Burd, C.G., and Lee, V.M.-Y. (2008). α-Synuclein–induced Aggregation of Cytoplasmic Vesicles in Saccharomyces cerevisiae. Molecular biology of the cell 19, 1093–1103.

12. Gai, W., Yuan, H., Li, X., Power, J., Blumbergs, P., and Jensen, P. (2000). In situ and in vitro study of colocalization and segregation of α-synuclein, ubiquitin, and lipids in Lewy bodies. Experimental neurology 166, 324–333.

13. Forno, L.S., and Norville, R.L. (1976). Ultrastructure of Lewy bodies in the stellate ganglion. Acta Neuropathol 34, 183–197. 10.1007/BF00688674.

14. Frigerio, R., Fujishiro, H., Ahn, T.B., Josephs, K.A., Maraganore, D.M., DelleDonne, A., Parisi, J.E., Klos, K.J., Boeve, B.F., Dickson, D.W., and Ahlskog, J.E. (2011). Incidental Lewy body disease: do some cases represent a preclinical stage of dementia with Lewy bodies? Neurobiol Aging 32, 857–863. 10.1016/j.neurobiolaging.2009.05.019.

15. Monaco, A., and Fraldi, A. (2020). Protein Aggregation and Dysfunction of Autophagy-Lysosomal Pathway: A Vicious Cycle in Lysosomal Storage Diseases. Front Mol Neurosci 13, 37. 10.3389/fnmol.2020.00037.

16. Klein, A.D., and Mazzulli, J.R. (2018). Is Parkinson’s disease a lysosomal disorder? Brain 141, 2255–2262. 10.1093/brain/awy147.

17. Durcan, T.M., and Fon, E.A. (2015). The three ‘P’s of mitophagy: PARKIN, PINK1, and post-translational modifications. Genes Dev *29*, 989-999. 10.1101/gad.262758.115.

18. Sato, S., Uchihara, T., Fukuda, T., Noda, S., Kondo, H., Saiki, S., Komatsu, M., Uchiyama, Y., Tanaka, K., and Hattori, N. (2018). Loss of autophagy in dopaminergic neurons causes Lewy pathology and motor dysfunction in aged mice. Sci Rep 8, 2813. 10.1038/s41598- 018-21325-w.

19. Boman, A., Svensson, S., Boxer, A., Rojas, J.C., Seeley, W.W., Karydas, A., Miller, B., Kagedal, K., and Svenningsson, P. (2016). Distinct Lysosomal Network Protein Profiles in Parkinsonian Syndrome Cerebrospinal Fluid. J Parkinsons Dis 6, 307–315. 10.3233/JPD-150759.

20. Navarro-Romero, A., Montpeyo, M., and Martinez-Vicente, M. (2020). The Emerging Role of the Lysosome in Parkinson’s Disease. Cells 9. 10.3390/cells9112399.

21. Eskelinen, E.L., Illert, A.L., Tanaka, Y., Schwarzmann, G., Blanz, J., Von Figura, K., and Saftig, P. (2002). Role of LAMP-2 in lysosome biogenesis and autophagy. Mol Biol Cell 13, 3355–3368. 10.1091/mbc.e02-02-0114.

22. Eskelinen, E.L. (2006). Roles of LAMP-1 and LAMP-2 in lysosome biogenesis and autophagy. Mol Aspects Med 27, 495–502. 10.1016/j.mam.2006.08.005.

23. Matheoud, D., Cannon, T., Voisin, A., Penttinen, A.M., Ramet, L., Fahmy, A.M., Ducrot, C., Laplante, A., Bourque, M.J., Zhu, L., et al. (2019). Intestinal infection triggers Parkinson’s disease-like symptoms in Pink1(-/-) mice. Nature 571, 565–569. 10.1038/s41586-019-1405-y.

24. Panagiotakopoulou, V., Ivanyuk, D., De Cicco, S., Haq, W., Arsic, A., Yu, C., Messelodi, D., Oldrati, M., Schondorf, D.C., Perez, M.J., et al. (2020). Interferon-gamma signaling synergizes with LRRK2 in neurons and microglia derived from human induced pluripotent stem cells. Nat Commun 11, 5163. 10.1038/s41467-020-18755-4.

25. Ahmadi Rastegar, D., Hughes, L.P., Perera, G., Keshiya, S., Zhong, S., Gao, J., Halliday, G.M., Schule, B., and Dzamko, N. (2022). Effect of LRRK2 protein and activity on stimulated cytokines in human monocytes and macrophages. NPJ Parkinsons Dis 8, 34. 10.1038/s41531-022-00297-9.

26. Burberry, A., Wells, M.F., Limone, F., Couto, A., Smith, K.S., Keaney, J., Gillet, G., van Gastel, N., Wang, J.Y., Pietilainen, O., et al. (2020). C9orf72 suppresses systemic and neural inflammation induced by gut bacteria. Nature 582, 89–94. 10.1038/s41586-020-2288-7.

27. Kulkarni, A., Ganesan, P., and O’Donnell, L.A. (2016). Interferon Gamma: Influence on Neural Stem Cell Function in Neurodegenerative and Neuroinflammatory Disease. Clin Med Insights Pathol 9, 9–19. 10.4137/CPath.S40497.

28. Roy, E., and Cao, W. (2022). Glial interference: impact of type I interferon in neurodegenerative diseases. Mol Neurodegener 17, 78. 10.1186/s13024-022-00583-3.

29. Seifert, H.A., Collier, L.A., Chapman, C.B., Benkovic, S.A., Willing, A.E., and Pennypacker, K.R. (2014). Pro-inflammatory interferon gamma signaling is directly associated with stroke induced neurodegeneration. J Neuroimmune Pharmacol 9, 679–689. 10.1007/s11481-014-9560-2.

30. Bayati, A., Banks, E., Han, C., Luo, W., Reintsch, W.E., Zorca, C.E., Shlaifer, I., Del Cid Pellitero, E., Vanderperre, B., McBride, H.M., et al. (2022). Rapid macropinocytic transfer of alpha-synuclein to lysosomes. Cell Rep 40, 111102. 10.1016/j.celrep.2022.111102.

31. Kawanokuchi, J., Mizuno, T., Takeuchi, H., Kato, H., Wang, J., Mitsuma, N., and Suzumura, A. (2006). Production of interferon-gamma by microglia. Mult Scler 12, 558–564. 10.1177/1352458506070763.

32. Wang, X., and Suzuki, Y. (2007). Microglia produce IFN-gamma independently from T cells during acute toxoplasmosis in the brain. J Interferon Cytokine Res 27, 599–605. 10.1089/jir.2006.0157.

33. Lv, D., Liu, Y., Guo, F., Wu, A., Mo, Y., Wang, S., and Chu, J. (2020). Combining interferon-gamma release assays with lymphocyte enumeration for diagnosis of Mycobacterium tuberculosis infection. J Int Med Res 48, 300060520925660. 10.1177/0300060520925660.

34. Yang, Y., Wang, H.J., Hu, W.L., Bai, G.N., and Hua, C.Z. (2022). Diagnostic Value of Interferon-Gamma Release Assays for Tuberculosis in the Immunocompromised Population. Diagnostics (Basel) 12. 10.3390/diagnostics12020453.

35. Ghanekar, S.A., Nomura, L.E., Suni, M.A., Picker, L.J., Maecker, H.T., and Maino, V.C. (2001). Gamma interferon expression in CD8(+) T cells is a marker for circulating cytotoxic T lymphocytes that recognize an HLA A2-restricted epitope of human cytomegalovirus phosphoprotein pp65. Clin Diagn Lab Immunol 8, 628–631. 10.1128/CDLI.8.3.628-631.2001.

36. Jorgovanovic, D., Song, M., Wang, L., and Zhang, Y. (2020). Roles of IFN-gamma in tumor progression and regression: a review. Biomark Res 8, 49. 10.1186/s40364-020-00228-x.

37. Nicolet, B.P., Guislain, A., van Alphen, F.P.J., Gomez-Eerland, R., Schumacher, T.N.M., van den Biggelaar, M., and Wolkers, M.C. (2020). CD29 identifies IFN-gamma-producing human CD8(+) T cells with an increased cytotoxic potential. Proc Natl Acad Sci U S A 117, 6686–6696. 10.1073/pnas.1913940117.

38. Fang, C., Weng, T., Hu, S., Yuan, Z., Xiong, H., Huang, B., Cai, Y., Li, L., and Fu, X. (2021). IFN-gamma-induced ER stress impairs autophagy and triggers apoptosis in lung cancer cells. Oncoimmunology 10, 1962591. 10.1080/2162402X.2021.1962591.

39. Soper, J.H., Roy, S., Stieber, A., Lee, E., Wilson, R.B., Trojanowski, J.Q., Burd, C.G., and Lee, V.M. (2008). Alpha-synuclein-induced aggregation of cytoplasmic vesicles in Saccharomyces cerevisiae. Mol Biol Cell 19, 1093–1103. 10.1091/mbc.e07-08-0827.

40. Bieri, G., Gitler, A.D., and Brahic, M. (2018). Internalization, axonal transport and release of fibrillar forms of alpha-synuclein. Neurobiol Dis 109, 219–225. 10.1016/j.nbd.2017.03.007.

41. Freeman, D., Cedillos, R., Choyke, S., Lukic, Z., McGuire, K., Marvin, S., Burrage, A.M., Sudholt, S., Rana, A., O’Connor, C., et al. (2013). Alpha-synuclein induces lysosomal rupture and cathepsin dependent reactive oxygen species following endocytosis. PLoS One 8, e62143. 10.1371/journal.pone.0062143.

42. Jiang, P., Gan, M., Yen, S.H., McLean, P.J., and Dickson, D.W. (2017). Impaired endo-lysosomal membrane integrity accelerates the seeding progression of alpha-synuclein aggregates. Sci Rep 7, 7690. 10.1038/s41598-017-08149-w.

43. Vergarajauregui, S., Connelly, P.S., Daniels, M.P., and Puertollano, R. (2008). Autophagic dysfunction in mucolipidosis type IV patients. Hum Mol Genet 17, 2723–2737. 10.1093/hmg/ddn174.

44. Rezai-Zadeh, K., Gate, D., and Town, T. (2009). CNS infiltration of peripheral immune cells: D-Day for neurodegenerative disease? J Neuroimmune Pharmacol 4, 462–475. 10.1007/s11481-009-9166-2.

45. Yang, Q., Wang, G., and Zhang, F. (2020). Role of Peripheral Immune Cells-Mediated Inflammation on the Process of Neurodegenerative Diseases. Front Immunol 11, 582825. 10.3389/fimmu.2020.582825.

46. Lees, J.R., and Cross, A.H. (2007). A little stress is good: IFN-gamma, demyelination, and multiple sclerosis. J Clin Invest 117, 297–299. 10.1172/JCI31254.

47. Duffy, S.S., Lees, J.G., and Moalem-Taylor, G. (2014). The contribution of immune and glial cell types in experimental autoimmune encephalomyelitis and multiple sclerosis. Mult Scler Int 2014, 285245. 10.1155/2014/285245.

48. Molteni, M., and Rossetti, C. (2017). Neurodegenerative diseases: The immunological perspective. J Neuroimmunol 313, 109–115. 10.1016/j.jneuroim.2017.11.002.

49. De Simone, R., Levi, G., and Aloisi, F. (1998). Interferon gamma gene expression in rat central nervous system glial cells. Cytokine 10, 418–422. 10.1006/cyto.1997.0314.

50. Suzuki, Y., Claflin, J., Wang, X., Lengi, A., and Kikuchi, T. (2005). Microglia and macrophages as innate producers of interferon-gamma in the brain following infection with Toxoplasma gondii. Int J Parasitol 35, 83–90. 10.1016/j.ijpara.2004.10.020.

51. Liu, X., and Quan, N. (2018). Microglia and CNS Interleukin-1: Beyond Immunological Concepts. Front Neurol 9, 8. 10.3389/fneur.2018.00008.

52. Zhang, W., Wang, T., Pei, Z., Miller, D.S., Wu, X., Block, M.L., Wilson, B., Zhang, W., Zhou, Y., Hong, J.S., and Zhang, J. (2005). Aggregated alpha-synuclein activates microglia: a process leading to disease progression in Parkinson’s disease. FASEB J 19, 533–542. 10.1096/fj.04-2751com.

53. Zhang, L., Wang, M., Sterling, N.W., Lee, E.Y., Eslinger, P.J., Wagner, D., Du, G., Lewis, M.M., Truong, Y., Bowman, F.D., and Huang, X. (2018). Cortical Thinning and Cognitive Impairment in Parkinson’s Disease without Dementia. IEEE/ACM Trans Comput Biol Bioinform 15, 570–580. 10.1109/TCBB.2015.2465951.

54. Polymeropoulos, M.H., Lavedan, C., Leroy, E., Ide, S.E., Dehejia, A., Dutra, A., Pike, B., Root, H., Rubenstein, J., Boyer, R., et al. (1997). Mutation in the alpha-synuclein gene identified in families with Parkinson’s disease. Science 276, 2045–2047. 10.1126/science.276.5321.2045.

55. Conway, K.A., Harper, J.D., and Lansbury, P.T. (1998). Accelerated in vitro fibril formation by a mutant alpha-synuclein linked to early-onset Parkinson disease. Nat Med 4, 1318–1320. 10.1038/3311.

56. Luk, K.C., Song, C., O’Brien, P., Stieber, A., Branch, J.R., Brunden, K.R., Trojanowski, J.Q., and Lee, V.M. (2009). Exogenous alpha-synuclein fibrils seed the formation of Lewy body-like intracellular inclusions in cultured cells. Proc Natl Acad Sci U S A 106, 20051–20056. 10.1073/pnas.0908005106.

57. Pajares, M., Rojo, A.I., Arias, E., Diaz-Carretero, A., Cuervo, A.M., and Cuadrado, A. (2018). Transcription factor NFE2L2/NRF2 modulates chaperone-mediated autophagy through the regulation of LAMP2A. Autophagy 14, 1310–1322. 10.1080/15548627.2018.1474992.

58. Dodson, M., Anandhan, A., Zhang, D.D., and Madhavan, L. (2021). An NRF2 Perspective on Stem Cells and Ageing. Front Aging 2, 690686. 10.3389/fragi.2021.690686.

59. Joshi, C.S., Mora, A., Felder, P.A., and Mysorekar, I.U. (2021). NRF2 promotes urothelial cell response to bacterial infection by regulating reactive oxygen species and RAB27B expression. Cell Rep 37, 109856. 10.1016/j.celrep.2021.109856.

60. Park, J.Y., Kim, S., Sohn, H.Y., Koh, Y.H., and Jo, C. (2019). TFEB activates Nrf2 by repressing its E3 ubiquitin ligase DCAF11 and promoting phosphorylation of p62. Sci Rep 9, 14354. 10.1038/s41598-019-50877-8.

61. Abokyi, S., Shan, S.W., To, C.H., Chan, H.H., and Tse, D.Y. (2020). Autophagy Upregulation by the TFEB Inducer Trehalose Protects against Oxidative Damage and Cell Death Associated with NRF2 Inhibition in Human RPE Cells. Oxid Med Cell Longev 2020, 5296341. 10.1155/2020/5296341.

62. Suzen, S., Tucci, P., Profumo, E., Buttari, B., and Saso, L. (2022). A Pivotal Role of Nrf2 in Neurodegenerative Disorders: A New Way for Therapeutic Strategies. Pharmaceuticals (Basel) 15. 10.3390/ph15060692.

63. Cortes, C.J., and La Spada, A.R. (2019). TFEB dysregulation as a driver of autophagy dysfunction in neurodegenerative disease: Molecular mechanisms, cellular processes, and emerging therapeutic opportunities. Neurobiol Dis 122, 83–93. 10.1016/j.nbd.2018.05.012.

64. Zhang, S., Wang, R., and Wang, G. (2019). Impact of Dopamine Oxidation on Dopaminergic Neurodegeneration. ACS Chem Neurosci 10, 945–953. 10.1021/acschemneuro.8b00454.

65. Meiser, J., Weindl, D., and Hiller, K. (2013). Complexity of dopamine metabolism. Cell Commun Signal 11, 34. 10.1186/1478-811X-11-34.

66. Callizot, N., Combes, M., Henriques, A., and Poindron, P. (2019). Necrosis, apoptosis, necroptosis, three modes of action of dopaminergic neuron neurotoxins. PLoS One 14, e0215277. 10.1371/journal.pone.0215277.

67. Nagakannan, P., Tabeshmehr, P., and Eftekharpour, E. (2020). Oxidative damage of lysosomes in regulated cell death systems: Pathophysiology and pharmacologic interventions. Free Radic Biol Med 157, 94–127. 10.1016/j.freeradbiomed.2020.04.001.

68. Zhu, S.Y., Yao, R.Q., Li, Y.X., Zhao, P.Y., Ren, C., Du, X.H., and Yao, Y.M. (2020). Lysosomal quality control of cell fate: a novel therapeutic target for human diseases. Cell Death Dis 11, 817. 10.1038/s41419-020-03032-5.

69. Huang, J., Pan, W., Ou, D., Dai, W., Lin, Y., Chen, Y., and Chen, X. (2015). LC3B, a Protein That Serves as an Autophagic Marker, Modulates Angiotensin II-induced Myocardial Hypertrophy. J Cardiovasc Pharmacol 66, 576–583. 10.1097/fjc.0000000000000306.

70. Runwal, G., Stamatakou, E., Siddiqi, F.H., Puri, C., Zhu, Y., and Rubinsztein, D.C. (2019). LC3-positive structures are prominent in autophagy-deficient cells. Sci Rep 9, 10147. 10.1038/s41598-019-46657-z.

71. Guerra, F., Girolimetti, G., Beli, R., Mitruccio, M., Pacelli, C., Ferretta, A., Gasparre, G., Cocco, T., and Bucci, C. (2019). Synergistic Effect of Mitochondrial and Lysosomal Dysfunction in Parkinson’s Disease. Cells 8. 10.3390/cells8050452.

72. Fan, Y., Li, C., Peng, X., Jiang, N., Hu, L., Gu, L., Zhu, G., Zhao, G., and Lin, J. (2020). Perillaldehyde Ameliorates Aspergillus fumigatus Keratitis by Activating the Nrf2/HO-1 Signaling Pathway and Inhibiting Dectin-1-Mediated Inflammation. Invest Ophthalmol Vis Sci 61, 51. 10.1167/iovs.61.6.51.

73. Fuyuno, Y., Uchi, H., Yasumatsu, M., Morino-Koga, S., Tanaka, Y., Mitoma, C., and Furue, M. (2018). Perillaldehyde Inhibits AHR Signaling and Activates NRF2 Antioxidant Pathway in Human Keratinocytes. Oxid Med Cell Longev 2018, 9524657. 10.1155/2018/9524657.

74. Tang, L.F., Ma, X., Xie, L.W., Zhou, H., Yu, J., Wang, Z.X., and Li, M. (2023). Perillaldehyde Mitigates Ionizing Radiation-Induced Intestinal Injury by Inhibiting Ferroptosis via the Nrf2 Signaling Pathway. Mol Nutr Food Res, e2300232. 10.1002/mnfr.202300232.

75. Zheng, W., Liu, B., and Shi, E. (2021). Perillaldehyde Alleviates Spinal Cord Ischemia-Reperfusion Injury Via Activating the Nrf2 Pathway. J Surg Res 268, 308–317. 10.1016/j.jss.2021.06.055.

76. Duffy, P.E., and Tennyson, V.M. (1965). Phase and electron microscopic observations of Lewy bodies and melanin granules in the substantia nigra and locus caeruleus in Parkinson’s disease. Journal of Neuropathology & Experimental Neurology 24, 398–414.

77. Forno, L.S., and Norville, R.L. (1976). Ultrastructure of Lewy bodies in the stellate ganglion. Acta neuropathologica 34, 183–197.

78. Watanabe, I., Vachal, E., and Tomita, T. (1977). Dense core vesicles around the Lewy body in incidental Parkinson’s disease: an electron microscopic study. Acta neuropathologica 39, 173–175.

79. Galloway, P., Mulvihill, P., and Perry, G. (1992). Filaments of Lewy bodies contain insoluble cytoskeletal elements. The American journal of pathology 140, 809.

80. Colosimo, C., Hughes, A., Kilford, L., and Lees, A. (2003). Lewy body cortical involvement may not always predict dementia in Parkinson’s disease. Journal of Neurology, Neurosurgery & Psychiatry 74, 852–856.

81. Lam, I., Ndayisaba, A., Lewis, A.J., Fu, Y., Sagredo, G.T., Zaccagnini, L., Sandoe, J., Sanz, R.L., Vahdatshoar, A., Martin, T.D., et al. (2022). Rapid iPSC inclusionopathy models shed light on formation, consequence and molecular subtype of α-synuclein inclusions. BioRxiv.

82. Smith, J.A., Das, A., Ray, S.K., and Banik, N.L. (2012). Role of pro-inflammatory cytokines released from microglia in neurodegenerative diseases. Brain Res Bull 87, 10–20. 10.1016/j.brainresbull.2011.10.004.

83. Mahul-Mellier, A.L., Burtscher, J., Maharjan, N., Weerens, L., Croisier, M., Kuttler, F., Leleu, M., Knott, G.W., and Lashuel, H.A. (2020). The process of Lewy body formation, rather than simply alpha-synuclein fibrillization, is one of the major drivers of neurodegeneration. Proc Natl Acad Sci U S A 117, 4971–4982. 10.1073/pnas.1913904117.

84. Fares, M.B., Jagannath, S., and Lashuel, H.A. (2021). Reverse engineering Lewy bodies: how far have we come and how far can we go? Nat Rev Neurosci 22, 111–131. 10.1038/s41583-020-00416-6.

85. Zhang, X., Wang, R., Chen, H., Jin, C., Jin, Z., Lu, J., Xu, L., Lu, Y., Zhang, J., and Shi, L. (2022). Aged microglia promote peripheral T cell infiltration by reprogramming the microenvironment of neurogenic niches. Immun Ageing 19, 34. 10.1186/s12979-022- 00289-6.

86. Frucht, D.M., Fukao, T., Bogdan, C., Schindler, H., O’Shea, J.J., and Koyasu, S. (2001). IFN-gamma production by antigen-presenting cells: mechanisms emerge. Trends Immunol 22, 556–560. 10.1016/s1471-4906(01)02005-1.

87. Kasen, A., Houck, C., Burmeister, A.R., Sha, Q., Brundin, L., and Brundin, P. (2022). Upregulation of alpha-synuclein following immune activation: Possible trigger of Parkinson’s disease. Neurobiol Dis 166, 105654. 10.1016/j.nbd.2022.105654.

88. Scudamore, O., and Ciossek, T. (2018). Increased Oxidative Stress Exacerbates alpha-Synuclein Aggregation In Vivo. J Neuropathol Exp Neurol 77, 443–453. 10.1093/jnen/nly024.

89. Morell, C., Bort, A., Vara-Ciruelos, D., Ramos-Torres, A., Altamirano-Dimas, M., Diaz-Laviada, I., and Rodriguez-Henche, N. (2016). Up-Regulated Expression of LAMP2 and Autophagy Activity during Neuroendocrine Differentiation of Prostate Cancer LNCaP Cells. PLoS One 11, e0162977. 10.1371/journal.pone.0162977.

90. Martinez-Vicente, M., Talloczy, Z., Kaushik, S., Massey, A.C., Mazzulli, J., Mosharov, E.V., Hodara, R., Fredenburg, R., Wu, D.C., Follenzi, A., et al. (2008). Dopamine-modified alpha-synuclein blocks chaperone-mediated autophagy. J Clin Invest 118, 777–788. 10.1172/JCI32806.

91. Malkus, K.A., and Ischiropoulos, H. (2012). Regional deficiencies in chaperone-mediated autophagy underlie alpha-synuclein aggregation and neurodegeneration. Neurobiol Dis 46, 732–744. 10.1016/j.nbd.2012.03.017.

92. Saha, S., Buttari, B., Profumo, E., Tucci, P., and Saso, L. (2021). A Perspective on Nrf2 Signaling Pathway for Neuroinflammation: A Potential Therapeutic Target in Alzheimer’s and Parkinson’s Diseases. Front Cell Neurosci 15, 787258. 10.3389/fncel.2021.787258.

93. Tanudjojo, B., Shaikh, S.S., Fenyi, A., Bousset, L., Agarwal, D., Marsh, J., Zois, C., Heman-Ackah, S., Fischer, R., Sims, D., et al. (2021). Phenotypic manifestation of alpha-synuclein strains derived from Parkinson’s disease and multiple system atrophy in human dopaminergic neurons. Nat Commun 12, 3817. 10.1038/s41467-021-23682-z.

94. Iannielli, A., Luoni, M., Giannelli, S.G., Ferese, R., Ordazzo, G., Fossati, M., Raimondi, A., Opazo, F., Corti, O., Prehn, J.H.M., et al. (2022). Modeling native and seeded Synuclein aggregation and related cellular dysfunctions in dopaminergic neurons derived by a new set of isogenic iPSC lines with SNCA multiplications. Cell Death Dis 13, 881. 10.1038/s41419-022-05330-6.

95. Gribaudo, S., Tixador, P., Bousset, L., Fenyi, A., Lino, P., Melki, R., Peyrin, J.M., and Perrier, A.L. (2019). Propagation of alpha-Synuclein Strains within Human Reconstructed Neuronal Network. Stem Cell Reports 12, 230–244. 10.1016/j.stemcr.2018.12.007.

96. Mohamed, N.V., Sirois, J., Ramamurthy, J., Mathur, M., Lepine, P., Deneault, E., Maussion, G., Nicouleau, M., Chen, C.X., Abdian, N., et al. (2021). Midbrain organoids with an SNCA gene triplication model key features of synucleinopathy. Brain Commun 3, fcab223. 10.1093/braincomms/fcab223.

97. Miura, Y., Li, M.Y., Revah, O., Yoon, S.J., Narazaki, G., and Pasca, S.P. (2022). Engineering brain assembloids to interrogate human neural circuits. Nat Protoc 17, 15–35. 10.1038/s41596-021-00632-z.

98. Birey, F., Andersen, J., Makinson, C.D., Islam, S., Wei, W., Huber, N., Fan, H.C., Metzler, K.R.C., Panagiotakos, G., Thom, N., et al. (2017). Assembly of functionally integrated human forebrain spheroids. Nature 545, 54–59. 10.1038/nature22330.

99. Castellanos-Montiel, M.J., Chaineau, M., Franco-Flores, A.K., Haghi, G., Carrillo-Valenzuela, D., Reintsch, W.E., Chen, C.X., and Durcan, T.M. (2023). An Optimized Workflow to Generate and Characterize iPSC-Derived Motor Neuron (MN) Spheroids. Cells 12. 10.3390/cells12040545.

100. Chen, C.X.-Q., Abdian, N., Maussion, G., Thomas, R.A., Demirova, I., Cai, E., Tabatabaei, M., Beitel, L.K., Karamchandani, J., Fon, E.A., and Durcan, T.M. (2021). Standardized quality control workflow to evaluate the reproducibility and differentiation potential of human iPSCs into neurons. bioRxiv, 2021.2001.2013.426620. 10.1101/2021.01.13.426620.

101. Maneca, D.-L., Luo, W., Krahn, A., Del Cid Pellitero, E., Shlaifer, I., Beitel, L.K., Rao, T., and Durcan, T.M. (2019). Production of Recombinant α Synuclein Monomers and Preformed Fibrils (PFFs). Zenodo. 10.5281/zenodo.3738335.

102. Wen, W., Zhang, J.P., Xu, J., Su, R.J., Neises, A., Ji, G.Z., Yuan, W., Cheng, T., and Zhang, X.B. (2016). Enhanced Generation of Integration-free iPSCs from Human Adult Peripheral Blood Mononuclear Cells with an Optimal Combination of Episomal Vectors. Stem Cell Reports 6, 873–884. 10.1016/j.stemcr.2016.04.005.

103. Del Cid Pellitero, E., Shlaifer, R., Luo, W., Krahn, A., Nguyen-Renou, E., Manecka, D.-L., Rao, T., Beitel, L., and Durcan, T.M. (2019). Quality Control Characterization of α-Synuclein Preformed Fibrils (PFFs). Zenodo. 10.5281/zenodo.3738340.

104. Feller, B., Fallon, A., Luo, W., Nguyen, P.T., Shlaifer, I., Lee, A.K., Chofflet, N., Yi, N., Khaled, H., Karkout, S., et al. (2023). alpha-Synuclein Preformed Fibrils Bind to beta-Neurexins and Impair beta-Neurexin-Mediated Presynaptic Organization. Cells 12. 10.3390/cells12071083.

105. Bayati, A., Luo, W., Del Cid-Pellitero, E., Fon, E.A., Durcan, T.M., and McPherson, P.S. (2023). Visualization of alpha-synuclein trafficking via nanogold labeling and electron microscopy. STAR Protoc 4, 102113. 10.1016/j.xpro.2023.102113.

106. Jefri, M., Bell, S., Peng, H., Hettige, N., Maussion, G., Soubannier, V., Wu, H., Silveira, H., Theroux, J.F., Moquin, L., et al. (2020). Stimulation of L-type calcium channels increases tyrosine hydroxylase and dopamine in ventral midbrain cells induced from somatic cells. Stem Cells Transl Med 9, 697–712. 10.1002/sctm.18-0180.

107. Chen, X.R., Cecilia; Loignon, Martin; Peng, Huasheng; Rao, Trisha; Durcan, Thomas Martin (2019). Induction of Dopaminergic or Cortical neuronal progenitors from iPSCs. Zenodo. 10.5281/zenodo.3364831.

108. Sheta, R., Teixeira, M., Idi, W., Pierre, M., de Rus Jacquet, A., Emond, V., Zorca, C.E., Vanderperre, B., Durcan, T.M., Fon, E.A., et al. (2022). Combining NGN2 programming and dopaminergic patterning for a rapid and efficient generation of hiPSC-derived midbrain neurons. Sci Rep 12, 17176. 10.1038/s41598-022-22158-4.

109. Zhang, Y., Pak, C., Han, Y., Ahlenius, H., Zhang, Z., Chanda, S., Marro, S., Patzke, C., Acuna, C., Covy, J., et al. (2013). Rapid single-step induction of functional neurons from human pluripotent stem cells. Neuron 78, 785–798. 10.1016/j.neuron.2013.05.029.

110. Meijer, M., Rehbach, K., Brunner, J.W., Classen, J.A., Lammertse, H.C.A., van Linge, L.A., Schut, D., Krutenko, T., Hebisch, M., Cornelisse, L.N., et al. (2019). A Single-Cell Model for Synaptic Transmission and Plasticity in Human iPSC-Derived Neurons. Cell Rep 27, 2199–2211 e2196. 10.1016/j.celrep.2019.04.058.

111. Chen, Y., Dolt, K.S., Kriek, M., Baker, T., Downey, P., Drummond, N.J., Canham, M.A., Natalwala, A., Rosser, S., and Kunath, T. (2019). Engineering synucleinopathy-resistant human dopaminergic neurons by CRISPR-mediated deletion of the SNCA gene. Eur J Neurosci 49, 510–524. 10.1111/ejn.14286.

112. Tong, L., Balazs, R., Soiampornkul, R., Thangnipon, W., and Cotman, C.W. (2008). Interleukin-1 beta impairs brain derived neurotrophic factor-induced signal transduction. Neurobiol Aging 29, 1380–1393. 10.1016/j.neurobiolaging.2007.02.027.

113. Labno, C. Two Ways to Count Cells with ImageJ. https://www.unige.ch/medecine/bioimaging/files/3714/1208/5964/CellCounting.pdf.

114. Legland, D., Arganda-Carreras, I., and Andrey, P. (2016). MorphoLibJ: integrated library and plugins for mathematical morphology with ImageJ. Bioinformatics 32, 3532–3534. 10.1093/bioinformatics/btw413.

115. Peters, A.E., Caban, S.J., McLaughlin, E.A., Roman, S.D., Bromfield, E.G., Nixon, B., and Sutherland, J.M. (2021). The Impact of Aging on Macroautophagy in the Pre-ovulatory Mouse Oocyte. Front Cell Dev Biol 9, 691826. 10.3389/fcell.2021.691826.

